# Single-molecule analysis reveals the mechanism of chromatin ubiquitylation by variant PRC1 complexes

**DOI:** 10.1101/2024.10.25.620026

**Authors:** Alexandra Teslenko, Beat Fierz

## Abstract

Chromatin regulation relies on ‘writer’ enzymes that add post-translational modifications (PTMs) to histone proteins. Variant Polycomb repressive complex 1 (PRC1) exists as several subtypes, which are key ‘writers’ of ubiquitylation on histone H2A lysine 119 (H2AK119ub), crucial for transcriptional repression during development and cell identity determination. The mechanism by which dynamic chromatin exploration by varia nt PRC1 complexes is coupled to ubiquitin writing is unknown. Here, we developed a single-molecule approach to directly observe chromatin interactions and ubiquitylation by PRC1. We find that variant PRC1 transiently samples chromatin until reaching a catalytically competent nucleosome-bound state, resulting in E2 recruitment and ubiquitin transfer. Variant PRC1 is weakly processive in ubiquitylating neighboring nucleosomes. Moreover, activity differences between PRC1 subtypes, containing either a PCGF1 or PCGF4 subunit, result from distinct probabilities of achieving a catalytically competent state. Our results thus demonstrate that the dynamic formation of an active complex between variant PRC1, E2 and chromatin is the critical determinant of subtype-specific variant PRC1 activity.

## Introduction

In eukaryotes, genomic DNA is compacted into chromatin (*1*), whose higher-order structure controls the accessibility of genetic information (*2*). Post-translational modifications (PTMs) installed on histone proteins form an epigenetic code, regulating chromatin organization and gene transcription, as well as other processes such as DNA replication or repair (*3*). Chromatin modifying enzymes, including ‘writers’ that site-specifically install PTMs, and ‘erasers’ that remove PTMs, often contain ‘reader’-domains that recognize and bind to specific PTMs, and together maintain this epigenetic landscape (*4–6*).

An important family of such regulatory factors are the Polycomb group (PcG) proteins, which form an evolutionarily conserved system for gene repression. PcG proteins include the chromatin modifying enzymes Polycomb repressive complex 1 (PRC1) and 2 (PRC2) (*7–9*). PRC1 complexes are E3 ubiquitin ligases that monoubiquitylate histone H2A at lysine 119 (H2AK119ub) (*10*). This PTM is then recognized by PRC2 (*11–13*), which is a methyltransferase specific for H3K27 (*14–17*), creating further ‘landing’ sites for PRC1. Together, PRC1 and 2 induce gene repression at target loci, crucial for cellular differentiation and embryonic development (*8*, *18*), by keeping promoters in an off-state (*19*). PRC1 complexes contain the RING protein RING1A/B, together with one of six PCGF proteins (PCGF1-6), forming heterodimeric E3 ubiquitin ligases (*10*, *20–22*). The presence of a given PCGF subunit, together with additional auxiliary proteins within the complex, allow further classification of PRC1 complexes into ‘canonical’ and ‘non-canonical’, or ‘variant’, PRC1. Canonical PRC1 generally contain PCGF2 or 4 and a chromobox (CBX) protein, whereas variant PRC1 complexes can contain each of the six PCGF subunits and are associated with RING1/YY1-binding protein (RYBP) or its paralogue YY1-associated factor 2 (YAF2) (*21*). These subunits are important for PRC1 targeting, with CBX proteins in canonical PRC1 binding H3K27me2/3 via the CBX domain(*23*), and RYBP in variant PRC1 binding H2AK119ub via its N-terminus (*24–27*). While canonical PRC1 is directly involved in gene silencing and chromatin compaction, variant PRC1 complexes function at the top of the cascade, installing H2AK119ub across the genome (*28*, *29*). Moreover, variant PRC1 plays a key role in controlling embryonic stem cell proliferation (*30*), repression of retroviruses (*31*), regulation of metabolic genes and progression of the cell cycle (*32*).

To ubiquitylate chromatin, PRC1 complexes engage the surface of the nucleosome in a defined orientation, contacting the nucleosomal acidic patch via nucleosome-binding loops within their RING domains (*33*). This orients the E2 ubiquitin-conjugating enzyme towards H2AK119 and positions the ubiquitin thioester in a catalytically competent position. Specifically, the RING1-PCGF heterodimer stabilizes the flexible conformation of the E2 that carries the activated ubiquitin moiety (E2∼Ub). In this conformation the ubiquitin C-terminus is extended and the thioester bond is oriented for the nucleophilic attack at H2AK119 (*34*, *35*). Therefore, the steps of nucleosome binding (i.e. chromatin state reading), establishment of a catalytically competent conformation of the bound E3 ligase, E2 recruitment, and catalysis of ubiquitin discharge are all potentially rate limiting and determine the spatiotemporal activity of PRC1 complexes, which varies greatly between different PRC1 subtypes. In particular, the nature of the incorporated PCGF proteins (*21*, *36*, *37*) determines both E2 discharge rates in the absence of substrate or on nucleosomes, with PCGF1 and 5 being more active, whereas PCGF4 shows overall lower activity (*38*, *39*). The molecular origins of these different activities on chromatin substrates are however still not clear. Moreover, auxiliary subunits play a key role in controlling the activity of the complex. RYBP, in particular, stimulates chromatin ubiquitylation by promoting nucleosomal interactions (*21*, *39*, *40*). A recent cryo-EM structure further demonstrated how RYBP functions in anchoring variant PRC1 on nucleosomes containing H2AK119ub, as a potential mechanism in spreading H2AK119ub (*27*). However, due to the absence of dynamic data of the whole catalytic process, we do not know how variant PRC1 searches and dynamically interacts with chromatin, how different PCGF subtypes differ in chromatin engagement, and how catalytic activity is determined. In fact, such information is generally unavailable for most chromatin readers and writers but is critically important for a better understanding of chromatin regulation.

Here, we develop a fluorescence-based method to directly observe the chromatin reading and writing activity of variant PRC1 in real time on the single-molecule level, building on our developments in observing chromatin interaction dynamics of chromatin regulatory proteins (*41–43*) including PRC2 (*44*), and expanding on methods in monitoring ubiquitylation reactions (*45*). We find that variant PRC1 complexes dynamically probe chromatin until they bind nucleosomes in a catalytically competent state. Transient recruitment of E2∼Ub leads to H2A ubiquitylation followed by PRC1 release. We observe weak processivity, with most complexes depositing either one or two ubiquitin moieties during a single chromatin engagement event. Moreover, E2∼Ub recruitment and E2-E3 complex positioning determine overall activities between variant PRC1 subtypes. This results in faster global ubiquitylation rates for PCGF1-based PRC1 complexes (PRC1.1), compared to PCGF4-containing enzymes (PRC1.4). Finally, we build a mechanistic model of chromatin ‘reading’ and ‘writing’ by variant PRC1 based on the measured binding and enzymatic reaction kinetics. Together, our approach provides a dynamic single-molecule view into the mechanisms of function of chromatin modifiers, and our findings demonstrate that the dynamic assembly of an active complex is crucial in defining the PCGF-specific activity of PRC1.

## Results

Histone-modifying enzymes have to efficiently sample the vast chromatin landscape to establish and maintain specific epigenetic chromatin states. Previous *in vitro* and cellular studies of the dynamics of PcG family proteins revealed that the methyltransferase PRC2 only transiently interacts with chromatin fibers with residence times in the seconds time regime (*44*, *46*). Similarly, single-molecule tracking of variant PRC1 in living cells revealed rapid diffusion of the complexes throughout the nucleus, with an overall low occupancy at target sites (*47*). These findings therefore raise the question of how efficient chromatin modification can be established by such dynamic enzymes, and how binding and enzymatic activity are coupled.

### Development of a multicolor single-molecule method to study PRC1

Here, we tackled this question for variant PRC1, building on methodology established in our laboratory (*44*, *48*). We envisioned an assay revealing chromatin binding and subsequent ubiquitin transfer of variant PRC1 in real time by multicolor single-molecule total internal reflection fluorescence microscopy (smTIRFM) (**Fig. 1A**). In such an assay, fluorescently labeled chromatin fibers are immobilized in a flow cell and their position is determined. Upon addition of variant PRC1, tagged in a different color, colocalization of the fluorescence signals indicates transient PRC1 binding to the immobilized chromatin fibers (‘reading’). In the presence of E2 and ubiquitin, carrying a third spectrally separated dye, chromatin ubiquitylation is followed in real time (‘writing’). The resulting single-molecule fluorescence time traces then yield mechanistic insights into variant PRC1 function, in particular on the coupling between chromatin reading and writing and on the processivity of PRC1 activity within a chromatin environment. Moreover, we expected that single-molecule dynamics of different variant PRC1 subtypes would allow us to reveal the kinetic determinants controlling their distinct activities.

**Fig. 1.**
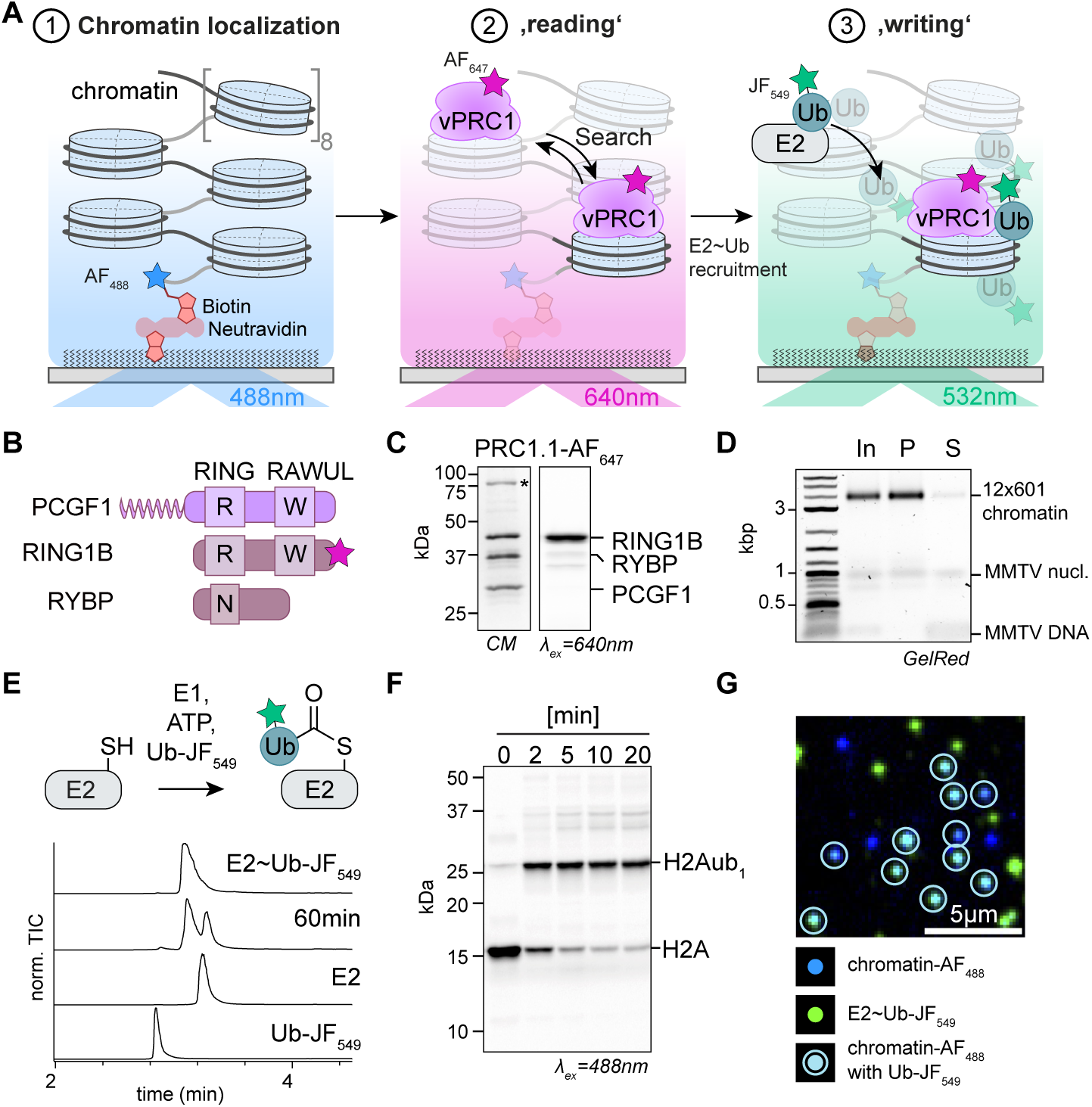
| Establishment of an in-situ chromatin ubiquitylation assay. **A)** Scheme of the three-color single-molecule experiment to observe chromatin ubiquitylation, with chromatin fibers detected via Alexa Fluor 488 (AF_488_) emission in the blue-green channel, variant PRC1 binding dynamics observed via Alexa Fluor 647 (AF_647_) emission in the red-far red channel, and chromatin ubiquitylation monitored by Janelia Fluor 549 (JF_549_) emission in the green-orange channel. **B**) PRC1.1 subunit composition. The pink star indicates the position of the AF_647_ dye (see also **Fig. S1**). **C**) SDS-PAGE of purified, fluorescently labeled PRC1.1. CM: Coomassie blue, λ_ex_ = 640 nm: AF_647_ fluorescence after 640 nm irradiation. Asterisk: Hsp70 associated with PRC1.1. **D**) Agarose gel electrophoretic analysis of magnesium precipitated 12×601 chromatin fibers. In: Input, P: pellet containing pure chromatin fibers, Sup: supernatant containing buffer DNA and ‘buffer’ nucleosomes on mouse mammary tumor virus (MMTV) DNA (**Fig. S2**). **E**) Loading of E2 (UbcH5c) with fluorescently labeled ubiquitin. Top: Reaction scheme, Bottom: Analysis of the reaction progress via LC-ESI-MS. see also **Fig. S3 and S4A-E**) **F**) SDS-PAGE analysis of ubiquitylation assay on chromatin fibers containing H2A labeled with Atto 488 dye (H2A-Atto_488_) with E2∼Ub and PRC1. λ_ex_ = 488 nm: fluorescence emission from H2A-Atto_488_. (see also **Fig. S4F**) **G**) smTIRF analysis of ubiquitylation products, showing chromatin fibers in blue, E2∼Ub and Ub in green. Cyan spots show ubiquitylated single chromatin fibers. For additional views, see **Fig. S4G**.

To develop the methodology to image chromatin ubiquitylation in real time, we recombinantly expressed and purified human PRC1.1 complex, containing the subunits RING1B, RYBP and PCGF1, from insect cells (**Fig. 1B**). During purification, we labeled the RING1B subunit with a Alexa Fluor 647 (AF_647_) fluorophore at the C-terminus via a short peptide tag (*49*) with an efficiency above 80% (**Fig. 1C** and **S1A-G**), allowing its detection in the far-red channel (**Fig. 1A**). We then performed ubiquitylation assays using the ubiquitin-conjugating enzyme E2 (UbcH5c), demonstrating that the introduction of the fluorescent label did not affect the activity of PRC1.1 (**Fig. S1H-I**). In parallel, we assembled chromatin fibers, using recombinant human core histones as well as a DNA template containing 12 copies of the Widom 601 nucleosome positioning sequence (*50*, *51*), separated by 30 base pairs (bp) of linker DNA (12-mer chromatin fiber), tagged with Alexa Fluor 488 (AF_488_) dye for single-molecule detection and biotin for immobilization (**Fig. 1Dl S2** and **S3A-F**). Finally, to observe chromatin ubiquitylation, we tagged ubiquitin at the N-terminus with Janelia Fluor 549 (JF_549_) (*52*) (**Fig. S3G-K**). Micromolar concentrations of fluorescently labeled ubiquitin will result in high background and render single-molecule detections via smTIRF impossible. We thus pre-loaded UbcH5c enzymatically with fluorescently labeled ubiquitin and purified the charged E2 (E2∼Ub), removing all unbound ubiquitin (**Fig. 1E, S3** and **S4**). We then probed the activity of E2∼Ub on chromatin fibers containing fluorescently labeled histone H2A (**Fig. S5**), revealing rapid chromatin modification as expected (**Fig. 1F** and **S4F**). Together, using pre-loaded E2∼Ub allowed us to drastically lower the concentration of labeled species in our assays.

Finally, to test our ability to detect ubiquitylation of single nucleosomes using single-molecule fluorescence, we immobilized chromatin fibers in a flow cell and exposed them to PRC1.1 and preloaded E2∼Ub. After a washing step, we observed the colocalization of chromatin fibers and ubiquitin via smTIRF (**Fig. 1G** and **S4G**). Together, this demonstrated that the enzymes are active and that variant PRC1-dependent chromatin ubiquitylation can be directly observed using preloaded E2∼Ub on the single-molecule level.

### Real-time detection of chromatin ubiquitylation by PRC1.1

We thus proceeded to image chromatin ubiquitylation dynamics in real time. After immobilizing chromatin fibers in flow channels, we injected fluorescently labeled PRC1.1 and E2∼Ub, while imaging using a 3-color excitation protocol (**Fig. S6A-B**). This allowed us to detect transient PRC1.1 binding events to the 12-mer chromatin fibers, whose positions were previously determined (**Fig. 2A** left panel), as colocalizing fluorescence emission spots in the far-red channel (now referred to as the ‘PRC1 channel’, **Fig. 2A**, middle panel). Importantly, for these experiments we chose a concentration of 0.5 nM for PRC1.1, which allowed us to mostly observe individual binding events on single chromatin fibers that did not overlap in time. Moreover, under the experimental conditions (5 mM MgCl_2_), chromatin fibers were in a compact state (*53*), which however remains highly dynamic and accessible (*48*). Extraction of fluorescence intensity time traces from single chromatin fiber positions allowed us to directly observe the dynamics of PRC1.1 chromatin binding in the PRC1-channel (**Fig. 2B-C**). PRC1.1 exhibited transient chromatin interactions, lasting between hundreds of milliseconds to several seconds.

**Fig. 2.**
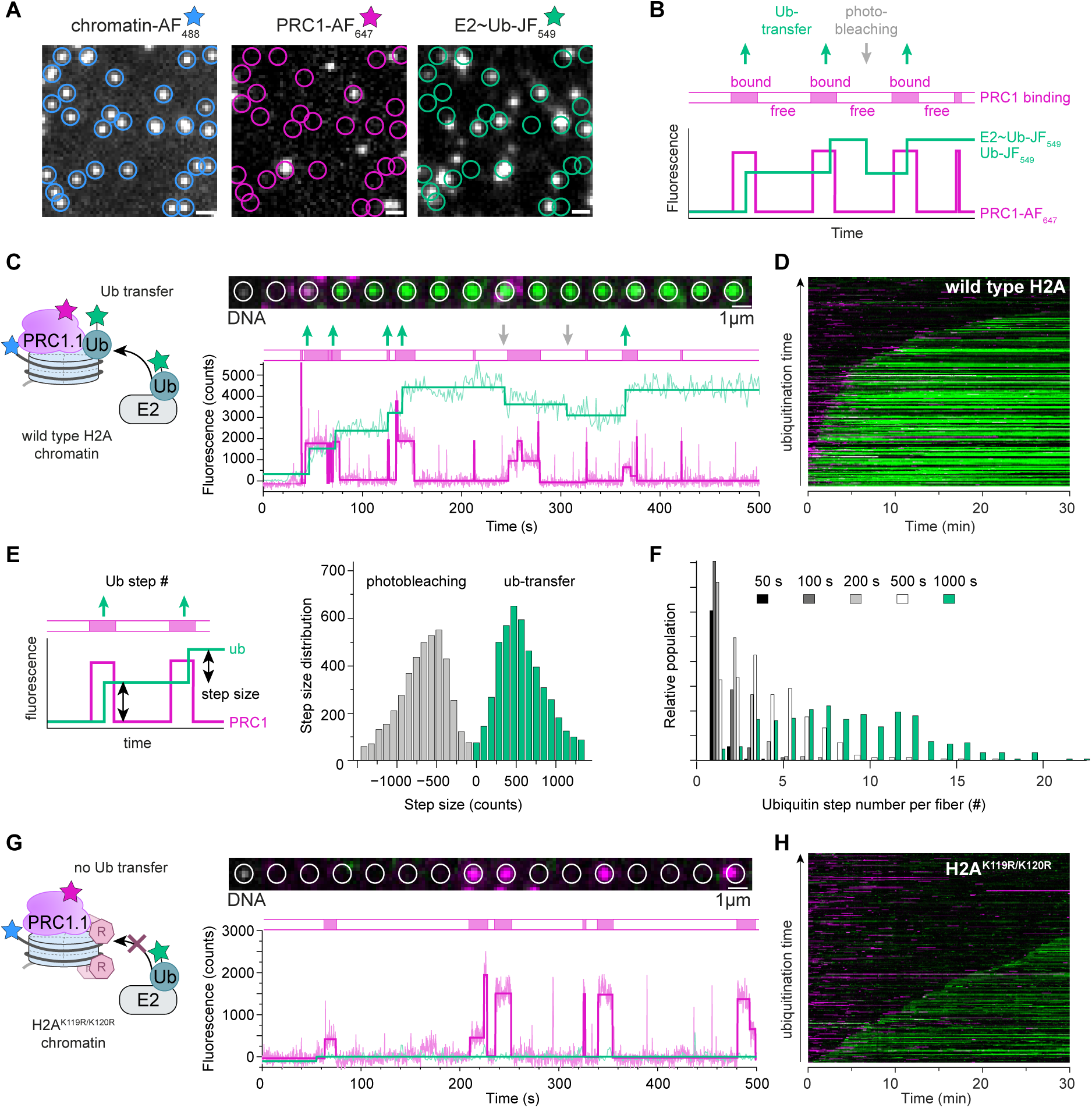
| Single-molecule observation of dynamic chromatin ubiquitylation by PRC1.1. **A**) Single movie frames from chromatin ubiquitylation experiment (for experimental scheme, see **Fig. 1A**). Left: chromatin channel, 488 nm excitation, 498-523 nm emission, middle: PRC1 channel, 640 nm excitation, 659 - 754 nm emission, right: ubiquitin channel, 532 nm excitation, 553 - 618 nm emission. **B**) Scheme of single-molecule fluorescence time trace. Magenta: PRC1-channel, showing PRC1-AF_647_ chromatin binding. Green: ubiquitin channel, showing chromatin ubiquitylation with Ub-JF_549_. Green arrows: E2∼Ub binding / ubiquitin transfer events. Grey arrows: Photobleaching events of JF_549_ dye (see also **Fig. S6F**). **C**) PRC1.1 chromatin ubiquitylation experiment observed by single-molecule fluorescence. Left: Reaction scheme. Top: fluorescence images from a single chromatin fiber (white circle: position of chromatin fiber, green: ubiquitin channel, magenta: PRC1 channel). Bottom: Fluorescence time trace, see **B**) for details, and see **Fig. S7A,C** for additional traces and images. **D**) Composite plots of PRC1.1 ubiquitylation process shown in **D**) over 30 min at [E2∼Ub] = 25 nM. Each line represents a time trace, fluorescence intensity is encoded in color intensity. n = 221 traces. See **Fig. S7E** for further plots. **E**) Analysis of step size in the ubiquitin channel. Left: Scheme and definitions. Right: Step size histogram of ubiquitylation and photobleaching events in the ubiquitin channel for the experiment shown in **C-D**), revealing an average step size of 500 counts. **F**) Histogram of the number of ubiquitylation events per chromatin fiber during the indicated time intervals. **G**) Ubiquitylation of chromatin fibers containing mutant H2A^K119R/K120R^ by PRC1.1. Left: Reaction scheme. Top: fluorescence images from a single chromatin fiber. Bottom: fluorescence time trace, refer to **B-C**) for details. See **Fig. S7B,D** for additional representative traces and images. **H**) Composite plots of PRC1.1 ubiquitylation process shown in **G**) over 30 min at [E2∼Ub] = 25 nM. n = 221 traces. See **Fig. S7F** for further plots.

To initiate the ubiquitylation reaction, we injected 25 nM of fluorescently labeled E2∼Ub into the flow channel. A major challenge when performing smTIRF imaging with fluorescently labeled proteins at high concentrations is their propensity to nonspecifically stick to the surface, resulting in overwhelming background fluorescence. We thus exhaustively optimized both surface passivation protocols and imaging procedures (**Fig. S6C-F** and **Methods**), until we reached conditions that provided a suitable signal-to-noise ratio to observe individual ubiquitylation events. Under these optimized conditions, we detected that a subset of transient PRC1.1 binding interactions was associated with the emergence of fluorescence detection in the green-orange channel (now referred to as the ‘ubiquitin channel’, **Fig. 2A**, right panel). We detected discrete steps in fluorescence intensity in the ubiquitin-channel (**Fig. 2B-C**), indicating E2∼Ub recruitment and ubiquitin transfer. The extended integration time of 1.05 s per frame in the ubiquitin channel, necessary to differentiate bound from freely diffusing ubiquitin molecules, prevented us from observing transient recruitment events of E2∼Ub lasting less than one second. However, the formation of a covalent bond between the ubiquitin moiety and H2AK119 within chromatin resulted in a stable increase in fluorescence emission (**Fig. 2C** and **S7C,D** for additional representative traces). Intriguingly, PRC1.1 and ubiquitin signals rarely appeared at the exact same moment but usually with a certain delay, indicating that PRC1.1 binds chromatin first, followed by E2∼Ub recruitment and ubiquitin transfer in a second step.

The full microscopic field of view contained hundreds of immobilized chromatin fibers. To obtain a global view of PRC1.1 binding and chromatin ubiquitylation dynamics, we generated composite plots, where each line represents the fluorescence emission gathered from a single chromatin locus, sorted by the first appearance of a stable signal in the ubiquitin-channel (**Fig. 2D**). This view reveals that for most chromatin loci, ubiquitylation appears within minutes, preceded by several non-productive PRC1.1 binding events. Moreover, for a small subset of chromatin loci, no stable ubiquitylation reaction was detected during the 30-minute observation time (**Fig. 2D**).

After the first ubiquitin transfer event, we generally observed further discrete steps in the fluorescence emission traces in the ubiquitin-channel (**Fig. 2C**) due to the attachment of additional ubiquitin moieties, resulting in progressive accumulation of H2AK119ub within the chromatin fibers. Analyzing step sizes across a whole experiment, we observed both positive and negative steps of similar height (**Fig. 2E**), corresponding to the addition or loss of a single JF_549_ fluorophore. As our reaction mixture did not contain any deubiquitination enzymes, we attributed downward steps to the photobleaching of JF_549_ dyes (**Fig. S6F**). We were further able to quantify the number of transfers per chromatin fiber by counting the upward steps in the ubiquitin channel across various time intervals (**Fig. 2F**). Within the reaction time of 1000 seconds, we observed on average about ten reactions per 12-mer chromatin fiber, corresponding to around 0.8 ubiquitin moieties per nucleosome.

To ensure that the observed stepwise increase in fluorescence emission indeed reflected ubiquitylation at H2AK119, we assembled chromatin containing H2A with lysines K119 and K120 mutated to arginine (H2A^K119R/K120R^) (*54*). Ensemble assays demonstrated that such chromatin was a poor substrate for PRC1-mediated ubiquitylation (**Fig. S8A,B**). Indeed, we did not observe pronounced ubiquitylation over extended time under our single-molecule conditions, showing mostly non-productive PRC1.1 binding events without accompanying signal in the ubiquitin-channel (**Fig. 2G**). Analyzing an extended set of traces via a composite plot (**Fig. 2H**) revealed weak ubiquitylation occurring at later times in the experiment. This indicates that PRC1.1 exhibits non-specific ubiquitylation activity when the preferred substrate lysines are not present, reacting with further available lysines within the nucleosomes, a common behavior for many E3 ligases (*54*, *55*). Importantly, in the absence of any E3, preloaded E2∼Ub by itself did not result in any marked activity on chromatin (**Fig. S8C,D**).

### PRC1.1 exhibits dynamic chromatin recognition stabilized by E2

Having validated the experimental approach, we proceeded to quantitatively analyze PRC1.1 binding and ubiquitylation kinetics to unravel how chromatin reading affects the PTM writing process. First, we explored how PRC1.1 searches chromatin, and specifically binds to the nucleosome surface in a catalytically competent conformation (**Fig. 3A**). We performed smTIRF analysis of PRC1.1 binding reactions in the absence of E2∼Ub (**Fig. 3B**) and generated time traces of binding events for individual chromatin fibers (**Fig. 3C,D**, and **S9A-C** for more traces). From these traces, we determined the length of bright binding events (t_bright_), and of dark states between binding events (t_dark_).

**Fig. 3.**
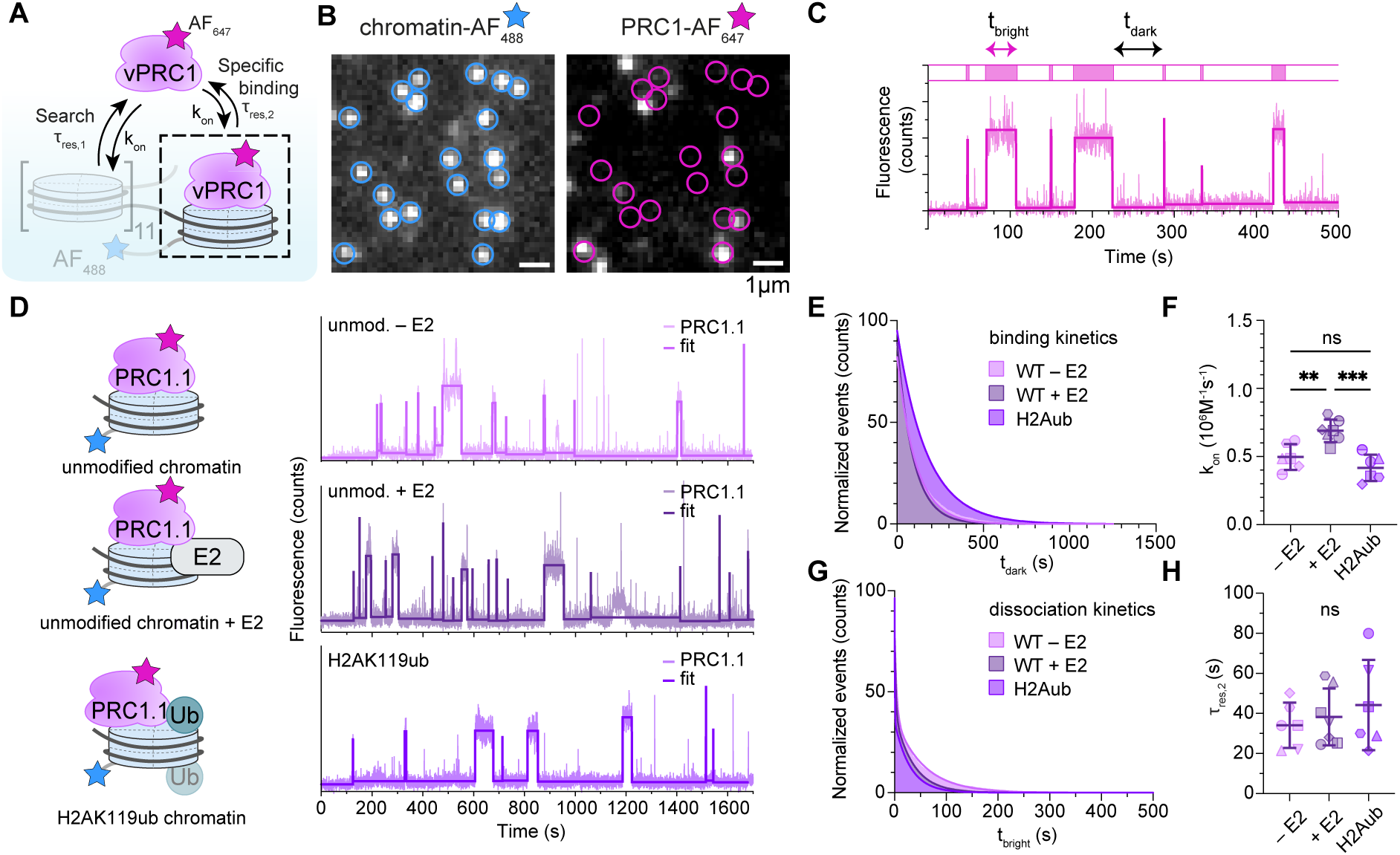
| PRC1.1 dynamically binds chromatin. **A**) Scheme of variant PRC1 nucleosome binding. **B**) Single movie frames from chromatin PRC1.1 chromatin binding experiment. Left: chromatin channel, right: PRC1 channel. **C**) Binding trace for PRC1.1 on chromatin fibers. t_bright_: time intervals for bound PRC1.1; t_dark_: search time intervals between binding events. **D**) Binding dynamics of PRC1.1 to unmodified chromatin fibers (unmod.), fibers containing H2AK119ub, in the absence (-E2) or presence of E2 (+E2, 50 nM), fitted by a step function. Time intervals t_dark_ and t_bright_ are determined via thresholding. See **Fig. S9A-C** for additional binding traces. **E**) Binding kinetics: cumulative histogram of t_dark_ times, solid lines: fit to mono-exponential function (see also **Fig. S9D**). **F**) k_on_ for PRC1.1 binding to different chromatin fibers (see also **Fig. S15F**). n = 6 (unmod. - E2), n = 7 (unmod. + E2), n = 6 (H2AK119ub). Error bars, s.d.; Statistical testing: one-way ANOVA, Tukey’s multiple comparison test (*) p < 0.05, (**) p < 0.01, (***) p < 0.001, ns: non-significant. **G**) Dissociation kinetics: cumulative histogram of t_bright_ times, solid lines: fit to bi-exponential function, see also **Fig. S9E**. **H**) Long residence times ***τ***_res,2_ for PRC1.1 on different chromatin fibers (see also **Methods** and **Fig. S9I**). For a description of statistical analysis see panel **F**). For all data see **Table S1**.

We then constructed lifetime histograms for PRC1.1 chromatin association kinetics using the obtained t_dark_ values (**Fig. 3E** and **S9D**), which revealed that PRC1.1 searches and binds chromatin with an association rate constant (k_on_) of 5.0 ± 0.9 x 10^5^ M^-1^ s^-1^ (**Fig. 3F** and **Table S1**). Conversely, residence time histograms based on t_bright_ values revealed that PRC1.1 dissociates via a bi-exponential process (**Fig. 3G,H** and **S9E**), exhibiting a fast initial decay with a time constant ***τ***_res,1_ = 1.9 ± 0.6 s and a second slower residence time ***τ***_res,2_ = 34.1 ± 10.4 s. This behavior is similar to other chromatin binding proteins, for example PRC2 (*44*). In the presence of E2 (without preloaded ubiquitin), PRC1.1 binding to chromatin was increased as previously reported(*33*). We could largely attribute this effect to an increase in k_on_ (**Fig. 3E,F**) whereas residence times were not significantly influenced in the presence of E2 (**Fig. 3G,H** and **S9F-I**).

Secondly, we wondered if H2A ubiquitylation may increase the residence time of PRC1.1, as direct interactions between the RYBP subunit and H2AK119ub have been reported (*11*, *25*, *27*, *40*). We thus prepared chromatin fibers containing H2AK119ub, which was produced using a click-chemistry-directed chemical ubiquitylation protocol(*56*, *57*) (**Fig. 3D, S10 and 11A-F**). However, we did not any changes in k_on_, and while ***τ***_res_ increased slightly for PRC1.1 binding to these ubiquitylated chromatin substrates, the changes were not significant (**Fig. 3E-H**, and **S9F-I**). This indicates that RYBP-mediated H2AK119ub binding does not play a major role in the chromatin recruitment process under our measurement conditions. Still, RYBP contributed to variant PRC1 activity, as complexes lacking this subunit were less active on chromatin fiber substrates (**Fig. S11G,H**), in line with recent reports (*40*).

### PRC1.1 is a weakly processive enzyme

The direct observation of the PRC1.1 chromatin binding process revealed two distinct interaction modes with either a short or long lifetime (***τ***_res,1_ and ***τ***_res,2_). We previously hypothesized that the fast process (***τ***_res,1_) reflected non-specific chromatin interactions and the slow process (***τ***_res,2_) reported on specific, catalytically competent enzyme-chromatin interactions (*44*). To test this hypothesis, we proceeded to investigate the relationship between enzyme binding and its activity using the single-molecule chromatin ubiquitylation data (**Fig. 2**). We thus generated libraries of individual PRC1.1 binding events that showed associated catalytic activity (i.e. steps in the ubiquitin-channel, or ‘active’’ binding events, **Fig. 4A-C** and **S12A**) from the recorded single-molecule traces of PRC1.1-mediated chromatin ubiquitylation in the presence of 25 nM E2∼Ub (**Fig. 2**). Using these datasets, we constructed a histogram of PRC1.1 residence times during active binding events (**Fig. 4D**). The resulting distribution was exponential and yielded an average ‘active’ residence time (***τ***_res,act_) of 19.54 ± 0.4 s, matching the long chromatin residence time ***τ***_res,2_ of PRC1.1 (**Fig. 3G,H**). Increasing the concentration of both PRC1.1 and E2∼Ub (from 25 to 40 nM) did not result in an altered duration of active residence times (**Fig. 4D**). Together, these findings confirm that the long PRC1.1 residence time ***τ***_res,2_ corresponds to the enzyme being bound to the nucleosome in a catalytically competent state. In contrast, short-lived PRC1.1 bound states with residence times of ∼1 s were generally not associated with activity and represent enzymes in the process of searching for substrates or being non-specifically bound to DNA or nucleosomes.

**Fig. 4.**
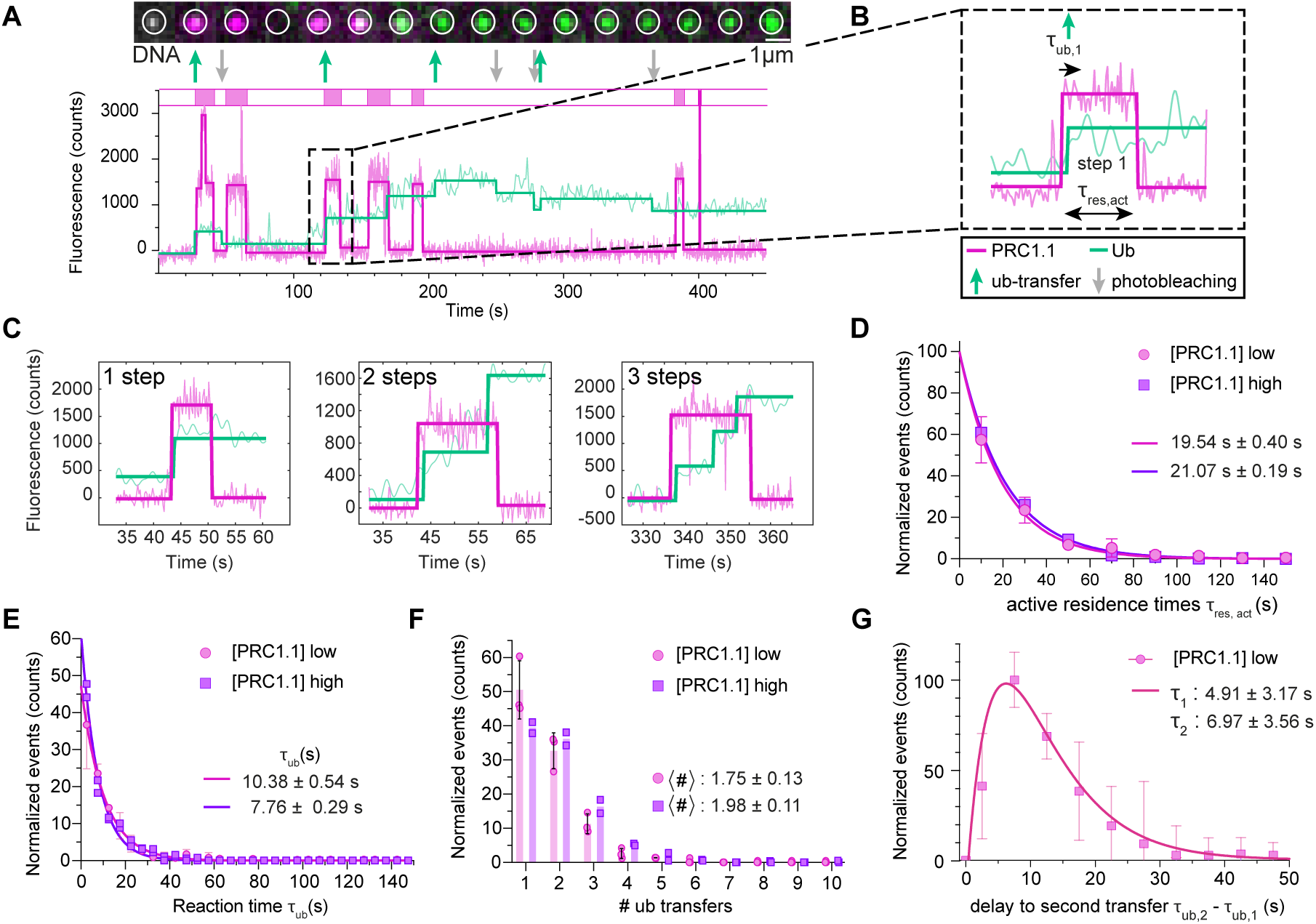
| Dissecting single chromatin writing events by PRC1.1. **A**) Top: fluorescence images from a single chromatin fiber (white circle: position of chromatin fiber, green: ubiquitin channel, magenta: PRC1 channel). Bottom: Fluorescence time trace. **B)** Binding event of PRC1.1 correlating with ubiquitylation steps. ***τ***_res,act_: productive PRC1 binding time, ***τ***_ub,i_: reaction time until i^th^ ubiquitin transfer. **C**) Examples of PRC1.1 binding events associated with indicated number of ubiquitin transfer steps. See **Fig. S12A** for more events. **D**) Histograms of binding events associated with ubiquitin transfer. [PRC1.1] low: 0.5 nM PRC1.1, 25 nM E2∼Ub, [PRC1.1] high: 0.8 nM PRC1.1 and 40 nm E2∼Ub. Counts are binned and normalized to the total bin number. Solid line: fit using mono-exponential function, with indicated fit parameters for ***τ***_res,act_. Error bars: s.d., n = 3 ([PRC1.1] low), n = 2 ([PRC1.1] high). **E**) Histograms of first reaction times, from beginning of binding to first ubiquitin transfer (***τ***_ub,1_, see **B**) for different concentrations (see **D**). Counts are binned and normalized to the total bin number, solid line: fit to mono-exponential function, with indicated fit parameters for ***τ***_ub,1_. Error bars are s.d., n = 3 ([PRC1.1] low), n = 2 ([PRC1.1] high). **F)** Histogram of the number of ubiquitin transfers (# steps) per binding event of PRC1.1 at different concentrations (see **B**). Counts are binned and normalized to the total bin number, error bars are s.d., n = 3 ([PRC1.1] low), n = 2 ([PRC1.1] high). Indicated are average steps per event<#>for different concentrations. **G**) Histogram of ***τ***_ub,2_, i.e. the time distribution between first and second ubiquitin transfer, at 0.5 nM PRC1.1, 25 nM E2∼Ub. Error bars are s.d., n = 3. Solid line: bi-exponential fit, yielding indicated time constants ***τ***_1_ and ***τ***_2_.

The ability to directly observe coupled PRC1.1 binding and activity allowed us to determine the kinetics of E2 recruitment and ubiquitin transfer upon the formation of the catalytic ternary complex on chromatin. Analyzing many individual catalytic events confirmed our initial observation of a delay between PRC1.1 binding and the emergence of fluorescence emission in the ubiquitin-channel (**Fig. 4A-B**). This not only indicates that PRC1.1:E2∼Ub complexes do not pre-form stable long-lived complexes in solution but also that PRC1.1 does not self-ubiquitylate during the time of the reaction under our conditions, which would result in co-appearance of PRC1.1 and ubiquitin signals. We then determined the delay times between PRC1.1 binding and the first ubiquitin transfer step (***τ***_ub_, **Fig. 4E**). The resulting distributions followed again a single-exponential decay, indicative of a stochastic process with a single rate-limiting step. At concentrations of 0.5 nM PRC1.1 and 25 nM E2∼Ub, we determined an apparent time constant of 10.38 ± 0.54 s for the ubiquitin transfer step, which corresponds to an estimated on-rate of 3.85 · 10^6^ M^-1^ s^-1^, assuming that E2∼Ub association to pre-bound PRC1.1 is rate-limiting. At the higher concentrations of 0.8 nM PRC1.1 and 40 nM E2∼Ub, the transfer kinetics increased, resulting in an apparent time constant of 7.76 ± 0.29 s, or an on-rate of 3.22 · 10^6^ M^-1^ s^-1^ (**Fig. 4E** and **Fig. S12C**). This direct concentration dependence of the chromatin ubiquitylation kinetics further corroborates that E2∼Ub recruitment is the rate-limiting step for PRC1.1 reactivity under our assay conditions.

Upon examining individual PRC1.1 binding and catalysis events, we observed that two or more ubiquitin transfer steps occurring in quick succession were common (**Fig. 4C,F and S12A**). While single transfers were most likely (∼50% for 25 nM E2∼Ub, ∼40% for 40 nM E2∼Ub), the probability for two transfer steps within a single binding event was also high (33% for 25 nM E2∼Ub, 36% for 40 nM E2∼Ub) (**Fig. 4F**). Again, we observed a concentration dependence, with higher propensity for more than one transfer step at higher E2∼Ub concentrations. The enzyme however did not show much processivity beyond two transfer steps, with low probability for three transfers (<15%) and above (<10%) (**Fig. 4F**). This shows that PRC1.1 does not diffuse along chromatin fibers spreading the ubiquitin mark, but remains constrained to a given site during a reaction. There, it may ubiquitylate 1-2 nucleosome sites before dissociating into the solution.

Finally, we analyzed the waiting time distribution between a first and second ubiquitylation step, for cases that exhibited more than one ubiquitin transfer per PRC1.1 chromatin binding event. The resulting distribution (**Fig. 4G**) showed a maximum at around seven seconds and could be fitted by a bi-exponential function with time constants ***τ***_1_ = 4.9 ± 3.2 s for the first process, and ***τ***_2_ = 7.0 ± 3.6 s. Such a bi-exponential distribution of waiting times indicates the presence of a hidden reaction step, which in this case could correspond to the exchange of E2∼Ub or a reorientation PRC1.1 to find a new substrate within the chromatin fiber.

### The PCGF subunit determines ubiquitylation kinetics

Different variants of PRC1 complexes display distinct chromatin ubiquitylation activities, which are determined by the specific identity of their PCGF subunit (*38*, *39*). To dissect the mechanistic origin of these differences, we further investigated a variant PRC1 complex containing PCGF4 (PRC1.4, **Fig. 5A**). PRC1.4 has been reported to be less active than PRC1.1 (*39*), although the underlying reasons for this difference in activity, particularly on chromatin substrates, remain unclear. To explore the variations in behavior among PRC1 complex variants, we recombinantly expressed, fluorescently labeled, and purified PRC1.4 using previously established methods (**Fig. 5B and S13A-H**). Indeed, when comparing the activity of PRC1.1 and PRC1.4 on reconstituted chromatin fibers in ensemble assays, we observed that PRC1.1 exhibited significantly faster H2A ubiquitylation kinetics compared to PRC1.4 (**Fig. 5C and S13I**), in agreement with earlier observations (*39*).

**Fig. 5.**
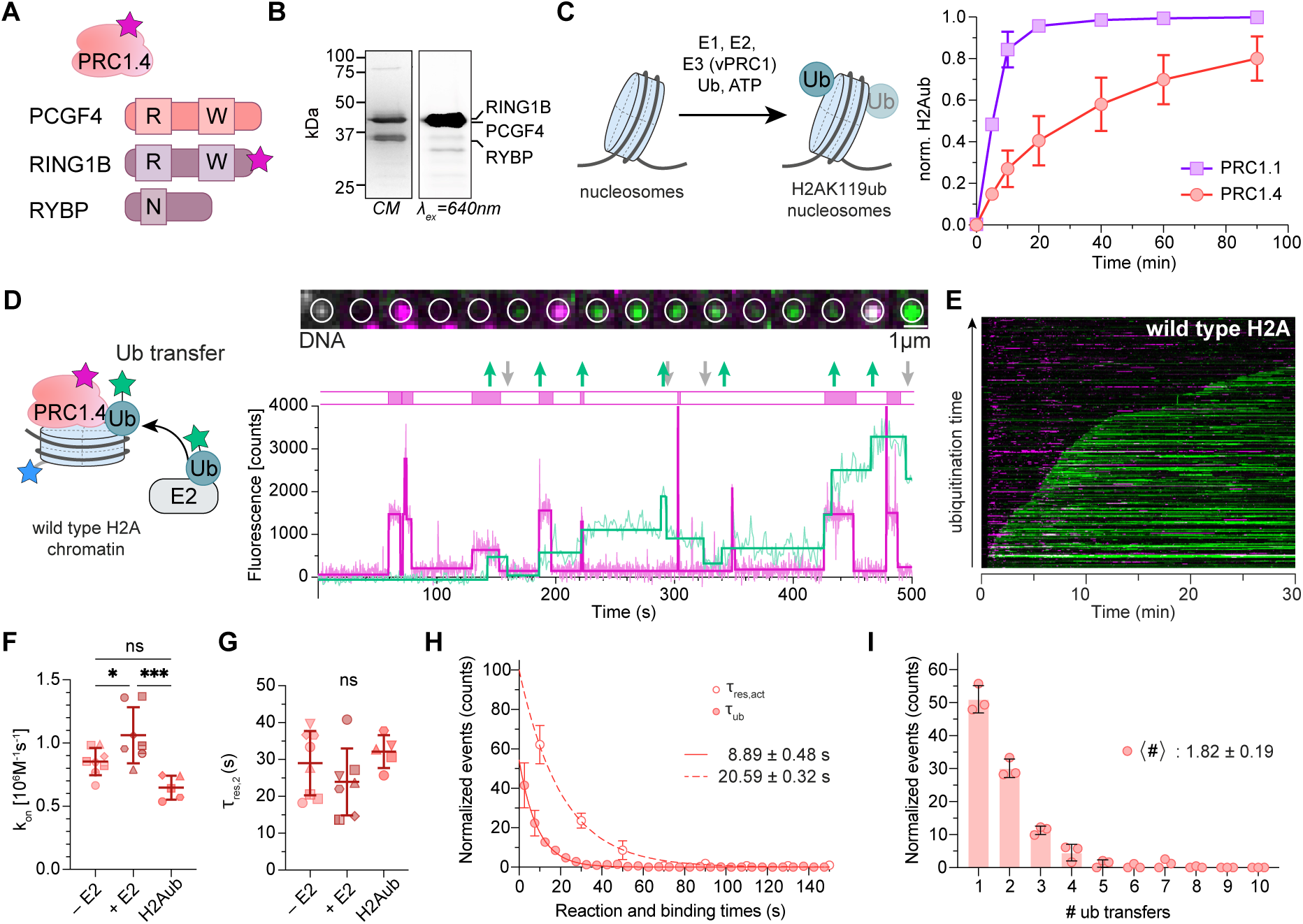
| Single-molecule analysis of PRC1.4 chromatin ubiquitylation. **A**) PRC1.4 subunit composition. The red star indicates the position of the AF_647_ dye. **B**) SDS-PAGE of purified, fluorescently labeled PRC1.4. CM: Coomassie blue, λ_ex_ = 640 nm: AF_647_ fluorescence after 640 nm irradiation. **C**) Quantification of ubiquitylated H2A from ensemble ubiquitylation assays on nucleosomes using PRC1.4 and PRC1.1 (see also **Fig. S13I**). Error bars: s.d., n = 3 independent experiments. **D**) PRC1.4 chromatin ubiquitylation experiment observed by single-molecule fluorescence. Left: Reaction scheme. Top: fluorescence images from a single chromatin fiber (white circle: position of chromatin fiber, green: ubiquitin channel, magenta: PRC1 channel). Bottom: Fluorescence time trace, see **Fig. 2B**) for details, and see **Fig. S14** for additional representative traces and images. **E**) Composite plots of PRC1.4 ubiquitylation process shown in **D**) over 30 min at [E2∼Ub] = 25 nM. n = 261 traces (see **Fig. S14E,F** for additional composite plots). **F**) k_on_ for PRC1.4 binding to chromatin under indicated conditions, for calculated dissociation constants and analysis details see **Fig. S15**. n = 8 (unmod. - E2), n = 7 (unmod. + E2), n = 5 (H2AK119ub). Error bars, s.d.; Statistical testing: one way ANOVA, Tukey’s multiple comparison test, (*) p < 0.05, (**) p < 0.01, (***) p < 0.001, ns: non-significant. **G**) Long residence times ***τ***_off,2_ for PRC1.4 on chromatin under indicated conditions. For short residence times ***τ***_off,_1 and analysis details see Methods and **Fig. S15.** For description of statistics see panel **F**). **H**) Open symbols: histograms of active binding events ***τ***_res,act_ of PRC1.4. Counts are binned and normalized to the initial bin. Fit: mono-exponential function (dashed line), error bars are s.d., n = 3. Closed symbols: Histograms of reaction times ***τ***_ub,1_ of PRC1.4. Counts are binned and normalized to the initial bin. Fit: mono-exponential function (full line), error bars are s.d., n = 3. See **Fig. S12D** for distribution of secondary transfer events ***τ***_ub,2_. **I**) Histogram of the number of ubiquitin transfers (# steps) per binding event of PRC1.4. Counts are binned and normalized to the initial bin, error bars are s.d., n = 3. For all data see **Table S2**.

We then resolved chromatin association and H2A K119 ubiquitylation by PRC1.4 using single-molecule detection. We employed the previously established conditions with Alexa647-labeled PRC1.4 at a concentration of 0.5 nM, in the presence of 25 nM JF549-labeled E2∼Ub (**Fig. 5D**). Similar to the reaction with PRC1.1, we observed transient PRC1.4 binding events to immobilized chromatin fibers, associated with stepwise ubiquitin transfer reactions (**Fig. 5D,E** and **S14A,C**). Mutating lysines 119 and 120 in H2A to arginine greatly delayed chromatin ubiquitylation, indicating that also PRC1.4 exhibits high specificities for these residues (**Fig. S14B,D**).

Our assay enabled the comparison of binding and reaction kinetics between PRC1.1 and PRC1.4, aiming to gain mechanistic insights into why PRC1.4, which contains a PCGF4 subunit, is associated with reduced chromatin ubiquitylation capacity. In particular, we wondered if binding kinetics or residence times are reduced for PRC1.4, if its ability to recruit E2∼Ub is diminished, or if PRC1.4 exhibits slower E2 discharge and ubiquitin transfer kinetics. Focusing on chromatin binding dynamics, we observed similar residence times between PRC1.4 and PRC1.1 both in the presence or absence of E2, or on chromatin carrying H2AK119ub (**Fig. 5F and S15**). Similar to PRC1.1, PRC1.4 exhibited a trend towards increased residence time on H2AK119ub chromatin, although this difference was not statistically significant (**Fig. 5F**). Moreover, PRC1.4 exhibited faster on-rates compared to PRC1.1 (**Fig. 5F**), resulting in a lower dissociation constant, and thus higher affinity. Overall, the differences in chromatin binding properties therefore cannot account for the variations in reactivity between the two complexes.

We next focused on the correlation between chromatin binding and ubiquitin transfer kinetics. Productive binding events of PRC1.4, resulting in chromatin ubiquitylation, lasted on average ***τ***_res,act_ = 20.59 ± 0.32 s, similar to the 19.54 s ± 0.40 s observed for PRC1.1 (**Fig. 5H)**. When determining the average time between PRC1.4 binding and detected ubiquitin transfer reaction, we determined ***τ***_ub,1_ = 8.89 s ± 0.5 s, very similar to PRC1.1 (**Fig. 5G** and **4D,E**). Moreover, we observed on average 1-2 transfer steps per binding PRC1.4 event with the highest probability of ∼50% for 1 ubiquitin transfer (**Fig. 5I**), indicating that the processivity of PRC1.4 is comparable to PRC1.1 (**Fig. 4F**). In summary, when exclusively focusing on individual active binding events associated with ubiquitin transfer reactions, PRC1.1 and PRC1.4 exhibit no significant difference in behavior.

**Fig. 6.**
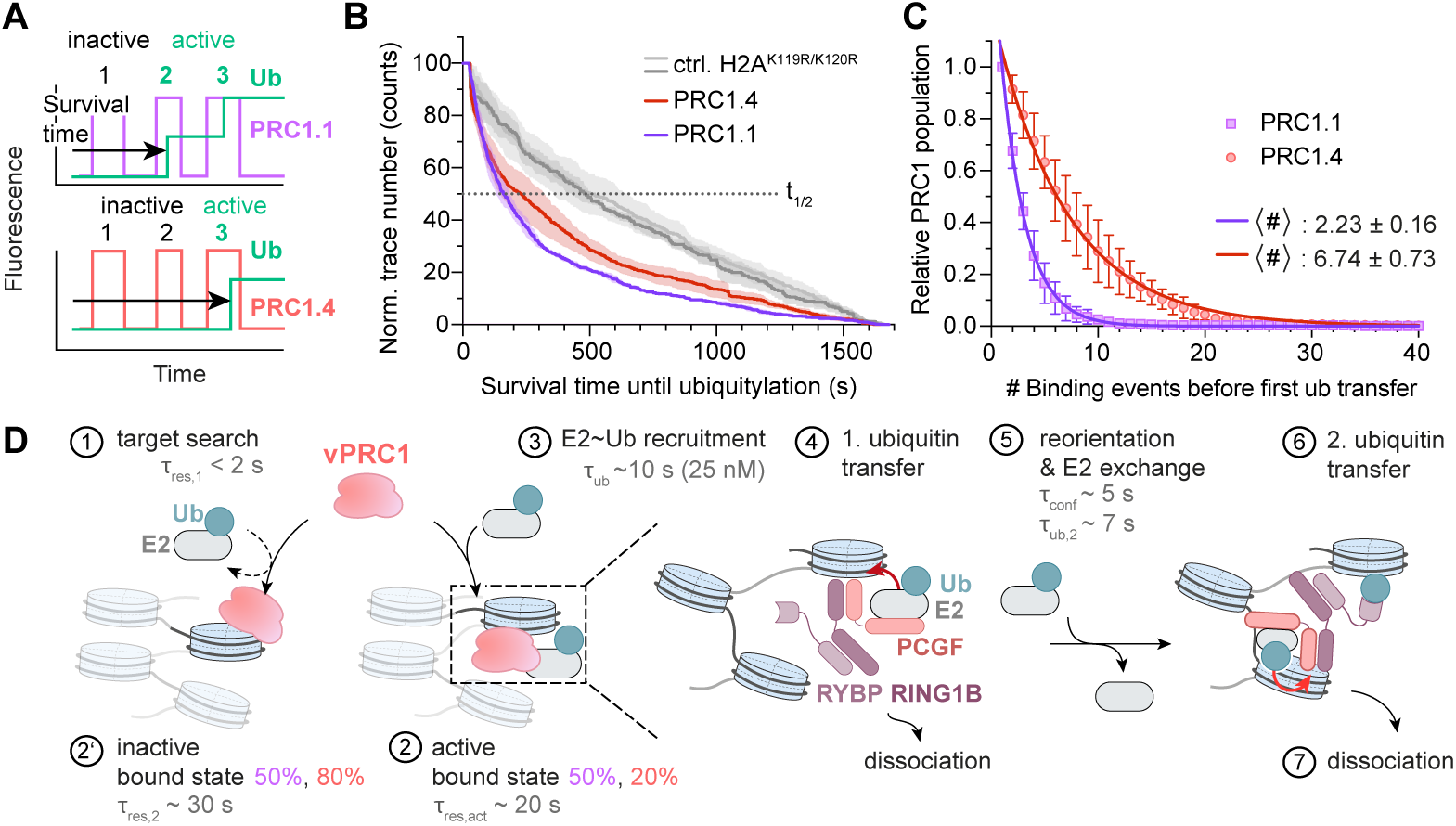
| PRC1.4 has a lower probability to form enzymatically active chromatin interactions. **A)** Analysis to determine the probability of ubiquitin transfer per variant PRC1 binding event. Survival time until ubiquitylation: Time delay between injection of variant PRC1 complexes and E2∼Ub to first chromatin ubiquitylation. (In-)active events: Variant PRC1 binding events (not) associated with ubiquitin transfer. **B**) Distribution of survival times until ubiquitylation, as defined in **A**) for PRC1.1 on wild-type chromatin (H2A^wt^, n = 3) and control chromatin (H2A^K119R/K120R^, n = 3), and PRC1.4 on wild-type (n = 3) and control chromatin (n = 3), data from **Fig. 2E, 5E**. Data is mean (solid line) with ± SEM, n=3. Observed half-times t_1/2_ are for PRC1.1: H2A^wt^ : 157 s, H2A^K119R/K120R^: 484 s; PRC1.4 : H2A^wt^ : 231 s, H2A^K119R/K120R^ : 507 s. **C**) Distributions of the number of inactive binding events before the first ubiquitylation event for PRC1.1 (n = 5) and PRC1.4 (n = 5). Analysis: mono-exponential fit, with indicated parameters for PRC1.1 and PRC1.4. **D**) Overall model and associated timescales for variant PRC1 chromatin ubiquitylation. Percentages in violet: population of PRC1.1, percentages in red: populations of PRC1.4. For further details, see text.For all data see **Table S3**.

While both binding kinetics and subsequent reaction dynamics are similar between the two complexes, it was evident that the ubiquitin signal emerges with an extended delay for reactions catalyzed by PRC1.4 (**Fig. 6A**). Indeed, when examining individual kinetic traces of reactions involving PRC1.4, many binding events did not lead to reactions and the enzyme dissociated from chromatin before any histone ubiquitylation could occur. While quantifying the ubiquitylation time across multiple traces, we observed that the global reaction time of PRC1.1 was significantly faster than for PRC1.4 (**Fig. 6B**). We thus determined the number of non-productive binding events preceding a ubiquitin transfer reaction for both variant PRC1 complexes. Under the selected conditions with 25 nM E2∼Ub, the resulting distributions were exponential, and the data revealed that PRC1.1 required an average of ∼2 trials before a ubiquitin transfer occurred (**Fig. 6C**). In contrast, PRC1.4 engaged chromatin approximately 7 times on average before initiating the first ubiquitylation reaction. This increased probability of non-productive chromatin binding events thus accounts for the reduced activity of the PRC1.4 complex.

## Discussion

The action of chromatin modifying enzymes is critical to shape the chromatin landscape. Detailed mechanistic information is required to be able to understand their regulation. Here, we have established a multicolor smTIRF approach to directly observe how a chromatin modifying enzyme engages both in chromatin reading and PTM writing dynamics, on the single-molecule level with high temporal precision. We focused on elucidating the activity of variant PRC1 complexes on defined chromatin templates. This allowed us to analyze the sequential steps of ubiquitin transfer from E2∼Ub to chromatin, catalyzed by two different variant PRC1 complexes, PRC1.1 and PRC1.4, and to dissect the contribution of their distinct PCGF subunits to catalysis. We find that PRC1.1 has much higher activity, dependent on its PCGF1 subunit. This is consistent with its role, together with PRC1.3 and PRC1.5, as the main driver of H2AK119 ubiquitylation both genome-wide and focused at key locations(*58*). In contrast, PRC1.4 contains a PCGF4 subunit, which is also a key subunit of canonical PRC1, and exhibits much lower ubiquitylation activity(*39*). As canonical PRC1 plays other key roles, e.g. compacting chromatin for gene repression(*59*), a low maintenance-level activity of H2AK119 ubiquitylation might be sufficient for this complex. Moreover, as H2A ubiquitylation actually inhibits chromatin compaction and high local H2Aub concentrations results in disruption of polycomb repression (*60*), reduced ubiquitylation activity may actually be beneficial for local repression.

Interestingly, our measurements do not reveal key differences in chromatin recruitment between the two complex types. Rather, the origins for their different activities lie in their ability to functionally engage chromatin. Our single-molecule observations allow us to construct an overall model of the interaction and reaction dynamics of variant PRC1 on chromatin (**Fig. 6D**). Both variant PRC1 complexes search and engage chromatin, forming transient interactions that last below ∼ 2 seconds (step 1), but generally do not result in E2 recruitment and catalysis, i.e. chromatin ubiquitylation. These states may involve variant PRC1 complexes interacting with nucleosomal DNA or histone residues without engaging the nucleosome surface and acidic patch in a catalytically competent manner. However, these transient complexes can transition into more stably bound states (steps 2 and 2’), which last 20 - 30 seconds, which is consistent with the binding time range observed for PCGF3 and PCGF6 using single-particle tracking (SPT) in ESCs(*47*). Based on our observation that a significant subset of stably bound states do not result in ubiquitin transfer, we hypothesize that not all of these states correspond to variant PRC1:nucleosome complexes in a catalytically competent conformation. In particular, the presence of the PCGF4 subunit in PRC1.4 results in a lower probability of forming such active complexes (20% vs. 80% for PCGF1). When an active complex is formed, however, downstream kinetics are similar for different variant PRC1 E3 ligases. E2∼Ub recruitment results in ubiquitin transfer to a target H2A K119 lysine within several seconds, dependent on the overall concentration of preloaded E2. E2∼Ub recruitment appears to be the rate-limiting step as it directly depends on E2∼Ub concentration and exhibits an apparent binding rate in the range of ∼3 · 10^6^ M^-1^ s^-1^ (steps 3-4). After an initial ubiquitylation event, variant PRC1 complexes dissociate with a high probability. In a subset of events, the enzyme can however reorient and engage a second lysine on the same histone, nucleosome, or even a neighboring nucleosome to catalyze a second transfer event (steps 5-6). H2A can be diubiquitylated, as there are several lysines in close proximity, and this is observed both *in vitro* on nucleosomes and on chromatin fibers (**Fig. S4, 8**), and is also observed *in vivo*(*54*). Moreover, the RYBP subunit can engage ubiquitylated H2A, it may serve as an ‘anchor’ enabling variant PRC1 complexes to remain bound to chromatin while engaging nucleosomes in the vicinity. This mechanism is supported by recent structural data on PRC1.4(*27*). Still, processive activity beyond two successive ubiquitylation reactions is rare, indicating that variant PRC1 complexes rather scan chromatin and deposit H2AK119ub in a distributive fashion.

Our results show that E3 ligase-chromatin complexes remain highly dynamic and can adopt multiple conformations with different lifetimes and reactivities. Recent structural studies on related RING type ligases, e.g. RNF168, provided structural data on the conformational heterogeneity of nucleosome-bound complexes(*61*). Here, we resolve the kinetics of state lifetimes and interconversion. A key determinant for variant PRC1 activity is the ability of the ligase to engage nucleosomes in a catalytically competent conformation. This depends on the nature of the PCGF subunit, and may involve either forming key contacts with the nucleosome, the ability of positioning E2∼Ub for ubiquitin transfer or recruiting the ubiquitin-conjugating enzyme in the first place. Previous studies comparing PCGF subunits have identified amino acid residues in the PCGF RING domain, which contact the interface of the E3:E2∼Ub complex thus having a significant impact on E2 discharge kinetics in the absence of substrate (*38*). However, effects of these mutants on activity towards chromatin substrates were less obvious. We thus hypothesize that structural heterogeneity of the variant PRC1:nucleosome complexes may play a key role, with PCGF1 promoting an active conformation, while further structural and mutational studies may be required to resolve this regulatory mechanism.

In summary, here we present a single-molecule multi-color approach to explore how ‘reader’ and ‘writer’ complexes dynamically modify chromatin. We reveal that variant PRC1 complexes transiently explore chromatin until they achieve a catalytically competent state for histone H2A ubiquitylation, with subtype-specific activity differences driven by the likelihood of forming this active complex. We expect that such methodology will be important in the future to further explore the complex interplay between chromatin recognition, modification and remodeling for a molecular view on chromatin regulation.

## Methods

### Experimental design

Our study was designed to directly observe the mechanism of chromatin ubiquitylation catalyzed by the E3 ligase variant PRC1 on a single-molecule level, focusing on key dynamic parameters such as the correlation between chromatin binding and enzymatic activity, ubiquitin transfer rates and the processivity of the enzyme, dependent on the variant PRC1 subtype. To this end, we established a single-molecule fluorescence total internal reflection microscopy (smTIRF) system to detect both variant PRC1 binding, E2 recruitment and ubiquitin transfer kinetics from fluorescence time traces. For all experiments we used a highly defined system composed of recombinant PRC1 complexes, and synthetic chromatin substrates.

### Histone octamer production

#### Expression and purification of recombinant histones

Human core histones (wt H2A, wt H2B, H3.2C110A, and wt H4; H2AN110C, H2AK119C, H2AK119R/K120R) were expressed and purified without purification tags according to published literature (*62*). pET3a (Amp^R^) expression plasmids were transformed into BL21 (DE3) pLysS E.coli (Cam^R^). A pre-culture was grown overnight at 37 °C in LB medium containing 100 µg/l ampicillin and 34 µg/l chloramphenicol. The main culture was inoculated (1:20), grown to an OD_600_ = 0.6 and induced with 0.5 mM IPTG to express the histones for 2.5 h to 3 h at 37 °C. Cells were harvested by centrifugation at 8000 rpm for 10 min at 4 °C and lysed by sonication (8×15 s bursts, 30s off, at 40% amplitude) in lysis buffer (20 mM Tris pH 7.5, 1 mM EDTA, 200 mM NaCl, 1 mM β-mercaptoethanol (βME), 100 µg/l lysozyme and 1x cOmplete™ Protease Inhibitor Cocktail tablets Roche). The insoluble fraction was pelleted by centrifugation at 12,000 rpm for 30 min at 4 °C. Histones were resolubilized from inclusion bodies in resolubilization buffer (6 M guanidine HCl, 20 mM Tris pH 7.5, 1 mM EDTA, 1 mM βME) and stirred for 2 h at 4 °C. The supernatant was cleared by centrifugation and filtered through a 0.45 µm filter. Dialysis was performed against urea buffer (7 M urea, 100 mM NaCl, 10 mM Tris pH 7.5, 1 mM EDTA, 1 mM βME, 0.2 mM PMSF) in three steps á 12 h, followed by centrifugation to remove aggregates. The resolubilized histone fractions were subjected to cation exchange chromatography using a 5 mL HiTrap™SP HP column. After washing in ion exchange buffer (7 M urea, 100 mM NaCl, 10 mM Tris pH 8, 1 mM EDTA, 1 mM DTT, 0.2 mM PMSF), histones were eluted with a gradient of 0 % to 30 % high-salt buffer (7 M urea, 1.5 M NaCl, 10 mM Tris, 1 mM EDTA, 1 mM DTT, 0.2 mM PMSF, pH 8) and analyzed via SDS-PAGE and Coomassie blue staining. Fractions containing pure histones were pooled, dialyzed into water, and lyophilized for storage. Final purification was performed via Agilent 1260 preparative HPLC system with a Zorbax 300SB-C18 preparative column (7 µm, 21.2 x 250 mm) with a gradient of 0 % to 70 % solvent B. The purified fractions were analyzed by ESI LC-MS, pooled, lyophilized, and stored at −20 °C until used for octamer refolding.

#### Cysteine-directed fluorescent labeling of proteins

The cysteine-containing proteins (including Cys-Ub, H2A N110C) were dissolved in PBS and treated with 1 equivalent of buffered tris-[2-carboxyethyl]phosphine (TCEP) at pH 7-7.5 at 25°C and agitation of 300 rpm for 20 min to reduce the cysteine residues. Subsequently, a 2-5 fold excess of fluorescent dye-maleimide conjugate (stocks in DMF or DMSO) was added and the reaction incubated at pH 7-7.5 at 25°C. Reaction progress was monitored by RP-HPLC analysis on Agilent Zorbax 300SB-C18 column (5 µm, 4.6 x 150 mm) before purification on Agilent Zorbax 300SB-C18 semi-preparative column (5 µm, 9.4x 250 mm). The peak corresponding to the protein-dye conjugate was collected, confirmed by ESI LC-MS, lyophilized, and stored at −20°C.

#### Octamer refolding

Equimolar amounts of human H3.2 containing the C110A mutation (H3C110A) and wild type human H4 (wt hH4) were mixed in unfolding buffer (6 M Guanidine HCl, 20 mM Tris, 5 mM DTT) with 1.1 equivalents of wt H2A and wt H2B, and the final concentration was adjusted to 1 mg/ml. Histone octamer refolding was performed by dialysis in a Slide-A-Lyzer Dialysis Cassette (7000 MW, 0.5-3 ml capacity, from Thermo Scientific) against refolding buffer (2 M NaCl, 10 mM Tris, 1 mM EDTA, 1 mM DTT, pH 7.5). Histone aggregates were removed by centrifugation at 15,000 rpm for 10 min at 4 °C. Refolded octamers were subsequently purified by size exclusion chromatography (SEC) on a Superdex S200 10/300GL column (GE Life Sciences) and analyzed on SDS-PAGE and RP-HPLC. Pure octamer-containing fractions were combined and concentrated to a volume of approximately 50 µM. Glycerol was added to a final concentration of 50%, and the octamer stocks were stored at −20 °C for later use.

### Preparation of H2AK119ub

H2AK119ub was prepared following published protocols (*56*, *57*) with some modifications as shown below.

#### Expression of Aha75Cub and purification of Ub-N3

A pGEX2TK plasmid encoding human UbG75MΔG76 was obtained as a gift from the laboratory of Andreas Marx (University of Konstanz). Methionine auxotrophic E. coli B834 (DE3) cells were transformed with pGEX2TK-G75MUb and cultured in LB medium with 100 mg/L carbenicillin at 37°C overnight. The pre-culture was diluted with new minimal medium (NMM): 20 mg/L of each natural amino acid (except methionine), 7.5 mM (NH_4_)_2_SO_4_, 8.5 mM NaCl, 22 mM KH_2_PO_4_, 50 mM K_2_HPO_4_, 1 mM MgSO4, 1 mg/L CaCl_2_, 1 mg/L FeCl_2_, 1 μg/L CuCl_2_, 1 μg/L MnCl_2_, 1 μg/L ZnCl_2_, 1 μg/L Na_2_MoO_4_, 3.6 g/L glucose, 10 mg/L thiamine, 10 mg/L biotin, and 100 μg/ml carbenicillin. An additional 0.06 mM Met were added to NMM and cells were grown to an OD600 of 1.3, harvested, and resuspended in fresh NMM with 0.5 mM L-Azidohomoalanine (Aha). After 30 min at 37°C, protein expression was induced with 1 mM IPTG, and cells were incubated overnight at 25°C. Cells were pelleted, resuspended in PBS with 1% Triton X-100, and lysed by sonication (3 min,2 s on/3 s off, at 35% amplitude). The lysate was clarified by centrifugation (50,000 g for 30 min at 4°C), and the supernatant was incubated with glutathione agarose beads at 4°C for 5 h. Beads were packed into a column, washed with PBS, and incubated with 10 units of thrombin at room temperature overnight. The released Aha75CxUbs were eluted with PBS, yielding over 5 mg of protein per liter of culture. Protein purity was analyzed by SDS-PAGE, Coomassie blue staining, concentration was measured using the UV-Vis and the pooled fractions were stored at −80 °C.

#### Michael addition of propargyl acrylate to H2AK119C

Lyophilized H2AK119C was dissolved in 20 mM Tris pH 7.5 and 150 mM NaCl (6 mg, 180 mM, 1 eq). 50 eq of propargyl acrylate (PA) diluted in acetonitrile was added, and the reaction was continued for 45 min. Reaction progress was monitored by RP-HPLC analysis on Agilent Zorbax 300SB-C18 column (5 µm, 4.6 x 150 mm) and MS analysis was conducted using direct injection (200-2000 m/z) on a Shimadzu LCMS-2020. H2AK119C-PA was purified on an Agilent Zorbax 300SB-C18 semi-preparative column (5 µm, 9.4x 250 mm). The peak corresponding to the protein-dye conjugate was collected, confirmed by ESI LC-MS, lyophilized, and stored at −20°C.

#### CuAAC click reaction between Ub-N3 and H2A-PA

Click reaction was performed in 20 mM Tris pH 7 and 20% DMSO mixture containing 10 mM THPTA, 5 mM CuSO4.5H2O, 20 µM H2A-PA by addition of 30 µM Ub-N3 (ratio 1:1.5). The reaction was monitored over 60 min by RP-HPLC analysis on Agilent Zorbax 300SB-C18 column (5 µm, 4.6 x 150 mm) followed my MS. The reaction was quenched by adding 50 mM Na L-ascorbate, followed by purification on Agilent Zorbax 300SB-C18 semi-preparative column (5 µm, 9.4x 250 mm). The peak corresponding to the protein-dye conjugate was collected, confirmed by ESI LC-MS, lyophilized, and stored at −20°C.

### Chromatin DNA generation

#### Large-scale generation of plasmid DNA

The plasmid containing 12×601 nucleosome positioning sequences was constructed through sequential restriction digests and DNA ligation, validated by test digest experiments (*48*). DH5α E.Coli cells were transformed with the plasmid (e.g. 12×601, or 8xMMTV), and individual clones were isolated, expanded, DNA was isolated using the QIAprep Spin Miniprep Kit (QIAGEN), analyzed by restriction digest and sequencing and glycerol stock preparation. A pre-culture was grown in 2×YT medium before scaling up to 6 l of 2×YT medium. The culture was incubated overnight at 37 °C and 200 rpm. Cells were harvested by centrifugation at 4000 g for 10 min at 15 °C. Cells were lysed using a series of buffers 120ml of lysis buffer I (50 mM glucose, 25 mM Tris pH 8, 10 mM EDTA), 240ml of lysis buffer II (0.3 M NaOH, 1% SDS) and 240 ml lysis buffer III (neutralization) (4 M KAc, 2 M acetic acid). The mixture was incubated at 4 °C for 15 min. Supernatants were filtered and subjected to isopropanol precipitation. The DNA pellet was dissolved in TE 10/50 buffer (10 mM Tris pH 7.5, 50 mM EDTA). Residual RNA was digested using 200 units of RNase A at 37 °C for 2 to 4 h. The solution was then adjusted to 2 M KCl in TE 10/50 buffer for size exclusion chromatography (SEC) using a XK 50/30 column packed with Sepharose 6 Fast Flow (GE Healthcare). Eluted fractions were analyzed on a 1% agarose gel, pooled, isopropanol precipitated, and the final DNA pellet was resuspended in TE 10/50 buffer and stored at −20 °C. The plasmid DNA was verified through sequencing and restriction digest tests using EcoRI, EcoRV, BsaI, and HindII enzymes from NEB.

#### Large-scale restriction digest and purification

Purified plasmid DNA was subjected to ethanol precipitation to remove residual EDTA from TE 10/50 buffer by addition of ice-cold 100% ethanol and 5 mM NaCl to achieve a final concentration of 75% ethanol and 3 mM NaCl, respectively. The mixture was then incubated at −20 °C for 20 min, centrifuged at 13.000 rpm for 30 min at 4 °C, washed with 70% ethanol, and finally dissolved in Milli-Q water. A typical digest reaction involved 500 µg to 1 mg of plasmid DNA in a total volume of 1 ml. 400 units of EcoRV-HF (New England Biolabs) and 25 units of Quick CIP were added for simultaneous dephosphorylation in 1× CutSmart buffer. The digest was carried out at 37 °C for 16 to 24 h. Progress was monitored by agarose gel electrophoresis on a 1% gel in 1× TBE buffer. If incomplete, an additional 100 units of EcoRV-HF and more CutSmart buffer were added, and the mixture was incubated for an additional 2 to 4 h at 37 °C. An additional 25 units of CIP were added for 30 min to ensure complete dephosphorylation and prevent re-ligation. The enzymes were then deactivated at 80 °C for 20 min. Following digestion, DNA was purified through successive rounds of PEG precipitation to remove the vector backbone. Varying percentages of PEG 6000 were added to achieve a final concentration of 4-6.5%, adjusting the NaCl concentration to 0.5 M. The mixtures were incubated on ice for 30 min and DNA fragments were collected by centrifugation at 13.000 rpm for 30 min at 4 °C. The supernatant was processed with additional PEG 6000 in 0.5% increments to purify the DNA, repeating the precipitation process to yield highly pure DNA fragments suitable for downstream applications. Chromatin DNA fragments were then isolated using QIAquick PCR purification spin columns (QIAGEN).

#### Oligonucleotide Labeling

Synthetic oligonucleotides were subjected to 2x ethanol precipitation. For the labeling reaction, 5 nmol to 10 nmol of oligonucleotide were dissolved in 50 µl of 0.1 mM sodium tetraborate buffer, pH 8.5. 5 eq to 10 eq of N-hydroxysuccinimide (NHS) ester-modified fluorophores were added. The mixture was incubated overnight at 25°C and 300 rpm in the absence of light. Reaction progress was monitored by RP-HPLC on an InertSustain C18 column (GL Sciences, 3 µm, 4.6 x 150 mm) using a 20 min gradient from 0 to 70% Solvent B (0.1 M triethylammonium acetate (TEAA) pH 7 and acetonitrile (ACN)) at 1 ml/min, detecting at 260 nm. The labeled oligonucleotide was purified by RP-HPLC, precipitated with ethanol to remove excess dye, dissolved in Milli-Q H_2_O to 2.5 mM, and stored at −20 °C.

#### Fluorophore and biotin attachment to 12×601 DNA

A biotin-linked oligonucleotide (5’-Biotin-21bp) was annealed to its complementary strand, carrying a 5’-phosphorylated BsaI overhang (iPhospho-12bp-iAmMC6T-12bp) previously labeled with Alexa Fluor 488 using an NHS-ester labeling protocol, in 1×T4 ligase buffer. Following this, 150 µg of chromatin DNA, previously digested with EcoRV-HF, was digested with BsaI-HFv2 (NEB) to introduce necessary overhangs for ligation, using 600 units of enzyme in 1 ml total volume at 37 °C for 16-24 hours and purified using QIAquick PCR purification columns (QIAGEN). For ligation, a 5-10 fold excess of the annealed and labeled chromatin anchor was added to 130 µg of BsaI-digested chromatin DNA in 400 µl of 1× T4 ligation buffer along with 50 units of T4 ligase (NEB) per 1ug of chromatin DNA, and the reaction was conducted at 25 °C in the absence of light. The ligation’s progress was monitored by 1% agarose gel electrophoresis in 1× TBE. The excess biotin anchor was removed via PEG precipitation, and the ligated DNA was further purified using QIAquick PCR purification (QIAGEN).

#### Preparation of 1×MMTV DNA

MMTV buffer DNA used for chromatin assembly was prepared from a plasmid containing an 8-mer repeat of the sequence (8×MMTV DNA). Large-scale plasmid cultures were prepared, followed by digestion using EcoRV-HF and Quick CIP enzymes (NEB). The mixture was incubated overnight at 37 °C under agitation. Enzymes were subsequently inactivated by heating at 80 °C for 10 min. The 1× MMTV fragments of 151 bp, resulting from the digest, were separated from the 2.5 kbp plasmid backbone using PEG precipitation, and any residual PEG was removed using PCR purification kits from QIAGEN.

### In vitro reconstitution of nucleosomes and chromatin arrays

Nucleosomes and chromatin arrays carrying up to 12× 601 nucleosome positioning sequences (NPSs) with 30bp linkers were reconstituted through a gradual buffer exchange via dialysis. Octamers were combined with DNA in ratios ranging from 1:1 up to 1:3 DNA:histone octamer in TEK2000 buffer (10 mM Tris, 1 mM EDTA, 2 M NaCl). Additionally, 0.25 eq MMTV DNA, which has a lower affinity for mononucleosome formation compared to 1× 601 DNA, was supplemented ot the chromatin assembly reactions to prevent fiber oversaturation. Overnight dialysis was performed using a peristaltic pump into TEK10 buffer (10 mM Tris, 1 mM EDTA, 10 mM NaCl). Aggregates were removed by centrifugation at 15,000 rpm for 10 min at 4 °C. The nucleosome / chromatin concentration was then determined via UV-VIS spectroscopy. The quality of chromatin fibers was evaluated based on the appearance of MMTV nucleosomes and the results of ScaI digestion, which visualizes single nucleosome formation. Approximately 0.6 pmol of undigested chromatin was analyzed on a 0.6% agarose gel in 0.25× TBE. Another 0.6 pmol of chromatin underwent ScaI digestion to assess nucleosome formation efficiency. The quality of both mononucleosome reconstitution and digested chromatin was further examined via native PAGE on 5% acrylamide gels in 0.5× TBE.

#### MNase digest to analyze chromatin conformation

MNase was diluted to 20 units/µL in 1x MNase buffer, with 2 units/µL per 1.5 pM chromatin used. A master mix of chromatin and MNase was prepared in low-binding Eppendorf tubes, diluted with TEK 10 (10 mM Tris HCl, 10 mM KCl, 100 µM EDTA) supplemented with 1x MNase buffer, 1% BSA before MNase addition. The tubes were incubated on ice. At indicated time points, a 12 µL aliquot was taken and quenched by addition to 150 µL PB and 10 µL of 3 M Na Acetate. The DNA was subsequently purified using MiniElute PCR purification columns and eluted in 30 µL Milli-Q H_2_O. For gel electrophoresis, samples with equivalent DNA amounts were mixed with 5% sucrose. A Criterion gel was set in the corresponding running chamber and pre-run at 200 V in 0.5X TBE buffer on ice for 10 min. Samples were loaded, and the gel was run at 200 V for 1 hour.

### Expression, labeling and purification of variant PRC1 complexes

#### Generation of baculovirus expression constructs

The Strep and hexahistidine-tagged variant PRC1 constructs carrying full-length human variant PRC1 complex subunits, specifically PRC1.1 (Ring1B-ybbR-Strep, PCGF1, 6xHis-RYBP), PRC1.4 (Ring1B-ybbR-Strep, PCFF4, 6xHis-RYBP), PRC1.1ΔRYBP (Ring1B-ybbR-Strep, PCGF1-6xHis), or PRC1.4ΔRYBP (Ring1B-ybbR-Strep, PCGF4-6xHis), were cloned into polycistronic pBIG1a plasmid (GenScript, Rijswijk, Netherlands) (Amp^R^, Sm^R^, Gm^R^). Baculovirus particles were generated after the instructions of Bac-to-Bac TOPO expression system (Invitrogen).

#### Variant PRC1 expression and purification

High Five™insect cells (Invitrogen) were grown to 1.5-2×10^6^ cells/ml. The cells were infected with the produced virus coding for all of the required subunits of variant PRC1 complex using a multiplicity of infection (MOI) of 5-10. Cells were grown at 27 °C at 130 rpm for 72 h before harvesting by centrifugation (4,000g, 10 min at 4 °C). At this step, the cell pellets were flash frozen and kept at −80 °C until further use. Pellets from 500 ml culture were thawed at room temperature for one hour and lysed in 50 ml of lysis buffer (50 mM Tris, pH 7.5, 500 mM NaCl, 20 % glycerol, 4 mM MgCl_2_, 10 µM ZnCl_2_, 0.10 % NP-40, 0.1 mM PMSF, 5 mM β-ME, Two cOmplete™ Protease Inhibitor Cocktail tablets Roche (Merck, Cal. no. 11697498001). Cells were lysed via sonication at 4°C, cleared by centrifugation at 150,000 g for 1 h at 4 °C, and the supernatant was filtered (using 0.45 µm pore size). The clarified lysate was loaded onto a 5 ml HiFliQ Co-NTA column connected to an ÄKTA FPLC system. The column, pre-equilibrated in binding buffer (50 mM Tris, 300 mM NaCl, 4 mM MgCl_2_, 10 µM ZnCl_2_, 5 mM β-ME, pH 7.5), was washed and then subjected to an elution gradient using same buffer containing 250 mM imidazole. Combined elution fractions were further purified on a gravity flow column packed with Strep-Tactin Superflow resin at 4 °C. After loading and washing with binding buffer, proteins were eluted in binding buffer containing 5 mM desthiobiotin and 0.5 mM DTT. Fractions were analyzed by SDS-PAGE. Elution fractions containing the variant PRC1 complex were concentrated on Amicon Ultra centrifugal filters (4ml, MWCO 50 kDa) to 10 µM.

#### ybbR-tag labeling of variant PRC1 complexes

To achieve fluorescent labeling using the ybbR peptide tag, a dye must first be conjugated to coenzyme A (*49*). Coenzyme A trilithium salt (CoA-SH, Applichem) was mixed with 1.1 equivalent TCEP in PBS and incubated at 25°C for 20 min to reduce the cysteine residues. Following the reduction step, ∼0.2 equivalents of fluorescent dye-maleimide conjugate (stocks in DMF or DMSO) was added to the reaction and the reaction incubated at pH 7.4-7.5 at 25°C in the absence of light. Reaction progress was monitored by RP-HPLC analysis on Agilent Zorbax 300SB-C18 column (5 µm, 4.6 x 150 mm) before purification on Agilent Zorbax 300SB-C18 semi-preparative column (5 µm, 9.4x 250 mm). The peak corresponding to the protein-dye conjugate was collected, confirmed by ESI LC-MS, lyophilized, dissolved in Milli-Q H_2_O and stored at −20 °C. Concentration was determined by UV spectroscopy. For labeling of the ybbR-tagged variant PRC1 complex the protein solution was mixed with 2 equivalents of CoA-conjugate and 0.5-1 µM recombinant Sfp synthase in labeling buffer (50 mM HEPES, 10 mM MgCl_2_, 150 mM NaCl, 1 mM DTT). The mixture was incubated overnight at 4°C in the dark. Labeling efficiency was assessed by SDS-PAGE and ESI LS-MS

#### Size exclusion chromatography of variant PRC1 proteins

The obtained labeled or non-labeled variant PRC1 complexes were concentrated using Amicon Ultra centrifugal filters (500ul, MWCO 50 kDa) to 500 µl in SEC buffer (50 mM Tris, 300 mM NaCl, 4 mM MgCl_2_, 10 µM ZnCl_2_) supplemented with 0.25 mM TCEP and loaded on Superdex 200 10/300 GL. The fractions corresponding to pure protein complexes were analyzed on SDS-PAGE, pooled and concentrated on Amicon Ultra centrifugal filters (500ul, MWCO 50 kDa) to a target concentration of approximately 10 µM in about 200 µl. Glycerol was added to a final concentration of 15% by weight, and the protein concentrations were measured on a UV-Vis spectrophotometer. Samples were aliquoted in 5 µl volumes, flash-frozen, and stored at −80°C. The quality of variant PRC1 preparations was ensured by analysis via Western blotting, high-resolution LC-MS, and native MS before the assays.

### Production of fluorescently labeled E2∼Ub

#### Ubiquitin expression and purification

The cysteine-containing ubiquitin was prepared from pET15b-His-Thrombin-Cys-ubiquitin(1–76) (Amp^R^). The plasmid was transfected into BL21 (DE3) E.coli, and an overnight preculture thereof was used to inoculate (1:20) LB medium supplemented with 100 µg/ml ampicillin. Cultures were grown at 37 °C until the OD_600_reached 0.6–0.8, followed by induction with 0.75 mM IPTG at 18 °C overnight. Cells were harvested by centrifugation (4,000g, 10 min at 4 °C), resuspended in lysis Buffer (HEPES 50 mM pH 8, NaCl 250 mM, Glycerol 10 % (w/v), DTT 1 mM, PMSF 0.1 mM, Lysozyme 25 mM, and 1x cOmplete™ Protease Inhibitor Cocktail tablets Roche) and lysed by sonication at 4°C (2 s on / 2 s off, total 5 min, 35% amplitude). The lysate was cleared by centrifugation at 100,000 g for 1 h at 4 °C and the supernatant was filtered (0.45 µm) and loaded ÄKTA FPLC system equipped with a HisTrap HP 5 ml column. The column was equilibrated with binding buffer (50 mM HEPES pH 8, 150 mM NaCl, 20 mM Imidazole, 1 mM DTT), and the lysate was applied. After washing, proteins were eluted with a gradient of 0– 100% elution buffer (50 mM HEPES pH 8, 150 mM NaCl, 300 mM Imidazole, 1 mM TCEP). Protein purity was assessed by SDS-PAGE and Coomassie blue staining. Fractions containing the protein of interest were pooled and concentrated on Amicon Ultra centrifugal filters (4ml, MWCO 3kDa). The protein was further purified via RP-HPLC on Agilent Zorbax 300SB-C18 semi-preparative column (5 µm, 9.4x 250 mm). Fractions were collected and analyzed using direct injection on XEVO G2-XS Q-TOF MS. Purified fractions were lyophilized, argon-flushed, sealed, and stored at −20 °C for long-term preservation.

#### Ubiquitin thrombin cleavage

His-Thrombin-Cys-ubiquitin (Ub-Cys), obtained after HPLC purification, was dissolved in dimethylformamide (DMF) before adding PBS pH 7.4. The reaction mixture was supplemented with 10mM CaCl_2_ and 300mM NaCl, followed by thrombin addition (Sigma (1 KU), 5 units per 100 µg of Ub). The mixture was incubated at 37°C in a thermocycler, and reaction progress was monitored on RP-HPLC analysis on Agilent Zorbax 300SB-C18 column (5 µm, 4.6 x 150 mm) and ESI MS. At this stage the cysteine ubiquitin was labeled according to the previously described maleimide labeling protocol and purified over Agilent Zorbax 300SB-C18, lyophilized and before use refolded in PBS pH 7.4 (3x 1l) through gradual dialysis.

#### Recombinant E2 (UbcH5c) purification

The preculture of pET15b-His-TEV-UbcH5c E2 (Amp^R^) in BL21 (DE3) E.coli was used to inoculate (1:20) 2xYT medium supplemented with 100 µg/ml ampicillin. Cultures were grown at 37 °C until the OD_600_ reached 0.6–0.8, followed by induction with 0.75 mM IPTG at 18 °C overnight. Cells were harvested by centrifugation (4,000g, 10 min at 4 °C), resuspended in 50 mM HEPES pH 8, 250 mM NaCl, 10 % glycerol, 100 µg/l lysozyme, 0.1 mM PMSF, 1x cOmplete™ Protease Inhibitor Cocktail tablets Roche and lysed by sonication at 4°C (1 s on/3 s off, 30–40% amplitude, total 4 min). The lysate was cleared by centrifugation at 100,000 g for 1 h at 4 °C, and the supernatant was filtered (0.45 µm). The clarified lysate was loaded on ÄKTA FPLC system equipped with a HisTrap HP column. The column was equilibrated with binding buffer (50 mM HEPES pH 8, 250 mM NaCl, 20 mM imidazole), and the lysate was applied. After washing, proteins were eluted with a gradient of 0–100% elution buffer (50 mM HEPES pH 8, 250 mM NaCl,400 mM imidazole). Protein purity was assessed by SDS-PAGE and Coomassie blue staining. Fractions containing the protein of interest were pooled and concentrated on Amicon Ultra centrifugal filters (4ml, MWCO 10kDa). Concentrated His-TEV-UbcH5c was dialyzed against TEV cleavage buffer (50 mM Tris-HCl pH 8, 100 mM NaCl, 1 mM DTT) and cleaved with 0.2 equivalent TEV-protease to remove the His-tag forming GGS-UbcH5c. The reaction was analyzed by SDS-PAGE to confirm cleavage efficiency. The cleaved protein was subjected to reverse IMAC (HisTrap HP column, in binding and elution buffer) to remove the His-tag and uncleaved protein. The flow-through containing the target protein was further purified by size exclusion chromatography on Superdex 75 10/300 GL column equilibrated with SEC buffer (50 mM HEPES pH 7.5, 150 mM NaCl, 1 mM TCEP). The fractions analyzed by SDS-PAGE and LC-MS corresponding to the correct mass were concentrated on Amicon Ultra centrifugal filters (4ml, MWCO 10kDa) to 100uM and stored in 50 mM HEPES pH 7.5, 150 mM NaCl, 1 mM TCEP supplemented with 10 % Glycerol at −80 °C.

#### E2∼Ub preloading

The GGS-UbcH5c was dissolved in PBS and treated with 1 equivalent of buffered tris-[2-carboxyethyl]phosphine (TCEP) at 7-7.5 at 25°C and 300 rpm for 20 min to reduce the cysteine residues. Subsequently, a 2-5 times excess of ubiquitin (Ub) or JF549 conjugated ubiquitin (Ub-JF549) was enzymatically loaded onto GGS-UbcH5c using human UBE1 (E1, E-304-050, Bio-techne) in 1x UB buffer (30 mM HEPES, 5 mM MgCl_2_, 10 mM Trisodium Citrate, 2 mM ATP, 10 mM Creatine phosphate tetrahydrate, ddH_2_O, 0.2 µg/ml Creatine phosphokinase) at 37°C for 1h. The reaction was monitored by SDS-PAGE and LC-MS. The reaction mixture was concentrated using Amicon Ultra centrifugal filters (500ul, MWCO 10kDa) and analyzed via LC-MS (XEVO G2-XS Q-ToF) to confirm the ubiquitylation. The preloaded E2∼Ub was further purified by size exclusion chromatography using a Superdex 75 10/300 GL column equilibrated with SEC buffer (50 mM HEPES pH 7.5, 150 mM NaCl, 200 µM TCEP). The E2∼Ub fractions were concentrated in Amicon Ultra centrifugal filters (4ml, MWCO 10kDa) to approximately 300 µl and analyzed by RP-HPLC, MS, and quantified via UV-Vis. The aliquots were flash-frozen and stored at −80 °C.

### General biochemical assays

#### Ensemble ubiquitylation reactions

In a typical reaction setup, 250nM nucleosomes (500nM H2A), 5 uM ubiquitin (Boston Biochem), 25nM E1 enzyme (UBE1, Boston Biochem), and 125nM E2 enzyme (UbcH5c, UBE2DE3, Boston Biochem or expressed) and 62.5 nM variant PRC1 were combined in 1× UB buffer (30 mM HEPES (pH 7.5), 5 mM MgCl_2_, 0.2 mM DTT, 10 mM trisodium citrate, 2 mM ATP, 10 mM creatine phosphate tetrahydrate, and 0.2 µg/ml creatine phosphokinase), ubiquitin was added last, and the mixture was incubated at 30 °C or 37 °C. At the selected time intervals, 10ul samples were taken, and the reaction was quenched by adding 5 µl of 4x Laemmli Sample Buffer (BioRad) containing 10% β-ME. The ubiquitin transfer was analyzed by SDS-PAGE with Coomassie blue staining or via in-gel fluorescence if H2A or ubiquitin were fluorescently labeled. The ubiquitylation progress was quantified via Fiji software (ImageJ), specifically employing the “Gel Analysis” plugin.

#### Ubiquitylation Reaction with E2∼Ub

The reaction was performed as the ubiquitylation reaction but in absence of E1 and any thioesters or reducing agents in 1×UB buffer (30 mM HEPES (pH 7.5), 5 mM MgCl_2_, 0.2 mM DTT, 10 mM trisodium citrate, 2 mM ATP, 10 mM creatine phosphate tetrahydrate, and 0.2 µg/ml creatine phosphokinase) and 1×IB (2.50 mM Trolox, 3.2 % Glucose, 30 mM HEPES, 10 mM Imidazole, 5 mM MgCl_2_, freshly supplemented with 50 mM KCl, 0.005 % Tween 20).

#### Electrophoretic mobility shift assay (EMSA)

For the EMSA, 20 nmol of nucleosomes (wt octamers, 15bp-’601’-15bp-Alexa Fluor 647) were incubated in 1× EMSA buffer (25 mM HEPES pH 7.9, 50 mM NaCl, 5 mM MgCl_2_, 0.05% Tween, 4% glycerol) in a 10 µl reaction volume with increasing amounts of variant PRC1 (0-1500nM). The mixture was incubated for 5min on ice, 10 min at 25 °C and chilled on ice for 5min. Sucrose was added to achieve a final concentration of 8% and samples were loaded onto a ice-cold 5% Criterion TBE gel (Bio-Rad). Gel electrophoresis was performed in ice-cold 0.5× TBE buffer (4 °C) and images were captured on a ChemiDoc MP™Imaging System (BioRad). Quantification was performed using ImageLab (Bio-Rad) software and ImageJ (Fiji) to quantify bound and unbound fractions.

### Microscopy slide preparation

#### Slide cleaning procedure

Glass slides (76 mm x 26 mm) with 16 pre-drilled holes and coverslips (24 mm x 40 mm) were thoroughly cleaned. Initially, the slides and coverslips were sonicated in 10% Alconox detergent solution for 20 min, followed by thorough rinsing in Milli-Q water. This cleaning step was repeated with acetone and ethanol, interspersed with additional rinses in Milli-Q water. Post cleaning, the slides were subjected to a Piranha solution (a 3:1 mixture of 3 M sulfuric acid and 60% hydrogen peroxide) for a minimum of 2 h, and then extensively washed with Milli-Q water, followed by a final sonication in acetone for 10 min. Following the cleaning process, the slides and coverslips were incubated in a 2% (3-Aminopropyl)triethoxysilane (APTES) solution in acetone for 5 min. This was followed by quenching in Milli-Q water and drying under a nitrogen stream.

#### Preparation of flow channels

Double-sided tape was applied to silanized slides to form channels aligned with the drilled holes, which were then covered by silanized coverslips. The open ends of the channels were sealed with epoxy glue. The assembled silanized chambers were stored in vacuum-sealed falcon tubes at −20 °C until use. Before an experiment, pipet tips were glued into the prepared holes using Epoxy glue, allowing for 30 min setting time at room temperature (RT). The channels were then subjected to a PEGylation process. A PEG solution was prepared by dissolving 1 mg of biotinylated PEG-bis-Succinimidyl Valerate (SVA) (5 kDa) and 40 mg of methoxy PEG-SVA (5 kDa) in 320 µl of 0.1 M sodium tetraborate buffer (pH 8.5). This solution was injected into the pre-formed channels (40 µl per channel) and incubated for 3 h at RT. Following this, PEG-SVA (1 kDa) was injected (50 µl per channel) and incubated for an additional 1 h at RT. Ultra-pure water (ROMIL) and T50 buffer (Tris HCl 100 mM pH 7.5, NaCl 500 mM) were degassed before use. A stock solution of 100 mg/ml bovine serum albumin (BSA) in ultra-pure water was prepared fresh, along with 10 % and 1 % Tween-20 solutions in 1X T50 buffer. A 1 M KCl solution was prepared for TetraSpeck beads immobilization. The oxygen-scavenging system (Gloxy) was prepared by dissolving 1 mg catalase in 100 µl of 50 mM phosphate buffer (pH 7.0) and 10 mg glucose oxidase in a mixture of 40 µl catalase solution and 60 µl 1X T50. This mixture was centrifuged at 21’000 g for 3 min, and the supernatant was stored at 4 °C for up to one week. Nanoparticles were diluted in degassed 1X T50 for use. Chromatin samples were spun down at 21’000 g for 10 min at 4 °C to clear aggregates, and the supernatant was transferred to a fresh low-binding tube for concentration measurement. Neutravidin 0.5 mg/ml stocks were prepared in 1xT50, flash frozen and used within one month.

### Single-molecule imaging

#### Microscope setup

Image acquisition was performed on a micro-mirror total internal reflection fluorescence (mmTIRF) microscopy setup (*48*, *63*) (**Fig. S6**), built around the RM21™ microscopy stage and a Nano-LPS nanopositioning stage (MadCityLabs). Laser excitation was achieved via Obis lasers (Coherent), emitting at 405 nm, 488 nm, 532 nm, and 640 nm. Excitation and detection was performed through a 60x NA 1.49 Nikon CFI Apochromat TIRF oil immersion objective (Nikon). In the emission path, images were spectrally separated using a MadView™ image splitter (MadCityLabs), and recorded using an iXon Ultra 897 EMCCD camera (Andor Technology), at a magnification of 100x or a pixel size of 160 nm. The camera was operated by custom-made LabView (National Instruments) software at 17 MHz readout and 900 EM gain. Videos were recorded under <u>al</u>ternating <u>ex</u>citation (ALEX) using the 488 nm, 532 nm and 640 nm laser lines.

#### General preparation of channels for imaging

Flow channels for all smTIRF experiments were prepared by sequentially washing with 500 µL of degassed ultrapure water (ROMIL) and 500 µL of 1x T50 buffer (10 mM Tris pH 7.5, 50 mM NaCl). 50 µL of 0.25 mg/ml neutravidin was injected into the channels, allowed to incubate for 10 min, and then flushed with 500 µL 1x T50. Imaging was performed in Imaging buffer (1xIB: 2.50 mM Trolox, 3.2% Glucose, 30 mM HEPES, 10 mM Imidazole, 5 mM MgCl_2_, freshly supplemented with 50 mM KCl, 0.005 % Tween 20, 5 mg/ml BSA, 1x Gloxy). A peristaltic pump was used for the injection into the microfluidic flow cell.

#### Imaging variant PRC1 chromatin binding dynamics

Background fluorescence was assessed using: 488 nm, 532 nm and 640 nm laser excitations before each experiment. For imaging alignment of multi-color experiments, nanoparticles (NPs, 100nm Biotin Gold NanoUrchins, Cytodiagnostics) were injected at a dilution of 1:100 in 1x T50, with unbound NPs cleared by a 400 µl 1x T50 wash. Alexa Fluor 488-labeled and biotin-modified 12×601 chromatin was introduced at concentrations ranging from 50 to 200 pM for surface immobilization in T50 buffer (Tris HCl 100 mM pH 7.5, NaCl 50 mM). Surface coverage was monitored via smTIRF imaging with 488 nm excitation. Channels were subsequently rinsed with 500 µl of 1x T50 and 500 µl of imaging buffer (1xIB: 2.50 mM Trolox, 3.2% Glucose, 30 mM HEPES, 10 mM Imidazole, 5 mM MgCl_2_, freshly supplemented with 50 mM KCl, 0.005 % Tween 20, 5 mg/ml BSA, 1x Gloxy). Alexa Fluor 647-labeled PRC1.1 or PRC1.4, diluted to 50 pM to 1000 pM in imaging buffer, was introduced for interaction studies. The flow was maintained using a peristaltic pump for consistent delivery. Imaging was performed using the following illumination protocol:

|  | Laser ton [ms] | Laser toff [ms] | Frame no. [n] | Cycles [n] |
| --- | --- | --- | --- | --- |
| Chromatin | 100 | 5 | 1 | ↓ 80-100 X |
| vPRC1 | 100 | 110 | 100 |  |

#### Imaging variant PRC1 chromatin ubiquitylation dynamics

After assessing clean background in all channels, NPs were immobilized for image alignment. For chromatin immobilization, 50 to 200 pM of Alexa Fluor 488-labeled 12×601 chromatin-biotin was injected. Surface coverage was monitored via smTIRF with 488 nm excitation. Channels were subsequently rinsed with 500 µl of 1x T50 and 500 µl of imaging buffer (1xIB: 2.50 mM Trolox, 3.2% Glucose, 30 mM HEPES, 10 mM Imidazole freshly supplemented with 2 mM MgCl_2_, 50 mM KCl, 0.01 % Tween 20, 2 mg/ml BSA, 1x Gloxy). Alexa Fluor 647-labeled variant PRC1 was diluted to 50 pM to 1000 pM and E2-Ub diluted to 5 NM to 200 nM in 1xIB before injection into the channel at 25 °C using a peristaltic pump. Imaging was performed using the following illumination protocol:

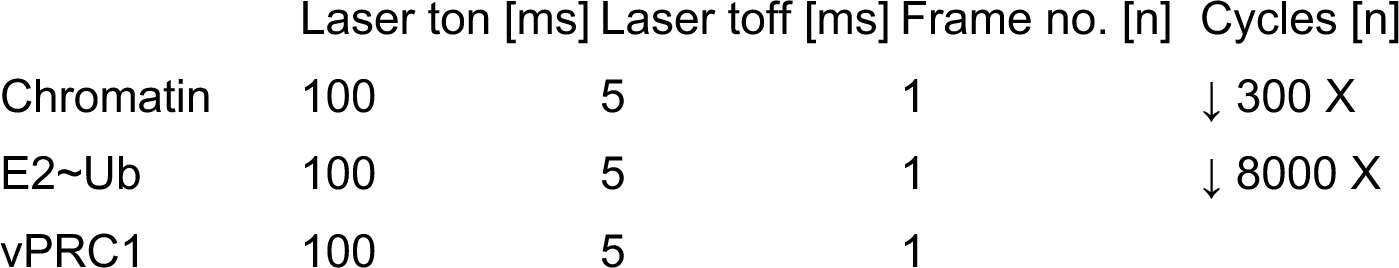

|  | Laser ton [ms] | Laser toff [ms] | Frame no. [n] | Cycles [n] |
| --- | --- | --- | --- | --- |
| Chromatin | 100 | 5 | 1 | ↓ 300 X |
| E2~Ub | 100 | 5 | 1 | ↓ 8000 X |
| vPRC1 | 100 | 5 | 1 |  |

Post-ubiquitylation, the channels were thoroughly washed with 400 µl IB supplemented with 200 mM KCl and 1×Gloxy, to remove nonspecifically bound proteins. A wash movie was recorded during and after the wash, for quality control and to assess the final degree of chromatin ubiquitylation.

### Data processing

#### Single-molecule binding data processing

Data analysis was performed as described previously (*41*). NPs were visible in each emission channel and used for image alignment. Using a macro in Fiji (ImageJ) (*64*), regions of interest (ROIs) were determined and saved, ensuring consistency across all movies. NP movies were then split into individual channels, with the 488 nm channel serving as a Z-projection of all frames and saved as an average. The 532 nm and 640 nm channels were saved as separate movies. Background subtraction was performed using a rolling ball method in ImageJ to correct for uneven illumination and background fluorescence. Subsequently, MATLAB (MathWorks, Natick, MA, USA, release R2022b) was employed for further alignment. The 488 nm AVG image served as the reference, onto which the 532nm and 640nm channels were corrected using a nonlinear image transform to achieve perfect image alignment for all channels and to correct any image drift. Subsequently, chromatin positions were identified via local maxima detection in Matlab. A peak-finding algorithm was employed to detect immobile chromatin fibers, with each diffraction-limited spot analyzed using a 2D-Gaussian fitting algorithm to extract precise coordinates. Co-localization of variant PRC1 with immobilized DNA/chromatin was determined using 2D-Gaussian fitting of detected single-molecule emitters to determine the shape of the point spread function (PSF), excluding peaks wider than the PSF expected for a single fluorophore. Fluorescence intensities were extracted within a 2-pixel radius at a resolution of 160 nm/pixel of the identified DNA peaks for each frame in the stack to obtain fluorescence intensity time traces corresponding to individual chromatin positions. Traces were excluded from analysis when one of the following conditions were met: (1) no emission in the 640 nm ‘PRC1’ channel over the whole time course (i.e. no PRC1 binding), (2) total emission in the 640 nm channel above a set threshold of 5000 counts (local aggregation of PRC1 molecules or binding of multiple PRC1 complexes), (3) constant emission throughout the whole time course (local aggregation of PRC1 molecules), (4) less than 10% of total number of extracted traces showed PRC1 binding.

Within single fluorescence intensity time traces, individual fluorescence detections were excluded from analysis if their corresponding PSF parameters exceeded limits set for single-molecule detections (>300 nm FWHM), and if they did not co-localize with determined chromatin positions within a 250 nm distance threshold. Moreover, overlapping binding events were excluded. The resulting fluorescence intensity time traces were analyzed using a step detection algorithm (*65*) followed by thresholding to extract binding (t_bright) and unbound times (t_dark). The obtained binding times and unbound times were collected into cumulative histograms. Residence times were determined using a semi-automatic approach. Individual binding events were detected with thresholding, excluding overlapping events. Cumulative histograms of bright times were fitted with bi-exponential functions to obtain nonspecific (τ_off,1_) and specific τ_off,2_) residence times. Association dark times histograms were fitted with mono-exponential functions to calculate apparent on-rates (k_app). Specific on-rates (k_on) were obtained by correcting for contributions from nonspecific binding events.

#### Single-molecule ubiquitylation data processing

The fluorescence intensity time traces were exported for variant PRC1 binding and ubiquitin transfer as presented before for variant PRC1 binding kinetics. In short, the determined emission ROIs were cut from individual image stacks in FiJi (ImageJ) and aligned by applying the transformation matrices for individual channel alignment in MATLAB. Due to the prevalence of background fluorescence caused by high E2∼Ub concentration of 25 nM to 40 nM, the obtained frames were each grouped into 5-10 average frames to minimize background. From each of the individual chromatin positions, determined using the 488 nm channel, fluorescence intensity time traces were exported in both 532 nm and 640 nm channels, followed by fitting with a step-detection algorithm (*65*).

Traces were excluded from analysis when one of the following conditions were met for the 640 nm ‘PRC1’ channel: (1) no emission in the 640 nm channel over the whole time course (i.e. no PRC1 binding), (2) total emission in the 640 nm channel above a set threshold of 5000 counts (local aggregation of PRC1 molecules), (3) constant emission throughout the whole time course (local aggregation of PRC1 molecules) and (4) chromatin occupancy by PRC1 was lower than 1%. Traces were excluded from analysis when one of the following conditions were met for the 532 nm ‘ubiquitin’ channel: (1) presence of 532 emission which is not colocalizing above 400nm with the chromatin position (surface aggregation), (2) chromatin occupancy by ubiquitin was below 20%. Within 640 and 532 nm single fluorescence intensity time traces, individual fluorescence detections were excluded from analysis if their corresponding PSF parameters exceeded limits set for single-molecule detections, and if they did not co-localize with determined chromatin positions within a 400 nm distance threshold. Moreover, overlapping binding events in the 640nm channel were excluded. Composite kymograph plots were generated from individual traces, while the traces were sorted by the first appearance of stable emission in the ‘ubiquitin’ channel.

Ubiquitin step number was determined by quantifying the fluorescence counts corresponding to a single step in the 532 nm fluorescence intensity traces, as determined by the step-finding function, with a minimal threshold of 50 counts. The obtained step size distribution was plotted as a histogram to obtain the average step size of a single ubiquitin corresponding to a single fluorophore Janelia Fluor 549. The same approach was applied to determine the photobleaching of Janelia Fluor 549 (see **Fig. 2E**).

From individual traces, libraries of ubiquitylation events were generated. PRC1 binding events associated with a step in ubiquitin emission (step size according to the above determined distribution), including additional 10 seconds before and after the event were extracted. Events were further filtered to not exceed a threshold value in the 532 nm channel emission, rendering the steps clearly identifiable. From these event libraries, values for ***τ***_res,act_, ***τ***_ub,1_, ***τ***_ub,2_ were automatically calculated by a custom-made Matlab protocol, and converted into histograms for further analysis.

### Statistical analysis

All statistical analysis and fitting was performed in GraphPad Prism 10 (GraphPad Software, La Jolla, CA, USA, version 10.3.1). The applied statistical methods are indicated in each figure. For multiple comparisons, one-way analysis of variance (ANOVA) with Tukey’s comparison test was applied. A *p* value of *p* < 0.05 was considered significant. The *p* values, number or independent repeats and statistical analysis methods, are described in the figure legends.

## Acknowledgments

We thank Prof. Dr. Titia Sixma for providing initial PRC1.4 constructs, Prof. Dr. Andreas Marx for the ubiquitin expression constructs, Yousuf Khan for help with the preparation of H2AK119ub containing octamers, Dr. Pauline Franz for expression of GST-thrombin-Ub1-75-(G75M>Aha), Dr. Iuliia Boichenko and Mathilde Le Jeune for cloning and purification of H2AK119C histone. We acknowledge Dr. Florence Pojer, Dr. Kelvin Lau, and Dr. Amédé Noredine Larabi (PTPSP, EPFL) for Baculovirus generation and expression of vPRC1 constructs.

## Author contributions

Conceptualization: AT, BF

Methodology: AT

Software: AT, BF

Formal analysis: AT, BF

Investigation: AT

Visualization: AT

Resources: AT

Supervision: BF

Writing—original draft: AT, BF

Writing—review & editing: AT, BF

Funding acquisition: BF

## Additional information

Supplementary information contains supplementary figures and tables with a detailed overview of purification, labeling, chemical synthesis, analytical data, and single-molecule data acquisition for used protein and DNA materials.

