## Supplemental Material for "Single-molecule analysis reveals the mechanism of chromatin ubiquitylation by variant PRC1 complexes"

**This PDF file includes:**

Supplementary Text
Figures S1 to S15
Tables S1 to S3
References (1 to 1)

### Supplementary Text

Protein sequences

All the proteins and complex subunits below were derived from the Homo sapiens (human) organism.

Color code: His-tag, Strep-tag, ybbR-tag, Thrombin site, TEV site, linker, HRV-3C site

**RING1B-ybbR-Strep**

MSQAVQTNGTQPLSKTWELSLYELQRTPQEAITDGLEIVVSPRSLHSELMCPICLDMLKNT
MTTKECLHRFCADCIITALRSGNKECPTCRKKLVSKRSLRPDPNFDALISKIYPSRDEYEAH
QERVLARINKHNNQQALSHSIEEGLKIQAMNRLQRGKKQQIENGSGAEDNGDSSHCSNAS
THSNQEAGPSNKRTKTSDDSGLELDNNNAAMAIDPVMDGASEIELVFRPHPTLMEKDDSA
QTRYIKTSGNATVDHLSKYLAVRLALEELRSKGESNQMNLDTASEKQYTIYIATASGQFTVL
NGSFSLELVSEKYWKVNKPMELYYAPTKEHKGSDSLEFIASKLAGSWSHPQFEK*

**PCGF1**

MASPQGGQIAIAMRLRNQLQSVYKMDPLRNEEEVRVKIKDLNEHIVCCLCAGYFVDATTITE
CLHTFCKSCIVKYLQTSKYCPMCNIKIHETQPLLNLKLDRVMQDIVYKLVPGLQDSEEKRIRE
FYQSRGLDRVTQPTGEEPALSNLGLPFSSFDHSKAHYYRYDEQLNLCLERLSSGKDKNKS
VLQNKYVRCSVRAEVRHLRRVLCHRLMLNPQHVQLLFDNEVLPDHMTMKQIWLSRWFGK
PSPLLLQYSVKEKRR*

**PCGF1-His**

MASPQGGQIAIAMRLRNQLQSVYKMDPLRNEEEVRVKIKDLNEHIVCCLCAGYFVDATTITE
CLHTFCKSCIVKYLQTSKYCPMCNIKIHETQPLLNLKLDRVMQDIVYKLVPGLQDSEEKRIRE
FYQSRGLDRVTQPTGEEPALSNLGLPFSSFDHSKAHYYRYDEQLNLCLERLSSGKDKNKS
VLQNKYVRCSVRAEVRHLRRVLCHRLMLNPQHVQLLFDNEVLPDHMTMKQIWLSRWFGK
PSPLLLQYSVKEKRRGLEVLFQGPSAAHHHHHH*

**His-RYBP**

MAHHHHHHSAALEVLFQGPGMTMGDKKSPTRPKRQAKPAADEGFWDCSVCTFRNSAEA
FKCSICDVRKGTSTRKPRINSQLVAQQVAQQYATPPPPKKEKKEKVEKQDKEKPEKDKEIS
PSVTKKNTNKKTKPKSDILKDPPSEANSIQSANATTKTSETNHTSRPRLKNVDRSTAQQLAV
TVGNVTVIITDFKEKTRSSSTSSSTVTSSAGSEQQNQSSSGSESTDKGSSRSSTPKGDMSA
VNDESF*

**Bmi1 (PCGF4)**

MHRTTRIKITELNPHLMCVLCGGYFIDATTIIECLHSFCKTCIVRYLETSKYCPICDVQVHKTR
PLLNIRSDKTLQDIVYKLVPGLFKNEMKRRRDFYAAHPSADAANGSNEDRGEVADEDKRIIT
DDEIISLSIEFFDQNRLDRKVNKDKEKSKEEVNDKRYLRCPAAMTVMHLRKFLRSKMDIPNT
FQIDVMYEEEPLKDYYTLMDIAYIYTWRRNGPLPLKYRVRPTCKRMKISHQRDGLTNAGELE
SDSGSDKANSPAGGIPSTSSCLPSPSTPVQSPHPQFPHISSTMNGTSNSPSGNHQSSFAN
RPRKSSVNGSSATSSG*

**Bmi1-His**

MHRTTRIKITELNPHLMCVLCGGYFIDATTIIECLHSFCKTCIVRYLETSKYCPICDVQVHKTR
PLLNIRSDKTLQDIVYKLVPGLFKNEMKRRRDFYAAHPSADAANGSNEDRGEVADEDKRIIT
DDEIISLSIEFFDQNRLDRKVNKDKEKSKEEVNDKRYLRCPAAMTVMHLRKFLRSKMDIPNT
FQIDVMYEEEPLKDYYTLMDIAYIYTWRRNGPLPLKYRVRPTCKRMKISHQRDGLTNAGELE
SDSGSDKANSPAGGIPSTSSCLPSPSTPVQSPHPQFPHISSTMNGTSNSPSGNHQSSFAN
RPRKSSVNGSSATSSGGLEVLFQGPSAAHHHHHH*

**His-thrombin-GlySerHisCys-Ub**

MGSSHHHHHHSSGLVPRGSHCMQIFVKTLTGKTITLEVEPSDTIENVKAKIQDKEGIPPDQQ
RLIFAGKQLEDGRTLSDYNIQKESTLHLVLRLRGG*

**GlySerHisCys-JF549-Ub**
GSHC(JF549)MQIFVKTLTGKTITLEVEPSDTIENVKAKIQDKEGIPPDQQRLIFAGKQLEDGR
TLSDYNIQKESTLHLVLRLRGG*

**Ub-G75M (Ub-N3)**
GSRRASVGSQIFVKTLTGKTITLEVEPSDTIENVKAKIQDKEGIPPDQQRLIFAGKQLEDGRT
LSDYNIQKESTLHLVLRLRM(Aha)

**His-TEV-E2**
MGSSHHHHHHSSGENLYFQGGSMALKRINKELSDLARDPPAQCSAGPVGDDMFHWQATI
MGPNDSPYQGGVFFLTIHFPTDYPFKPPKVAFTTRIYHPNINSNGSICLDILRSQWSPALTIS
KVLLSICSLLCDPNPDDPLVPEIARIYKTDRDKYNRISREWTQKYAM*

**GGS-E2**
GGSMALKRINKELSDLARDPPAQCSAGPVGDDMFHWQATIMGPNDSPYQGGVFFLTIHFP
TDYPFKPPKVAFTTRIYHPNINSNGSICLDILRSQWSPALTISKVLLSICSLLCDPNPDDPLVP
EIARIYKTDRDKYNRISREWTQKYAM*

**H2A K119C**
MSGRGKQGGKARAKAKSRSSRAGLQFPVGRVHRLLRKGNYAERVGAGAPVYMAAVLEYL
TAEILELAGNAARDNKKTRIIPRHLQLAIRNDEELNKLLGKVTIAQGGVLPNIQAVLLPCKTES
HHKAKGK*

**H2A K119C -Propagyl acrylate (PA)**
MSGRGKQGGKARAKAKSRSSRAGLQFPVGRVHRLLRKGNYAERVGAGAPVYMAAVLEYL
TAEILELAGNAARDNKKTRIIPRHLQLAIRNDEELNKLLGKVTIAQGGVLPNIQAVLLPC(PA)K
TESHHKAKGK*

**H2A K119C -PA- Ub**
MSGRGKQGGKARAKAKSRSSRAGLQFPVGRVHRLLRKGNYAERVGAGAPVYMAAVLEYL
TAEILELAGNAARDNKKTRIIPRHLQLAIRNDEELNKLLGKVTIAQGGVLPNIQAVLLPC(PA-
Ub)KTESHHKAKGK*

**wild type H2A**
MSGRGKQGGKARAKAKSRSSRAGLQFPVGRVHRLLRKGNYAERVGAGAPVYMAAVLEYL
TAEILELAGNAARDNKKTRIIPRHLQLAIRNDEELNKLLGKVTIAQGGVLPNIQAVLLPKKTES
HHKAKGK*

**H2A N110C**
MSGRGKQGGKARAKAKSRSSRAGLQFPVGRVHRLLRKGNYAERVGAGAPVYMAAVLEYL
TAEILELAGNAARDNKKTRIIPRHLQLAIRNDEELNKLLGKVTIAQGGVLPCIQAVLLPKKTES
HHKAKGK*

**H2A N110C-Atto488**
MSGRGKQGGKARAKAKSRSSRAGLQFPVGRVHRLLRKGNYAERVGAGAPVYMAAVLEYL
TAEILELAGNAARDNKKTRIIPRHLQLAIRNDEELNKLLGKVTIAQGGVLPC(Atto488)IQAVLL
PKKTESHHKAKGK*

**wild type H2B**
MPEPAKSAPAPKKGSKKAVTKAQKKDGKKRKRSRKESYSVYVYKVLKQVHPDTGISSKAM
GIMNSFVNDIFERIAGEASRLAHYNKRSTITSREIQTAVRLLLPGELAKHAVSEGTKAVTKYT
SAK*

**H3.2 C110A**
MARTKQTARKSTGGKAPRKQLATKAARKSAPATGGVKKPHRYRPGTVALREIRRYQKSTE
LLIRKLPFQRLVREIAQDFKTDLRFQSSAVMALQEASEAYLVGLFEDTNLAAIHAKRVTIMPK
DIQLARRIRGERA*

**wild type H4**
MSGRGKGGKGLGKGGAKRHRKVLRDNIQGITKPAIRRLARRGGVKRISGLIYEETRGVLKV
FLENVIRDAVTYTEHAKRKTVTAMDVVYALKRQGRTLYGFGG*

**H2A K119R/K120R**
MSGRGKQGGKARAKAKSRSSRAGLQFPVGRVHRLLRKGNYAERVGAGAPVYMAAVLEYL
TAEILELAGNAARDNKKTRIIPRHLQLAIRNDEELNKLLGKVTIAQGGVLPNIQAVLLPRRTES
HHKAKGK*

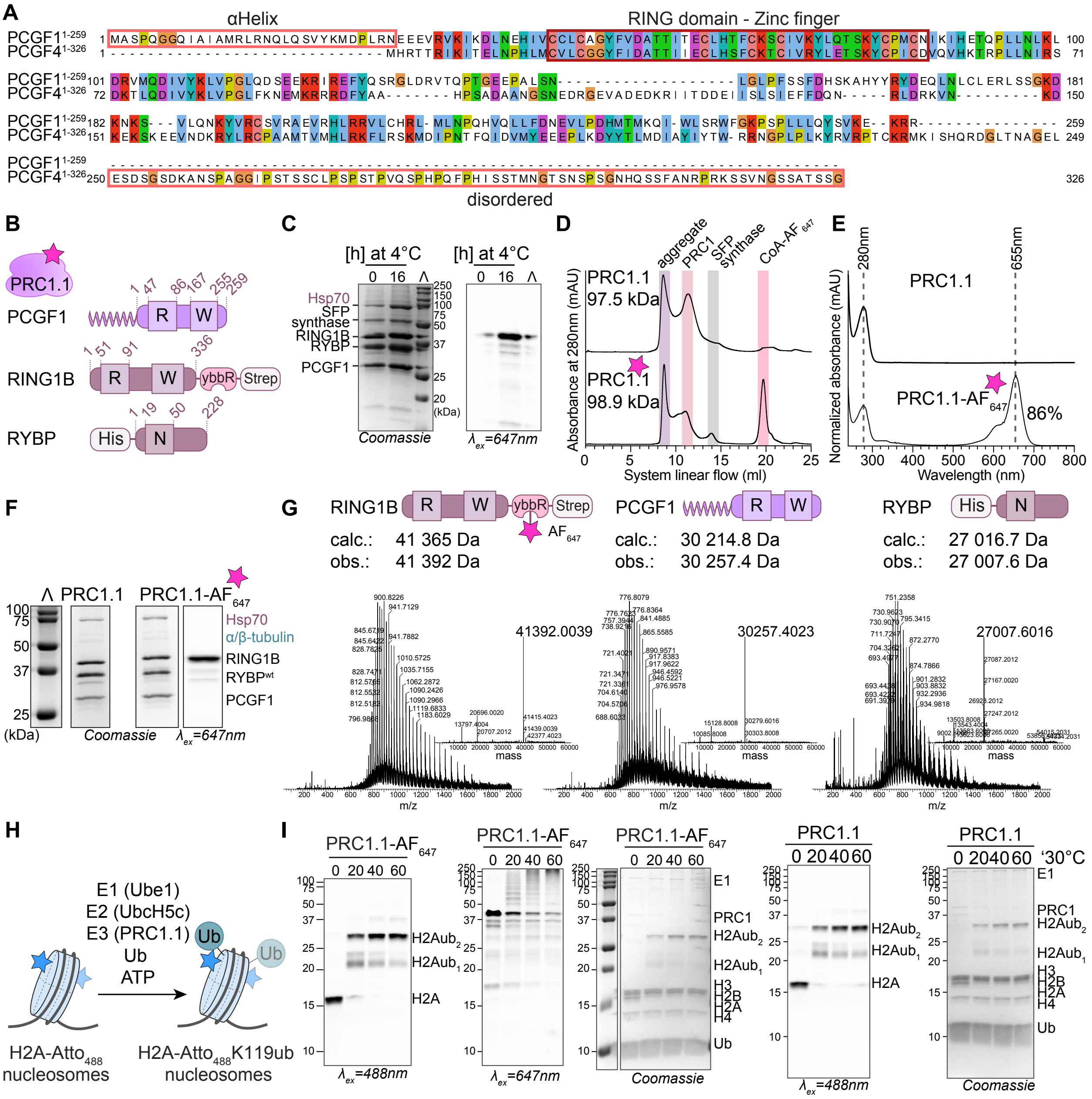

**Fig. S1. Purification of PRC1.1. A)** Sequence alignment of human PCGF1 and PCGF4. The RING domain is marked in red, the N-terminal helix of PCGF1 and the C-terminal disordered domain of PCGF4 are indicated in orange. **B**) PRC1.1 complex subunits and domains. ybbR-, His- and Strep-tag are indicated. Numbers correspond to amino acid residues. R: RING domain W: RAWUL domain. **C**) SDS-PAGE analysis of ybbR-tag labeling reaction of PRC1.1 with Alexa Fluor 647 ($AF_{647}$) C2 maleimide detected via in-gel fluorescence, excitation at $\lambda_{ex}$=647nm (PRC1.1-$AF_{647}$), and Coomassie blue. **D**) SEC profiles, **E**) UV-Vis absorbance spectra and **F**) SDS-PAGE analysis (Coomassie and in-gel fluorescence $\lambda_{ex}$ = 647 nm) of unlabeled PRC1.1 and labeled PRC1.1-$AF_{647}$. **G**) ESI-MS m/z and deconvolved mass spectra of: RING1B-ybbR-$AF_{647}$-Strep (calc.: 41365 Da, obs.: 41392 Da), PCGF1 (calc.: 30214.8 Da, obs.: 30257.4 Da) and His-RYBP (calc.: 27016.7 Da, obs.: 27007.6 Da). Calculated mass (calc.), observed mass (obs.). **H**) Ubiquitylation reaction scheme with indicated components, in particular using Atto 488 ($Atto_{488}$)-labeled H2A for detection. **I**) Ubiquitylation reactions using labeled PRC1.1-$AF_{647}$ and unlabeled PRC1.1, detected via in-gel fluorescence, excitation at $\lambda_{ex}$=488nm (H2A-$Atto_{488}$) and $\lambda_{ex}$=647nm (PRC1.1-$AF_{647}$), and Coomassie blue.

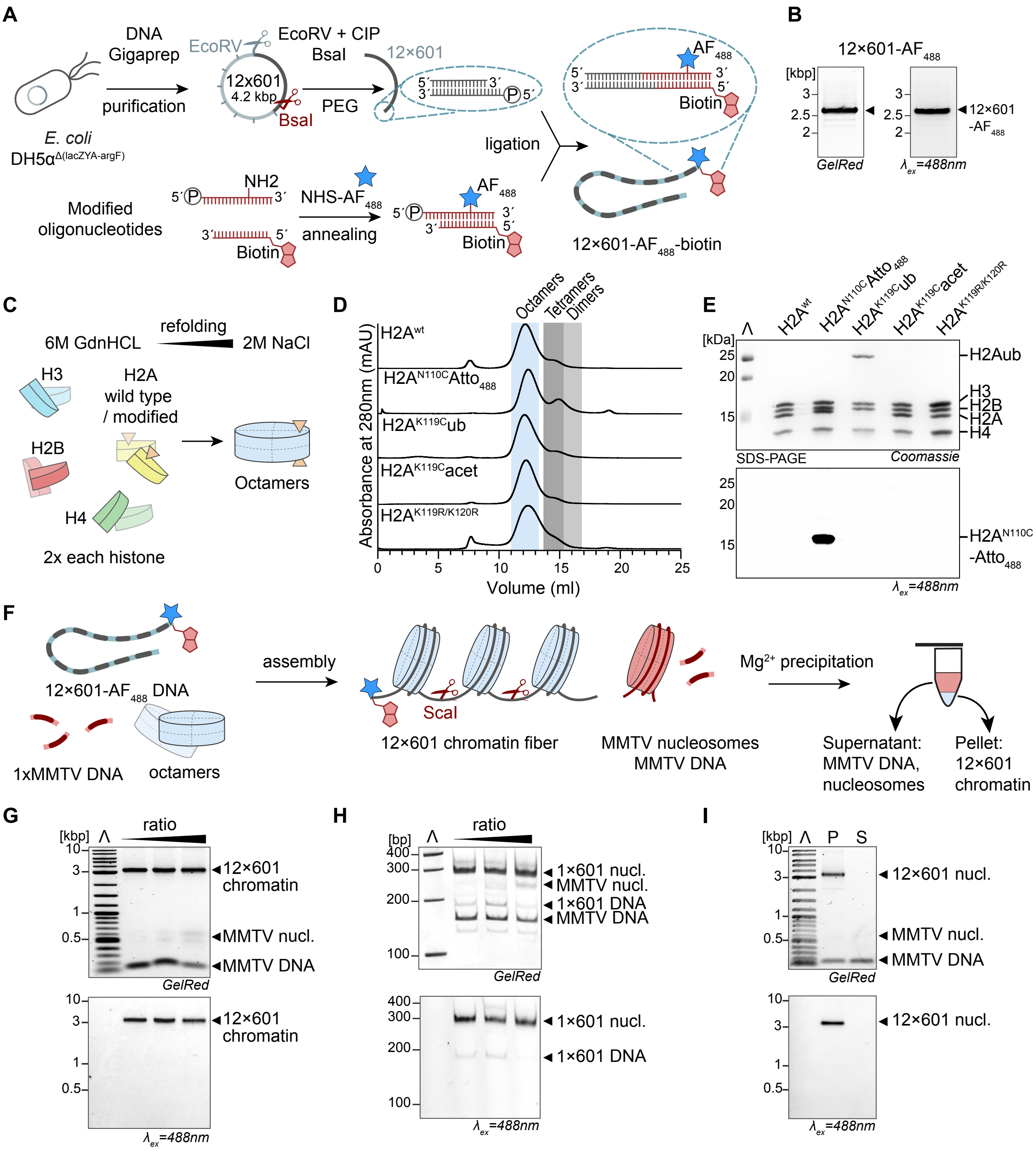


**Fig. S2. Chromatin preparation. A**) Top: 12×601 DNA obtained from a plasmid after EcoRV digest, dephosphorylation, and polyethyleneglycol (PEG) precipitation. BsaI digest yields 12×601 DNA with a 4 bp 5'-phosphate overhang for the subsequent ligation. Bottom: chromatin anchor design. Sense oligonucleotide contains a 5'-phosphate group and 4 bp overhang for the ligation, as well as an internal amine (C6 thymidine) for fluorescent labeling with Alexa Fluor 488 (AF$_{488}$) NHS ester. Antisense oligonucleotide is designed with a complementary sequence and 5'-biotin anchor. After the labeling reaction and HPLC purification, both primers are annealed. Ligation of chromatin DNA to the annealed oligonucleotide yields labeled 12×601-AF$_{488}$-biotin DNA. **B**) Final purification to pure 12×601-AF$_{488}$-biotin DNA, detected via in-gel fluorescence, excitation at $\lambda_{ex}$=488nm (12×601-AF$_{488}$-biotin), and GelRed: (DNA intercalating stain). **C**) Scheme for histone octamer assembly via dialysis from high to low salt. **D**) SEC purification profiles of histone octamers. The H2A modification is indicated in each line, while the other histones did not change:

$H2B^{wt}$, $H3.2^{C110A}$ and $H4^{wt}$. **E**) SDS-PAGE analysis of assembled octamers after SEC purification, modified H2A is indicated as in D). **F**) Assembly scheme of chromatin fibers with 12×601-$AF_{488}$-biotin DNA and histone octamers, in the presence of MMTV buffer DNA to avoid octane overload, followed by purification via magnesium precipitation. **G**) Agarose gel analysis of 12×601-$AF_{488}$-biotin chromatin fibers with increased octamer:DNA ratio. **H**) Native-PAGE analysis of ScaI digested 12×601-$AF_{488}$-biotin chromatin. 1×601/MMTV nucleosomes and DNA observed. **I**) Agarose gel analysis of magnesium precipitated 12-mer chromatin fibers. Input (In), pellet (P), supernatant (S). In-gel fluorescence, excitation at $\lambda_{ex}$=488nm (12×601-$AF_{488}$-biotin), and GelRed: (DNA intercalating stain).

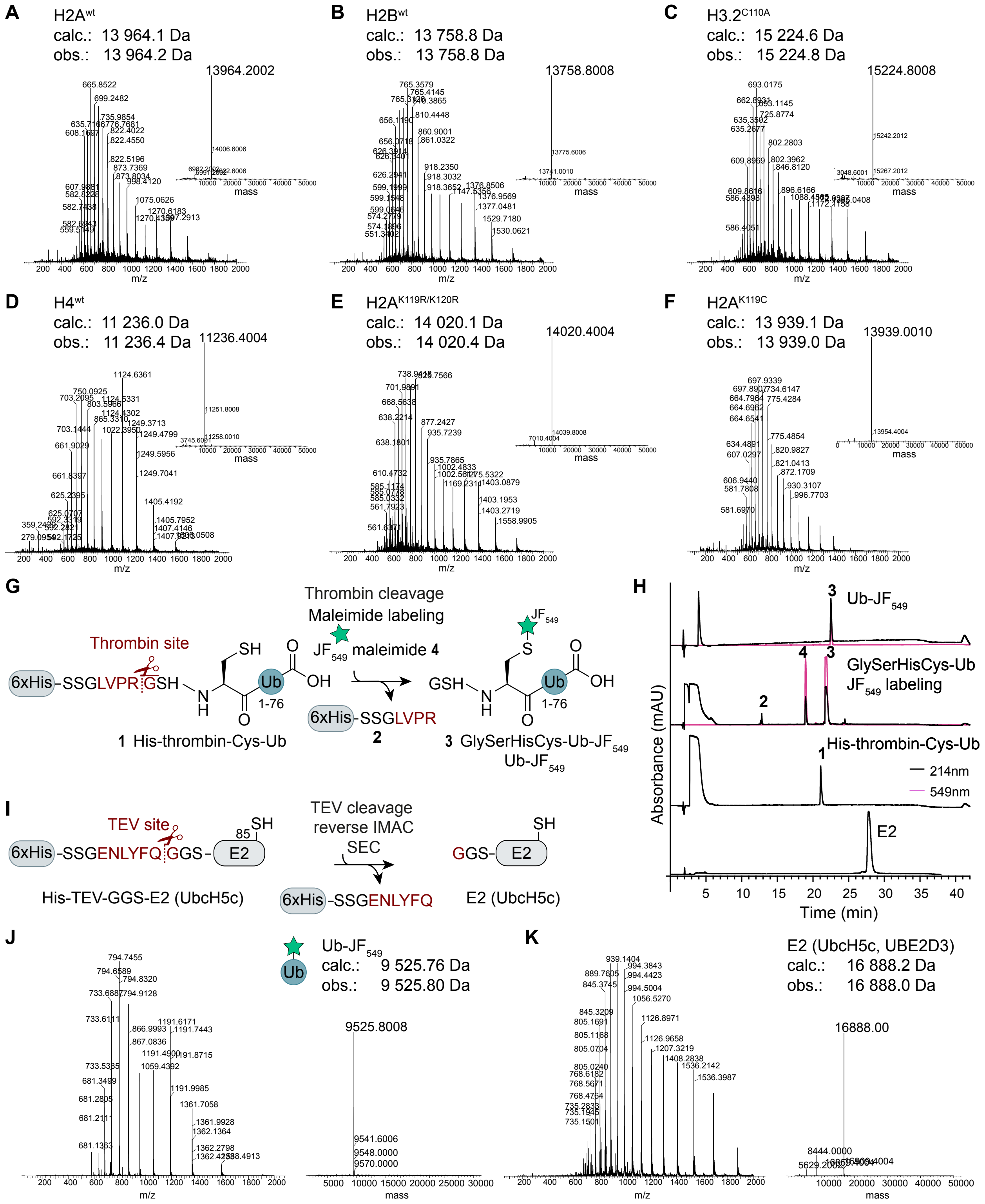


**Fig. S3. Preparation of histones, fluorescent ubiquitin (Ub-JF549) and E2.** ESI-MS m/z and deconvolved mass spectra of the following human histones: **A)** H2A$^{wt}$ (calc.: 13964.1 Da, obs.: 13964.2 Da), **B)** H2B$^{wt}$ (calc.: 13758.8 Da, obs.: 13758.8 Da), **C)** H3.2$^{C110A}$ (calc.: 15224.6 Da, obs.: 15224.8 Da), **D)** H4$^{wt}$ (calc.: 11236.0 Da, obs.: 11236.4 Da), **E)** H2A$^{K119R/K120R}$ (calc.: 14020.1 Da, obs.: 14020.4 Da), **F)** H2A$^{K119C}$ (calc.: 13939.1 Da, obs.: 13939.0 Da). Calculated mass (calc.), observed mass (obs.). **G**) Scheme of the preparation of labeled ubiquitin. After the expression of 6xHis-thrombin-Cys-Ub and IMAC purification, the 6xHis-tag is removed via thrombin cleavage reaction, followed by labeling with Janelia Fluor 549 ($JF_{549}$) maleimide. **H**) Analytical RP-HPLC of thrombin cleavage, $JF_{549}$ labeling and E2. **1:** 6xHis-thrombin-Cys-Ub, **2:** 6xHis-SSGLVPR, **3:** Ub-$JF_{549}$ (GlySerHisCys-$JF_{549}$-Ub), **4:** $JF_{549}$-maleimide. **I)** Scheme of the TEV cleavage reaction of 6xHis-TEV-GGS-E2 yields E2 (UbcH5c) with a short three amino acids long scar (GGS-E2). **J**) ESI-MS m/z and deconvolved mass spectrum of Ub-$JF_{549}$ (calc.: 9525.76 Da, obs.: 9525.80 Da) and **K**) of E2 (UbcH5c, UBE2D3) (calc.: 16888.2 Da, obs.: 16888.0 Da).

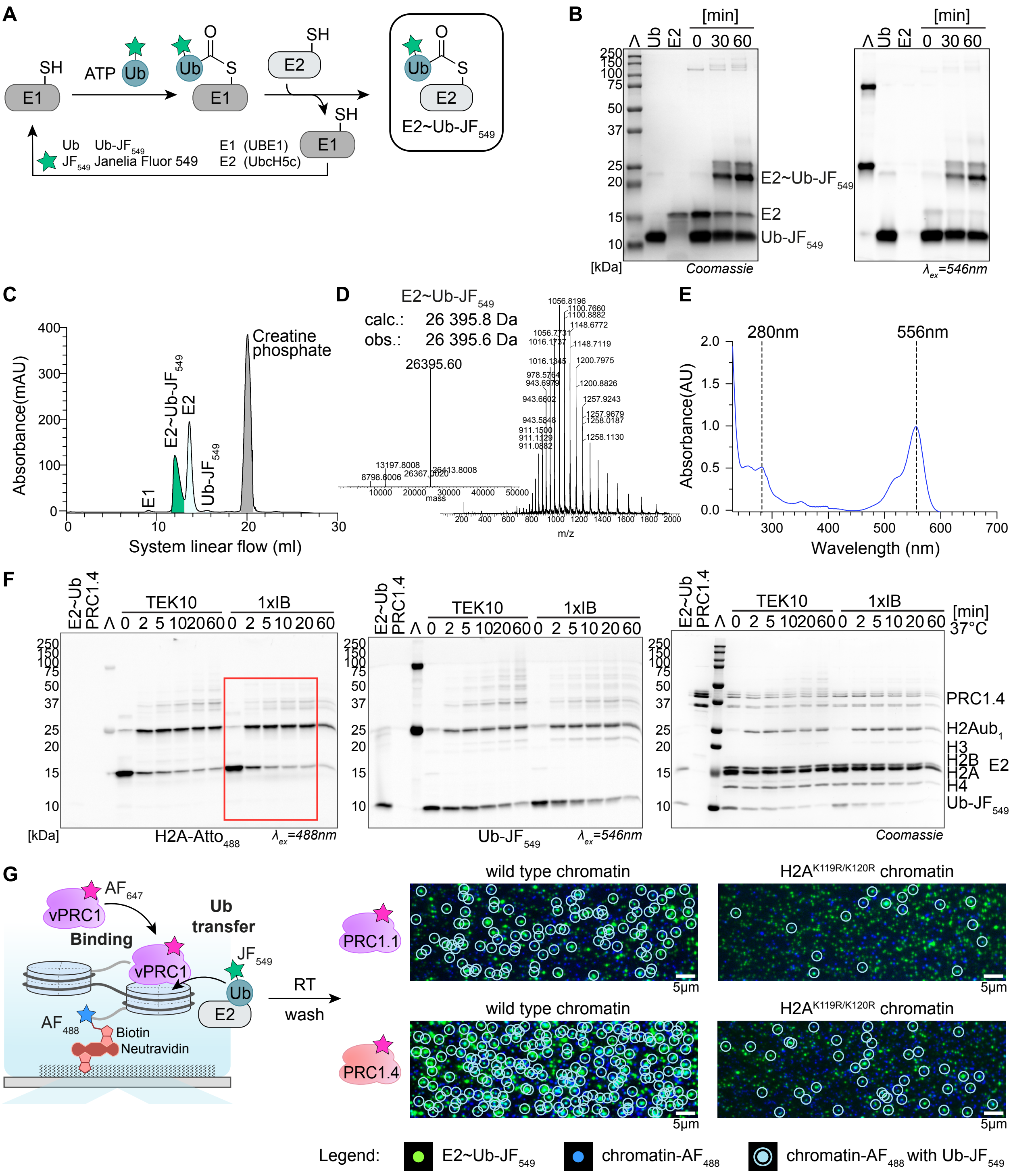

**Fig. S4. Preloaded E2~Ub induces ubiquitylation in the presence of PRC1. A**) Scheme of enzymatic preloading of E2~Ub labeled with $JF_{549}$. Ub-$JF_{549}$: fluorescent ubiquitin, E1: ubiquitin-activating enzyme, here UBE1, E2: ubiquitin-conjugating enzyme, UbcH5c. **B**) SDS-PAGE analysis of enzymatic preloading of E2 with Ub-$JF_{549}$. After 1 h the E2~Ub-$JF_{549}$ adduct is formed, detected via Coomassie blue and in-gel fluorescence, excitation at $\lambda_{ex}$=546nm (Ub-$JF_{549}$). **C**) SEC (Superdex 75 10/300 GL) purification profile of E2~Ub-$JF_{549}$. E1, E2, Ub-$JF_{549}$ and creatine phosphate elute separately. **D**) ESI-MS m/z and deconvolved mass spectrum of E2~Ub-$JF_{549}$ (calc.: 26395.8 Da, obs.: 26395.6 Da). **E**) UV-Vis absorbance spectrum of the formed E2~Ub-$JF_{549}$. **F**) Comparison of ubiquitylation efficiency between imaging buffer (1xIB) and TEK10 buffer on 12-mer chromatin arrays. SDS-PAGE analysis of the ubiquitylation assay with preloaded E2~Ub-$JF_{549}$ and PRC1.4 in the absence of E1 and ATP. The red frame shows the image presented in **Fig. 1F**. In-gel fluorescence, excitation at $\lambda_{ex}$=488nm (H2A$^{N110C}$-$Atto_{488}$) and $\lambda_{ex}$=546nm (Ub-$JF_{549}$) and Coomassie blue. Most of the H2A is ubiquitylated after 5 min incubation. **G**) Colocalization of ubiquitin on chromatin after single-molecule ubiquitylation. Left: scheme of ubiquitylation of H2A$^{wt}$ or H2A$^{K119R/K120R}$ chromatin fibers, using labeled PRC1-$AF_{647}$ complexes. Right; ubiquitylation imaged after 20 min incubation at room temperature (RT) followed by washing the channel. Legend at the bottom: chromatin is in blue, E2~Ub and Ub in green and ubiquitylated chromatin in cyan. Scale bar: 5µm.

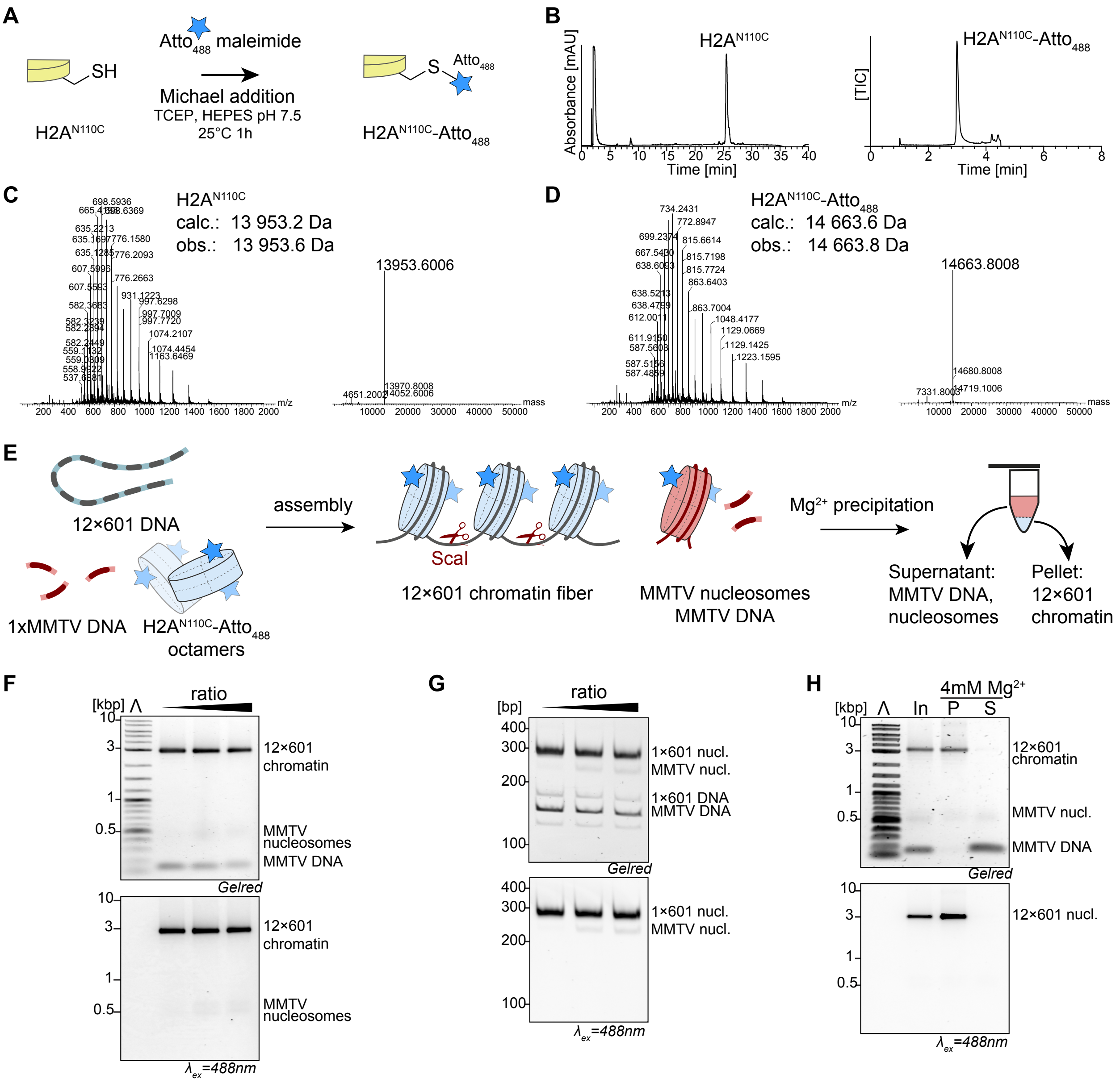


**Fig. S5. Preparation of H2A-Atto$_{488}$ containing chromatin. A)** Reaction scheme for fluorescent labeling of H2A$^{N110C}$ with Atto 488 (Atto$_{488}$) maleimide. **B)** Analytical RP-HPLC analysis of purified H2A$^{N110C}$ (left panel) and LC analysis of purified H2A$^{N110C}$-Atto$_{488}$ (right panel). TIC: total ion count. **C)** ESI-MS m/z and deconvolved mass spectra of H2A$^{N110C}$ (calc.: 13953.2 Da, obs.: 13953.6 and **D**) H2A$^{N110C}$-Atto$_{488}$ (calc.: 14663.6 Da, obs.: 14663.8) . Calculated mass (calc.), observed mass (obs.). **E**) Assembly scheme of 12-mer chromatin fibers using 12×601 DNA and H2A$^{N110C}$-Atto$_{488}$ containing octamers, in the presence of MMTV buffer DNA to avoid histone octamer overload, followed by purification via magnesium precipitation. **F**) Agarose gel analysis of 12-mer chromatin fibers on H2A$^{N110C}$-Atto$_{488}$ containing octamer with increased octamer:DNA ratio. **G**) Native-PAGE analysis of ScaI digested 12×601 chromatin from F). 1×601/MMTV nucleosomes and DNA observed. **H**) Agarose gel analysis of magnesium precipitated 12-mer chromatin fibers. Input (In), pellet (P), supernatant (S). In-gel fluorescence, excitation at $\lambda_{ex}$=488nm (H2A$^{N110C}$-Atto$_{488}$), and GelRed: (DNA intercalating stain).

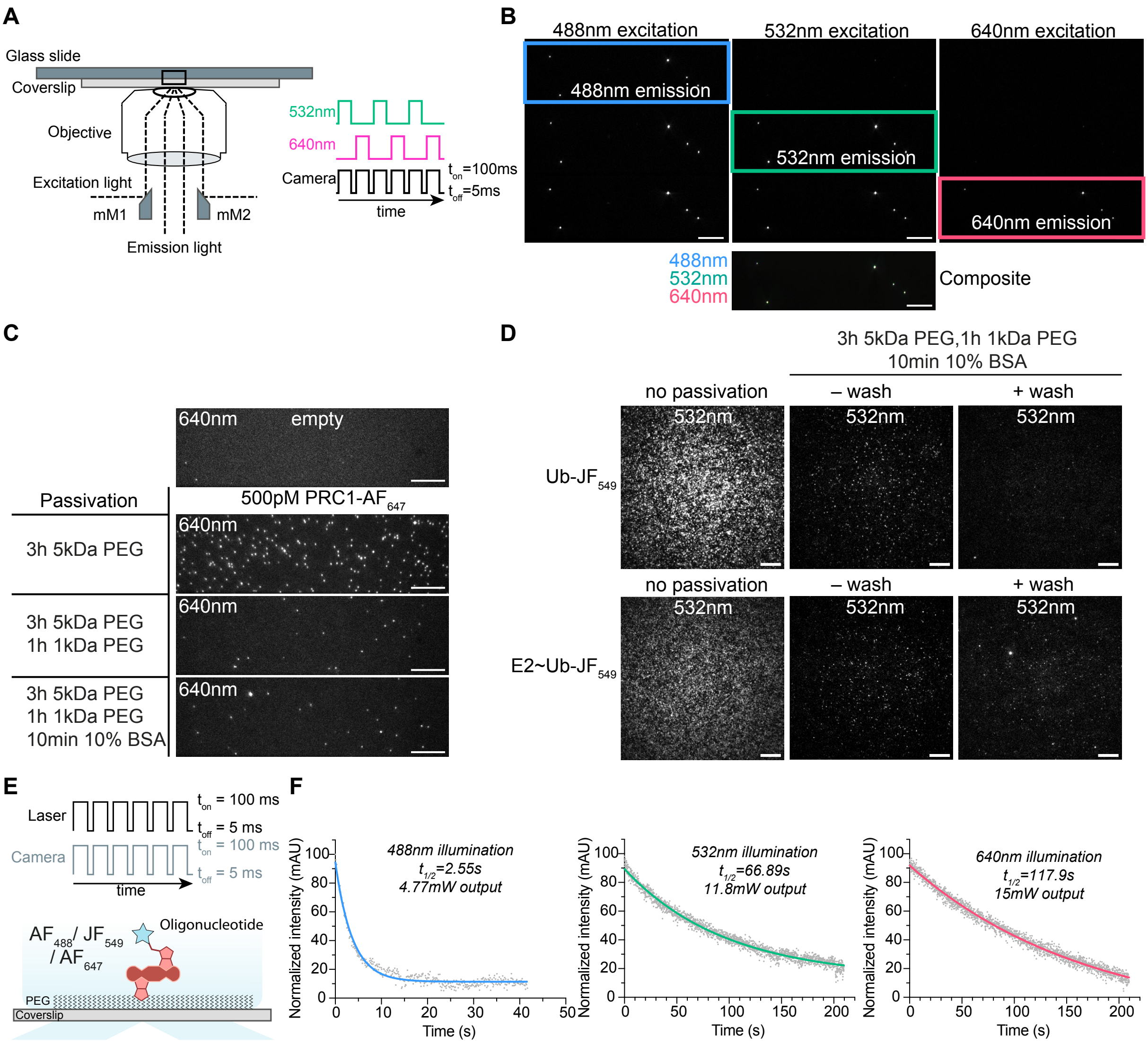


**Fig. S6. Details of single-molecule imaging protocol. A**) Scheme of micro-mirror TIRF (mmTIRF) setup (MadCityLabs). The excitation laser is guided into the objective lens and coupled back out using a pair of micro-mirrors (mM). Emission is collected through the center, without the use of a dichroic mirror thus spatially separating the excitation from the emission. **B)** Slide illumination and alignment with TetraSpeck™ microspheres 0.1µm. Top: full camera view at the specific wavelength, with the selected regions of interest (ROIs). Illumination at 488 nm, 532 nm, 640 nm. Bottom: 3-color composite image after channel alignment. Scale bar: 10µm. **C)** Optimization of flow chamber passivation conditions. Unspecific binding of 500 pM PRC1-$AF_{647}$ injected into flow chambers and imaged after washing with imaging buffer (see Methods for buffer composition). ROIs of 640 nm excitation at different passivation conditions. Optimal passivation is achieved via 3 h treatment of amino-silanized flow channels with 5kDa PEG-bis-Succinimidyl Valerate (SVA), followed by 1 h treatment using 1kDa PEG-SVA, and finally 10 min with 10% BSA. Scale bar: 10µm. **D)** Unspecific binding of 100 pM Ub-$JF_{549}$ and 500 pM E2~Ub-$JF_{549}$ on non-passivated and passivated surface (3 h 5kDa PEG, 1 h 1kDa PEG, 10 min 10% BSA). Wash with imaging buffer supplemented with 20 mM imidazole and 0.025% Tween removes most of the unspecifically bound fluorescent ubiquitin. Scale bar: 10µm. **E)** Scheme of photobleaching experiment with fluorescent oligonucleotides labeled with Alexa Fluor 488 ($AF_{488}$), Janelia Fluor 549 ($JF_{549}$) or Alexa Fluor 647 ($AF_{647}$) immobilized on a coverslip of a microfluidic device. Laser and camera in stroboscopic imaging mode. **F)** Fluorophore photobleaching profiles following a monoexponential decay: experimental half-life of Alexa Fluor

488 ($AF_{488}$) at 488 nm $t_{1/2}$=5.1 s, of Janelia Fluor 549 ($JF_{549}$) at 532 nm $t_{1/2}$=445.9 s, and of Alexa Fluor 647 ($AF_{647}$) at 640 nm $t_{1/2}$=117.9 s.

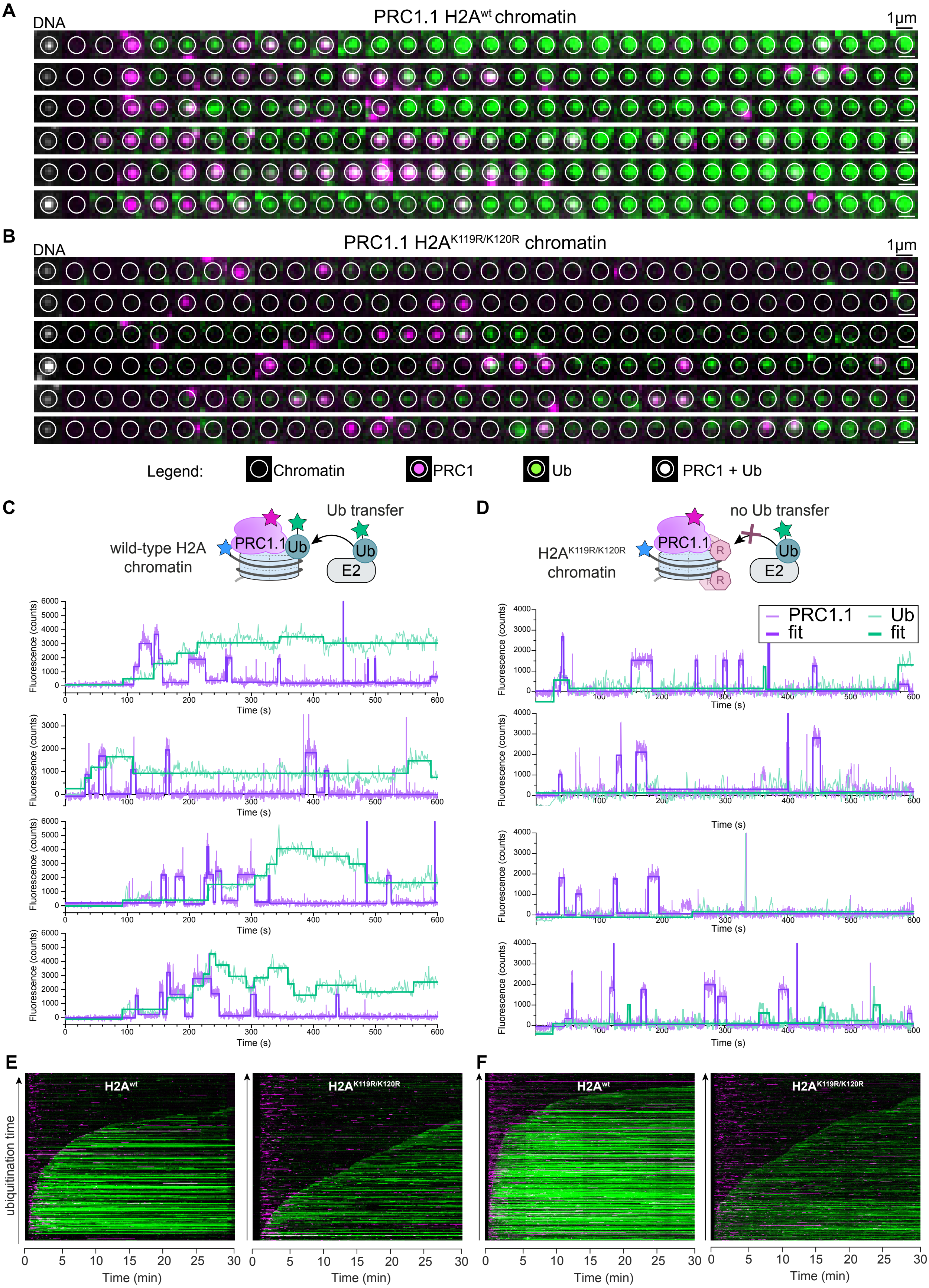


**Fig. S7. PRC1.1 ubiquitylation progress. A**) Extracted fluorescence images from a single 12×601-AF488-biotin chromatin fiber (white circle: position of the chromatin fiber, green: E2~Ub-

$JF_{549}$ / Ub-$JF_{549}$ emission, magenta: PRC1.1-$AF_{647}$ emission). PRC1.1 on wild-type H2A chromatin and **B**) on $H2A^{K119R/K120R}$ mutant chromatin. Scale bar: 1µm. **C**) Top: scheme of ubiquitylation reaction, bottom: single-molecule fluorescence time traces of chromatin ubiquitylation. PRC1.1-$AF_{647}$ on wild-type H2A and **D**) on $H2A^{K119R/K120R}$ mutant 12×601-$AF_{488}$-biotin chromatin. **E**) Example composite plots of PRC1.1 ubiquitylation process over 30 min at [E2~Ub] = 25 nM, [PRC1.1]=500 pM. Each line represents a time trace, fluorescence intensity is encoded in color intensity. n = 206 traces. Left: wild-type H2A, right: $H2A^{K119R/K120R}$. **F**) same as B) but with [E2~Ub] = 40 nM, [PRC1.1]=800 pM.

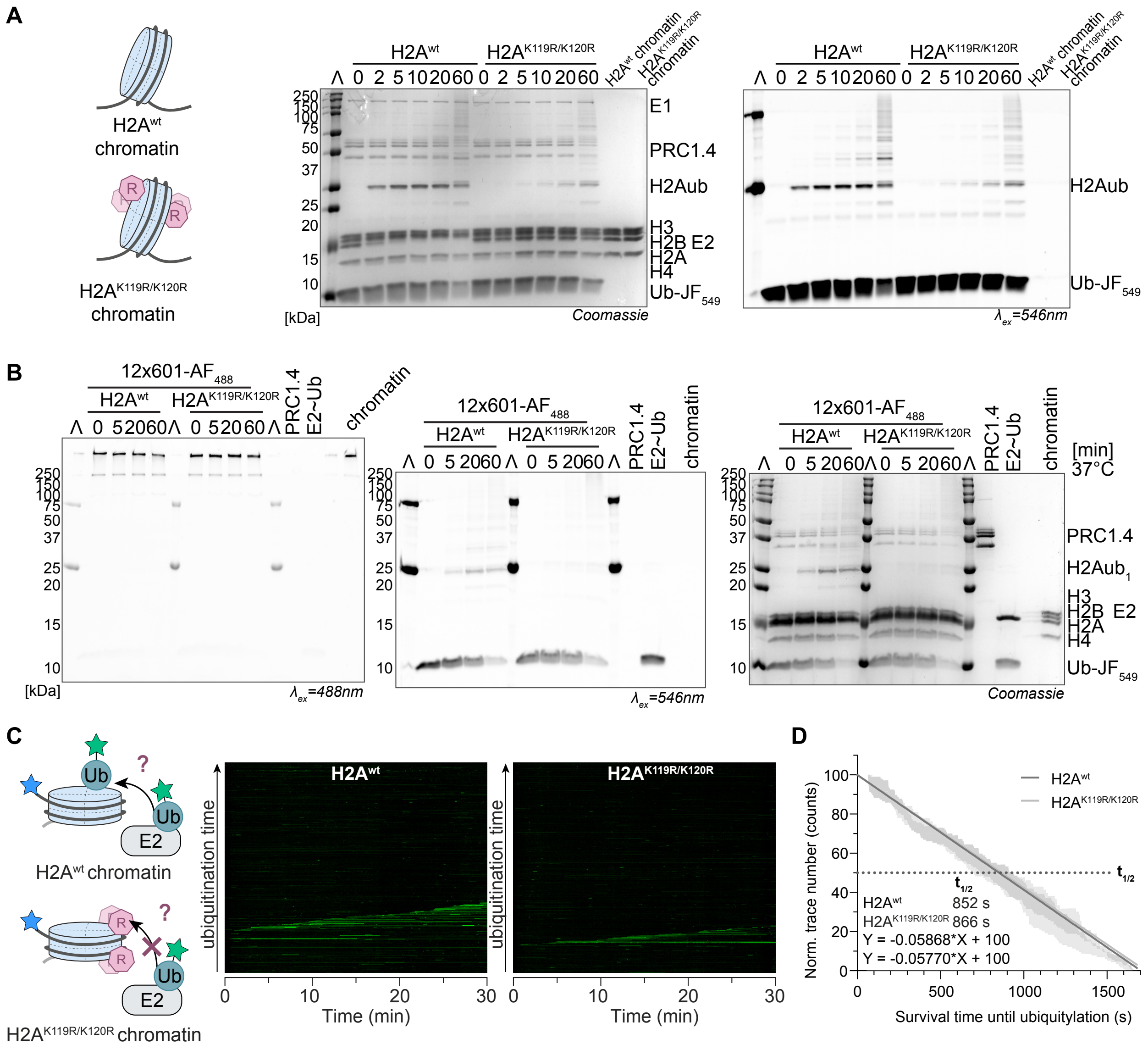

**Fig. S8. Ubiquitylation control assays. A)** SDS-PAGE analysis of the ubiquitylation progress on wild-type H2A and H2A$^{K119R/K120R}$ 12-mer chromatin fibers. Coomassie and in-gel fluorescence, excitation at $\lambda_{ex}$=546nm (Ub-JF$_{549}$). H2A$^{K119R/K120R}$ ubiquitylation is delayed as the specific ubiquitylation sites are blocked (*1*). **B)** Ubiquitylation on biotinylated chromatin. PRC1.4 ubiquitylation assay on H2A$^{wt}$ and H2A$^{K119R/K120R}$ 12×601-biotin-AF$_{488}$ chromatin with E2~Ub, no E1 and ATP. In-gel fluorescence, excitation at $\lambda_{ex}$=488nm (12×601-biotin-AF$_{488}$ DNA) and $\lambda_{ex}$=546nm (Ub-JF$_{549}$) and Coomassie blue. **C)** Composite plots of ubiquitylation process on wild-type H2A and H2A$^{K119R/K120R}$ 12-mer chromatin fibers over 30 min at [E2~Ub] = 25 nM. Each line represents a time trace; fluorescence intensity is encoded in color intensity. **D)** Quantification of the unspecific E2~Ub catalyzed ubiquitylation. Distribution of survival times until ubiquitylation, calculated from **C**), on wild-type chromatin (H2A$^{wt}$, n = 2) and control chromatin (H2A$^{K119R/K120R}$, n = 2). The y-axis is normalized by the number of traces. Data is linear fit (solid line) with ± SEM. Observed half-times $t_{1/2}$ are for H2A$^{wt}$ : 852 s, H2A$^{K119R/K120R}$ : 866 s.

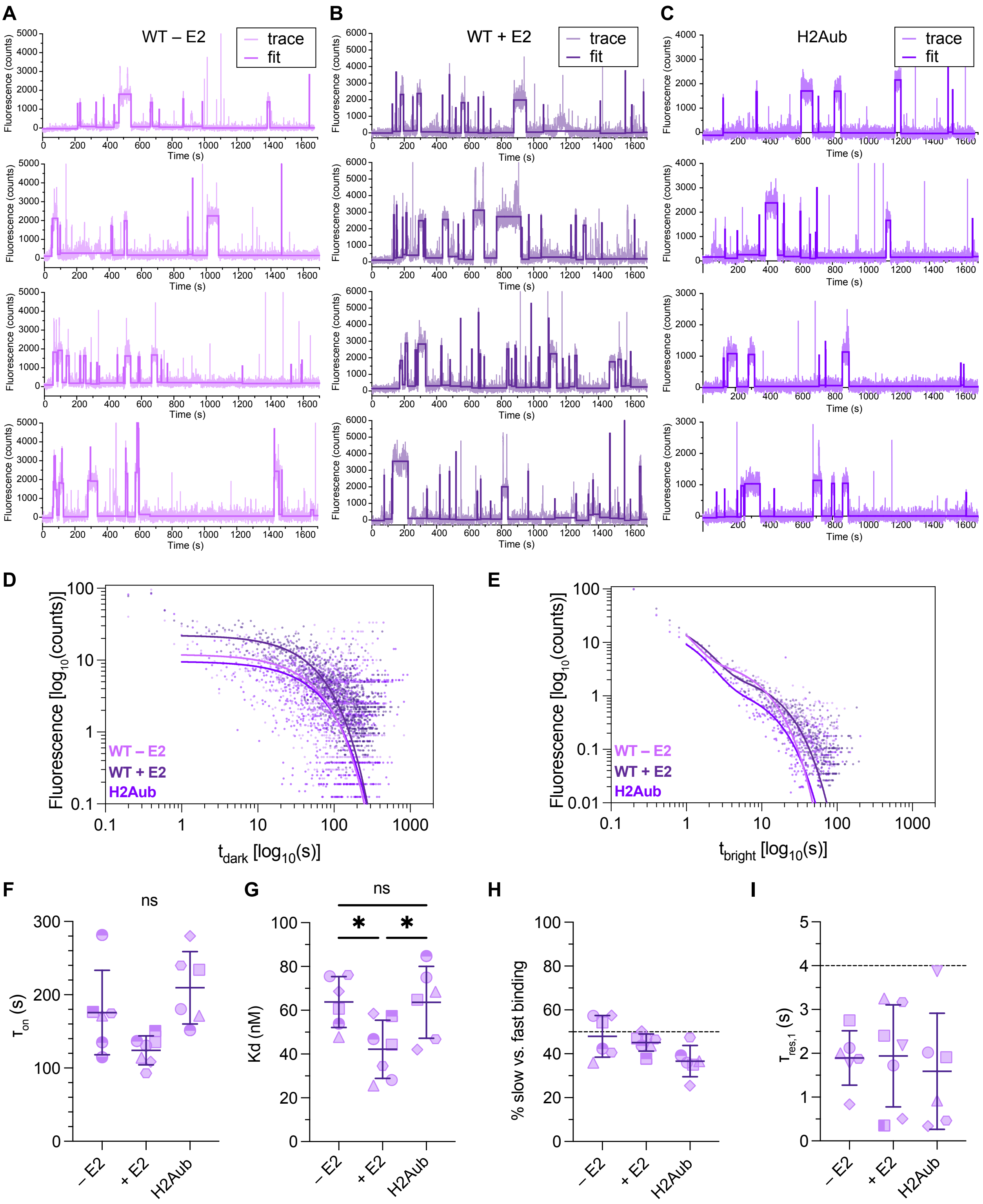


**Fig. S9. Binding dynamics of PRC1.1 to chromatin fibers.** PRC1.1 binding to **A)** unmodified, or 'wild-type', in the absence of E2 (WT – E2) or to **B)** unmodified chromatin in the presence of E2 (WT + E2, 50 nM) or to **C**) H2AK119ub-containing (H2Aub) 12-mer chromatin fibers. The single-molecule fluorescence traces are fitted by a step regression function. Free ($t_{dark}$) and bound times ($t_{bright}$) are determined via thresholding. **D)** Association time histograms fitted by a mono-exponential decay function $f(t) = A * exp(-t/\tau_{on})$ to obtain the $\tau_{on}$ of PRC1.1 binding.

Compare to **Fig. 3**, showing the cumulative association time histograms. **E)** Dissociation time histograms fitted by a two-exponential decay function $f(t) = \sum_{i=1}^{2} A_i * exp(-t/\tau_{res,i})$ to obtain the $\tau_{res}$ of PRC1.1 binding. Compare to **Fig. 3**, showing the cumulative dissociation time histograms. Plotted mean of n = 6 replicates for PRC1.1 on wild-type chromatin in the absence of E2 (WT – E2, pink), n = 7 replicates for PRC1.1 on WT chromatin in the presence of [E2]=50nM (WT + E2, dark violet), n = 6 replicates for PRC1.1 on H2AK119ub chromatin (H2Aub, violet), from independent experiments. **F)** Association time constants $\tau_{on}$; **G)** dissociation rate calculated from the $\tau_{res,2}$ and $\tau_{on}$; **H)** percentage of slow dissociation reactions with $\tau_{res,2}$ over all dissociation events and **I)** dissociation time constants $\tau_{res,1}$ of PRC1.1 binding kinetics on wild-type chromatin in the absence or presence of E2 (–/+ E2) and on H2AK119ub (H2Aub) chromatin. Each symbol represents individual results: n = 6 replicates for PRC1.1 on wild-type chromatin in the absence of E2 (WT – E2), n = 7 replicates for PRC1.1 on WT chromatin in the presence of [E2] = 50 nM (WT + E2), n = 6 replicates for PRC1.1 on H2AK119ub chromatin (H2Aub) from independent experiments. Plotted mean, error bars, s.d.; Statistical testing: one-way ANOVA, Tukey's multiple comparison test (*) $p < 0.05$, ns: non-significant.

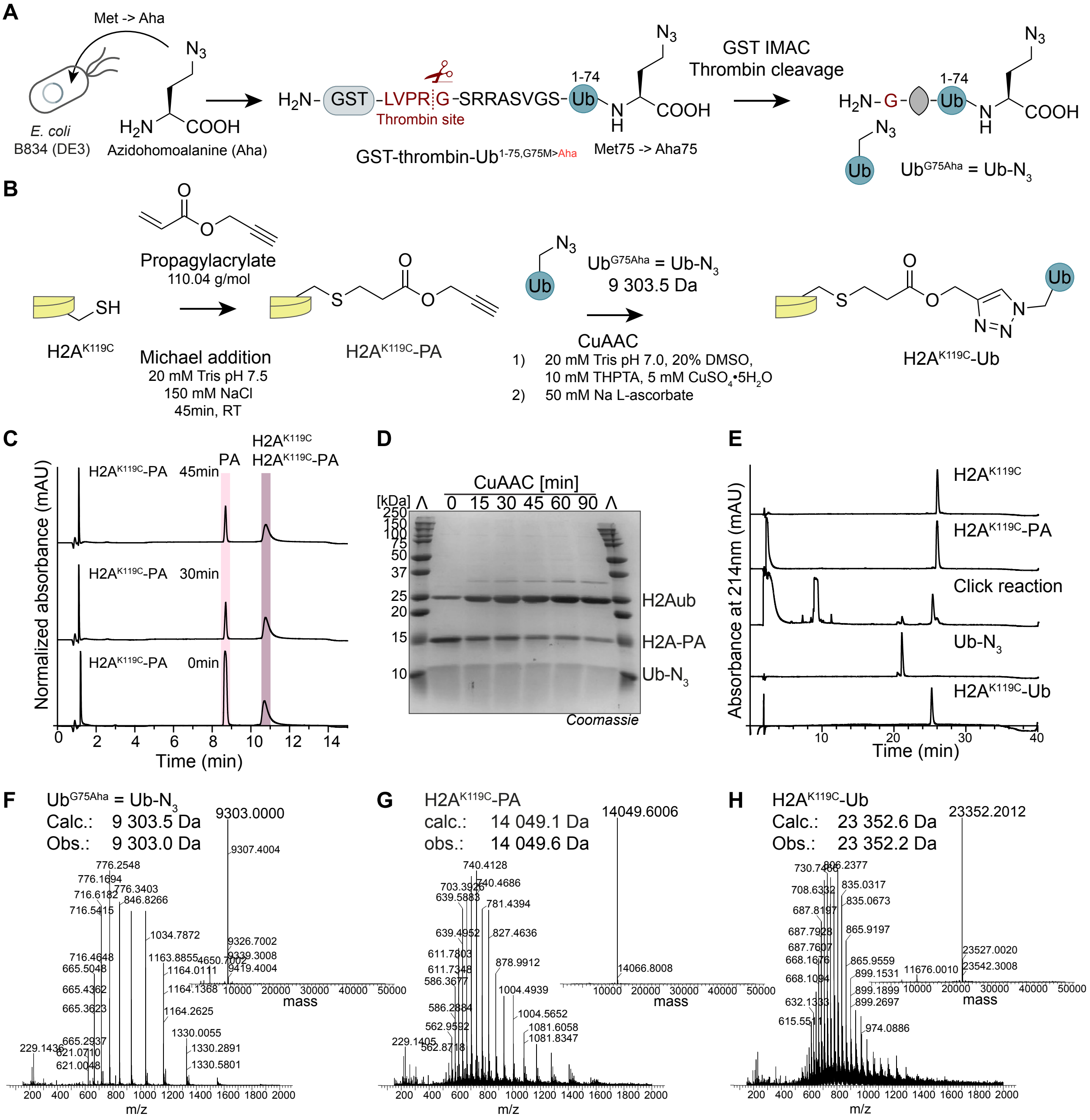


**Fig. S10. Synthesis of H2A$^{K119C}$-Ub. A)** Scheme of GST-thrombin-Ubiquitin$^{1\text{-}75,G75M}$ expression and azidohomoalanine (Aha) incorporation. GST IMAC and thrombin cleavage remove the GST-tag and yield ubiquitin$^{1\text{-}75,G75M}$-Aha (Ub-$N_3$). **B**) Preparation of H2A$^{K119C}$-PA-Ub via Michael's addition of propargyl acrylate (PA) on H2A$^{K119C}$ followed by CuAAC click reaction. **C**) Analytical RP-HPLC chromatogram of H2A$^{K119C}$ modification reaction with PA. Bottom: 0 min, middle: 30 min, top: 45 min of PA with H2A$^{K119C}$ at room temperature (RT). H2A$^{K119C}$ and the product H2A$^{K119C}$-PA co-elute at $t_R$=10.8min. **D**) SDS-PAGE analysis of copper(I)-catalyzed azide-alkyne cycloaddition (CuAAC) reaction monitored over 90 min at RT, Coomassie Blue. **E**) Analytic RP-HPLC analysis to confirm the purity of H2A$^{K119C}$, H2A$^{K119C}$-PA, CuAAC reaction, Ub-$N_3$ and H2A$^{K119C}$-Ub (H2Aub). ESI-MS m/z and deconvolved mass spectra of **F)** Ub-N3 (calc.: 9303.5 Da, obs.: 9303.0 Da), **G**) H2A$^{K119C}$-PA ( calc.:14049.1 Da, obs.: 14049.6 Da) and **H**) H2A$^{K119C}$-PA-Ub (=H2Aub) (calc.: 23353.6 Da, obs.: 23353.2 Da).

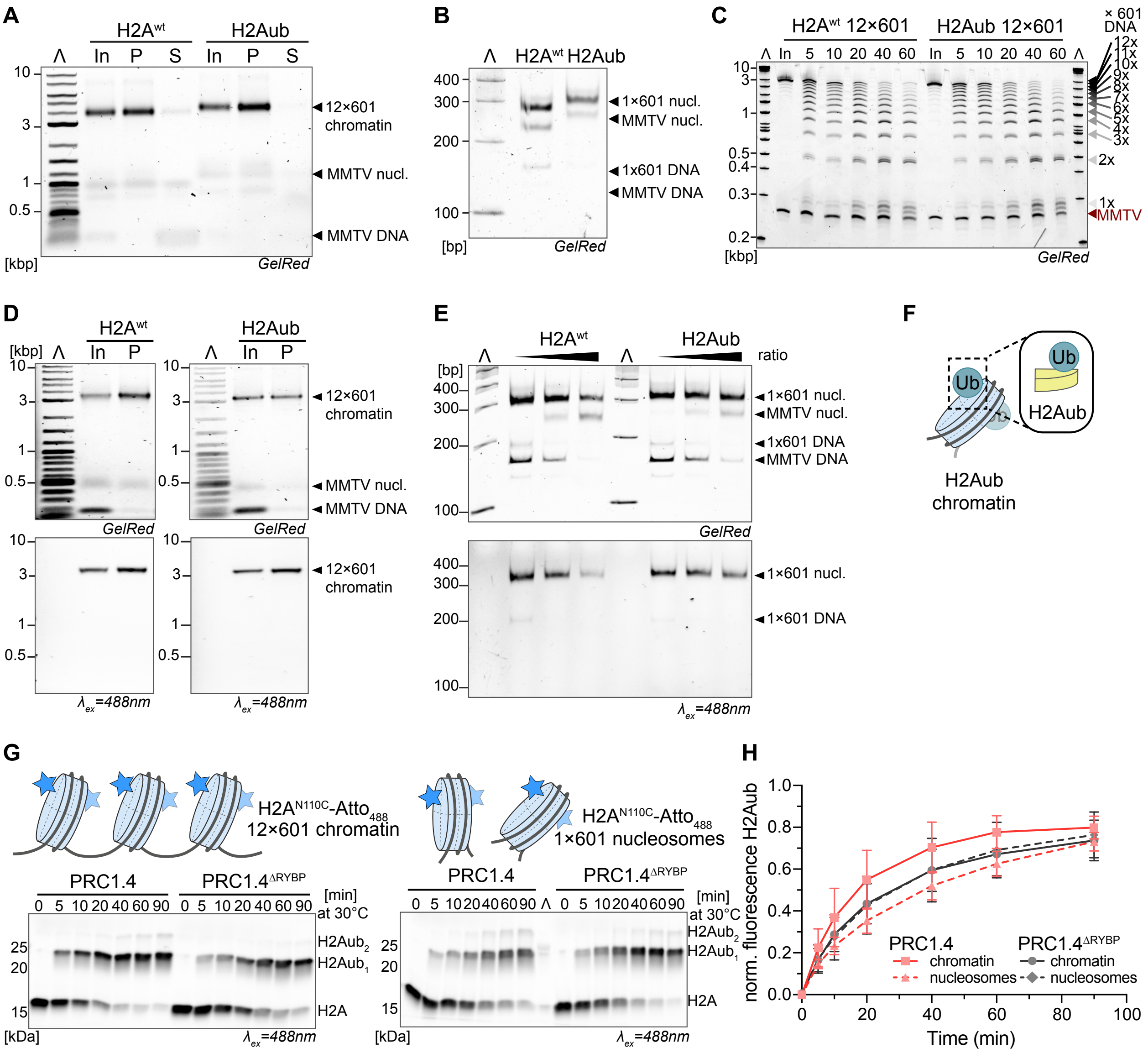


**Fig. S11. H2Aub impact on chromatin conformation and E3 ligase activity. A**) Agarose gel analysis of magnesium precipitated wild-type H2A and H2Aub 12-mer chromatin fibers. Input (In), pellet (P) and supernatant (S) samples, GelRed (DNA intercalating stain). **B**) Native-PAGE analysis of ScaI digested wild-type H2A and H2Aub 12-mer chromatin fibers, 1×601/MMTV buffer nucleosomes and DNA. **C**) Native-PAGE analysis of an MNase digest over time of wild-type (wt) and H2AK119ub (H2Aub) 12×601 chromatin DNA. Analysis of DNA fragments after PCR purification. MMTV and ×601 DNA pieces indicated. Chromatin accessibility is not affected by H2Aub. **D**) same as **A**) but with wild-type H2A and H2Aub 12×601-biotin-$AF_{488}$ fluorescent chromatin fibers. Only input (In) and pellet (P) fractions are shown. GelRed and in-gel fluorescence, excitation at $\lambda_{ex}$=488nm (12×601-biotin-$AF_{488}$). **E**) same as in **B**) but with wild-type H2A and H2Aub 12×601-biotin-$AF_{488}$ fluorescent chromatin fibers. Different ratios are shown. **F)** Scheme of ubiquitylated nucleosomes carrying H2Aub. **G**) Ubiquitylation assay of PRC1.4 and $PRC1.4^{\Delta RYBP}$ on 12-mer chromatin fibers (left) and 1×601 nucleosomes (right). In-gel fluorescence, excitation at $\lambda_{ex}$=488nm ($H2A^{N110C}$-$Atto_{488}$). **H**) Quantification of the H2Aub normalized to the total lane fluorescence. RYBP subunit stimulates PRC1.4 activity on chromatin but not on nucleosomes. Data are mean ± s.d., n = 3 independent experiments.

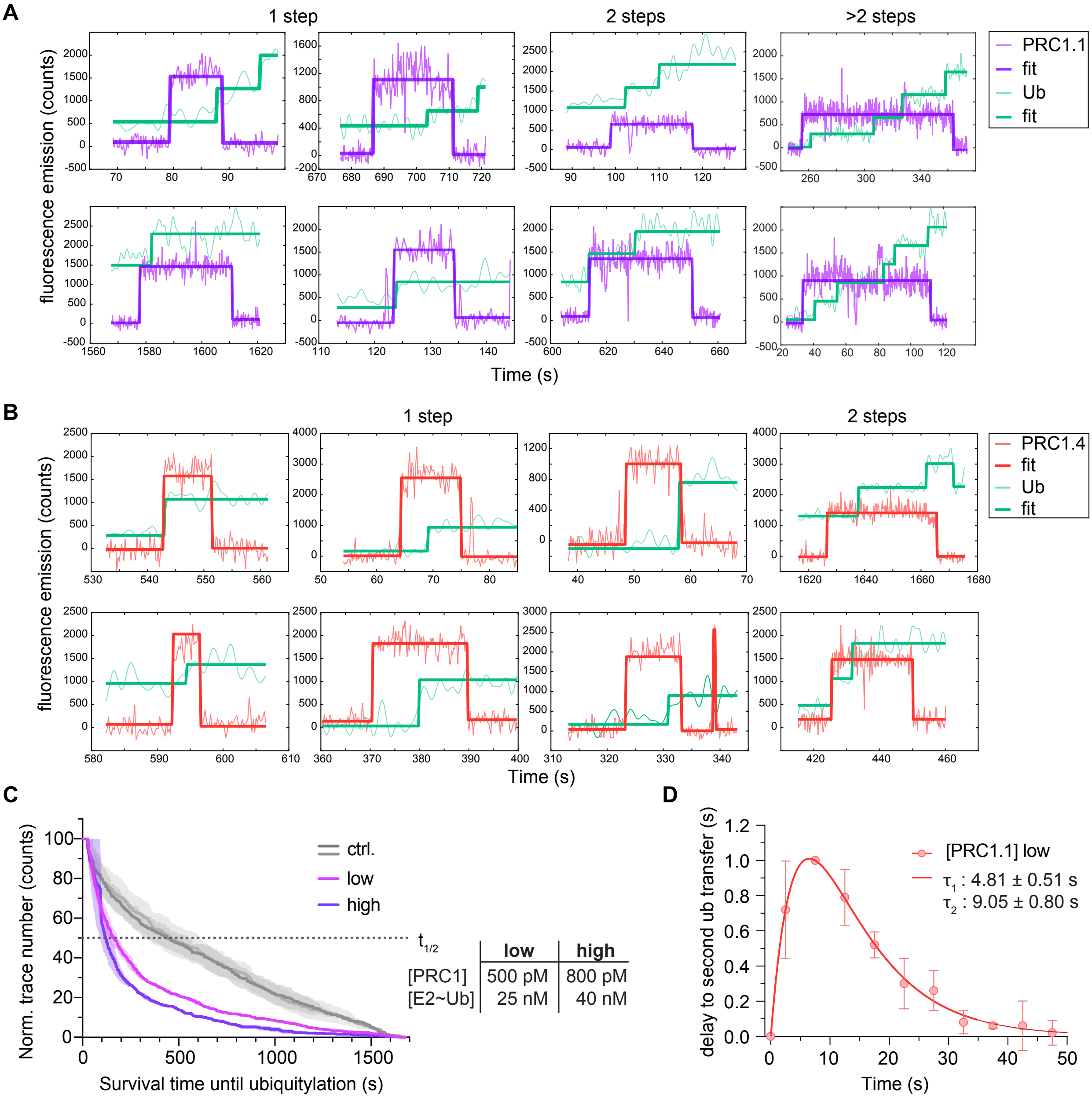


**Fig. S12. Isolated and global ubiquitylation events of PRC1**. **A**) Examples of PRC1.1 binding events on wild-type H2A 12×601-biotin-$AF_{488}$ chromatin fibers associated with one, two or multiple ubiquitin transfer steps. PRC1.1 binding in violet and ubiquitin transfer in green. Traces fitted with linear stepwise regression function. **B**) Examples of PRC1.4 binding events on wild-type H2A 12×601-biotin-$AF_{488}$ chromatin fibers associated with one or two ubiquitin transfer steps. PRC1.4 binding in salmon and ubiquitin transfer in green. Traces fitted with linear stepwise regression function. **C**) Quantification of the E2~Ub catalyzed ubiquitylation. Distribution of survival times until ubiquitylation, calculated from composite plots, on wild-type H2A chromatin ($H2A^{wt}$, low n = 3, high n = 2) and control chromatin ($H2A^{K119R/K120R}$, low n = 3, high n = 2). The y-axis is normalized by the number of traces. Data is mean (solid line) with ± SEM, n=3. Observed half-times $t_{1/2}$ are indicated on the right. **D**) For PRC1.4 binding events associated with more than one ubiquitin transfer steps: Histogram of $\tau_{ub,2}$, i.e. the time distribution between first and second ubiquitin transfer, at 0.5 nM PRC1.4, 25 nM E2~Ub. Solid line: bi-exponential fit, yielding time constants $\tau_1$ = 4.81 ± 0.51 s for the first process, and $\tau_2$ = 9.05 ± 0.80 s for the second process.

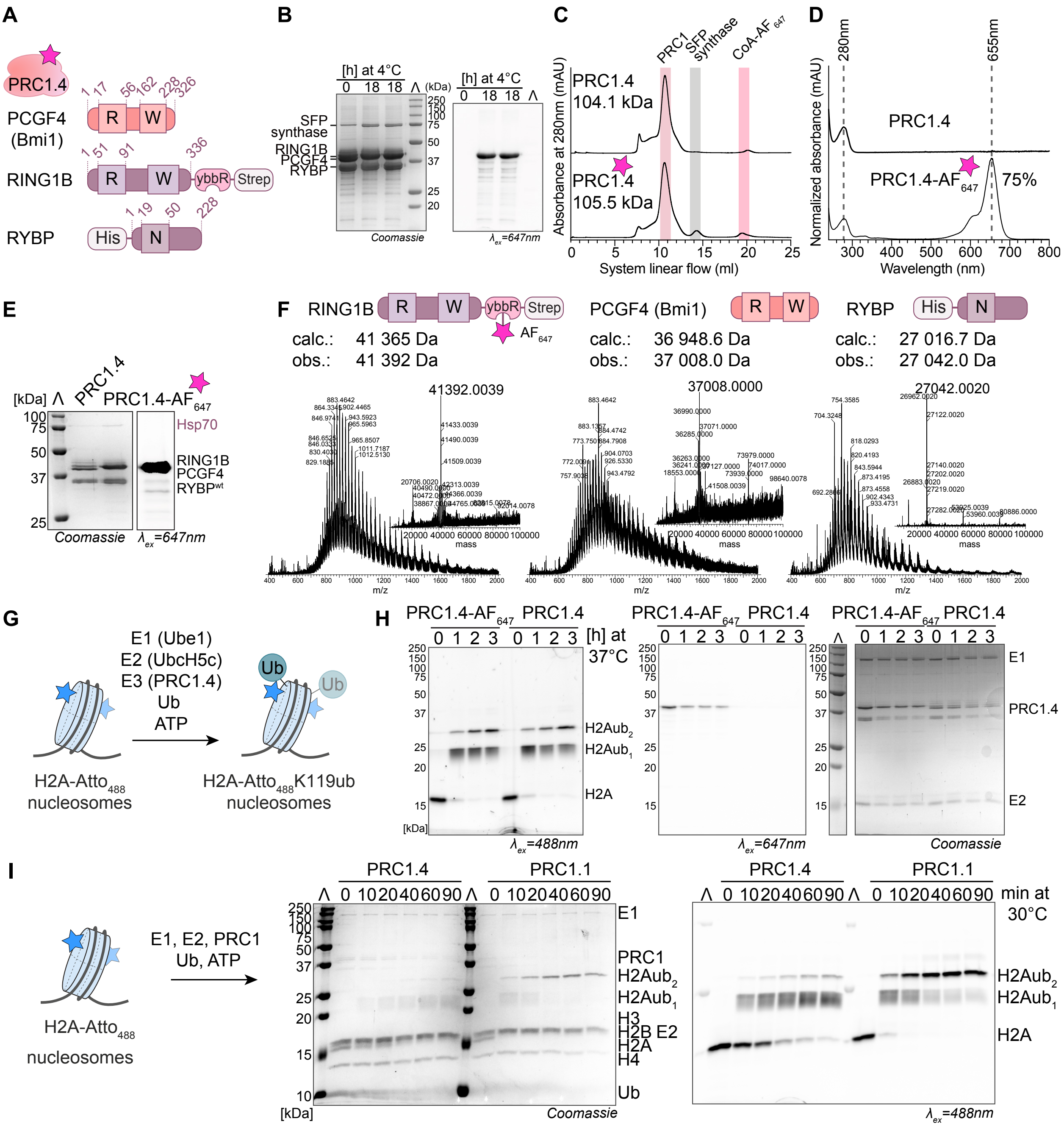


**Fig. S13. Purification of PRC1.4. A**) PRC1.4 complex subunits and domains. ybbR-, His- and Strep-tag are indicated. Numbers correspond to amino acid residues. R: RING domain W: RAWUL domain. **B**) SDS-PAGE analysis of ybbR-tag labeling reaction of PRC1.4 with Alexa Fluor 647 ($AF_{647}$) C2 maleimide detected via in-gel fluorescence, excitation at $\lambda_{ex}$=647nm (PRC1.4-$AF_{647}$), and Coomassie blue. **C**) SEC profiles, **D**) UV-Vis absorbance spectra, and **E**) SDS-PAGE analysis (Coomassie and in-gel fluorescence $\lambda_{ex}$ = 647 nm) of unlabeled PRC1.4 and labeled PRC1.4-$AF_{647}$. **F**) MS spectra of the PRC1.4-$AF_{647}$ complex subunits. ESI-MS m/z and deconvolved mass spectra of: RING1B-ybbR-$AF_{647}$-Strep (calc. 41365 Da, obs. 41392 Da), PCGF4 (Bmi1) (calc. 36948.6 Da, obs. 37008.0 Da) and His-RYBP (calc. 27016.7 Da, obs. 27042.0 Da). Calculated mass (calc.), observed mass (obs.). **G**) Ubiquitylation reaction scheme with indicated components, in particular using Atto 488 ($Atto_{488}$) labeled H2A for detection. **H**) Ubiquitylation reactions using labeled PRC1.4-$AF_{647}$ and unlabeled PRC1.4, detected via in-gel fluorescence, excitation at $\lambda_{ex}$=488nm (H2A-$Atto_{488}$) and $\lambda_{ex}$=647nm (PRC1.4-$AF_{647}$), and Coomassie blue.

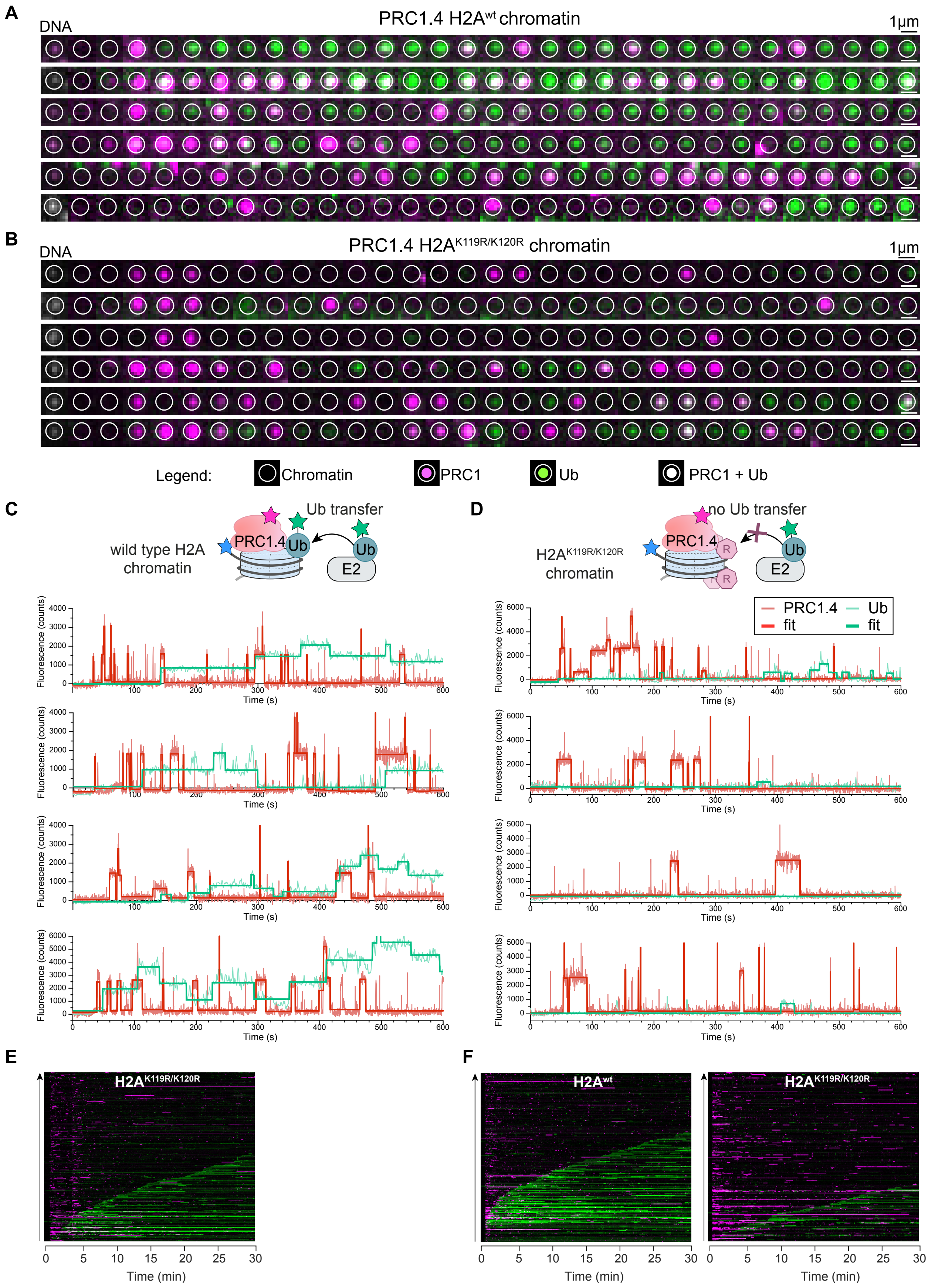
A
DNA
PRC1.4 H2A$^{wt}$ chromatin
1µm
B
DNA
PRC1.4 H2A$^{K119R/K120R}$ chromatin
1µm
Legend:
Chromatin
PRC1
Ub
PRC1 + Ub
C
Ub transfer
wild type H2A chromatin
PRC1.4
Ub
Ub
E2
D
no Ub transfer
H2A$^{K119R/K120R}$ chromatin
PRC1.4
R
R
Ub
E2
PRC1.4
fit
Ub
fit
Fluorescence (counts)
Time (s)
E
H2A$^{K119R/K120R}$
Time (min)
F
H2A$^{wt}$
H2A$^{K119R/K120R}$
Time (min)

**Fig. S14. PRC1.4 ubiquitylation progress. A**) Extracted fluorescence images from a single 12×601-$AF_{488}$-biotin chromatin fiber (white circle: position of the chromatin fiber, green: E2~Ub-$JF_{549}$ / Ub-$JF_{549}$ emission, magenta: PRC1.4-$AF_{647}$ emission). PRC1.4 on wild-type H2A chromatin and **B**) on $H2A^{K119R/K120R}$ mutant chromatin. Scale bar: 1μm. **C**) Top: scheme of ubiquitylation reaction, bottom: single-molecule fluorescence time traces of chromatin ubiquitylation. PRC1.4-$AF_{647}$ on wild-type H2A and **D**) on $H2A^{K119R/K120R}$ mutant 12×601-$AF_{488}$-biotin chromatin. **E**) Example composite plots of PRC1.4 ubiquitylation process over 30 min at [E2~Ub] = 25 nM, [PRC1.4]=500 pM. Each line represents a time trace, fluorescence intensity is encoded in color intensity. Control $H2A^{K119R/K120R}$, n = 261 traces to Fig. 5E wild-type H2A. **F**) same as in E) but with n = 198 traces. Left: wild-type H2A, right: $H2A^{K119R/K120R}$.

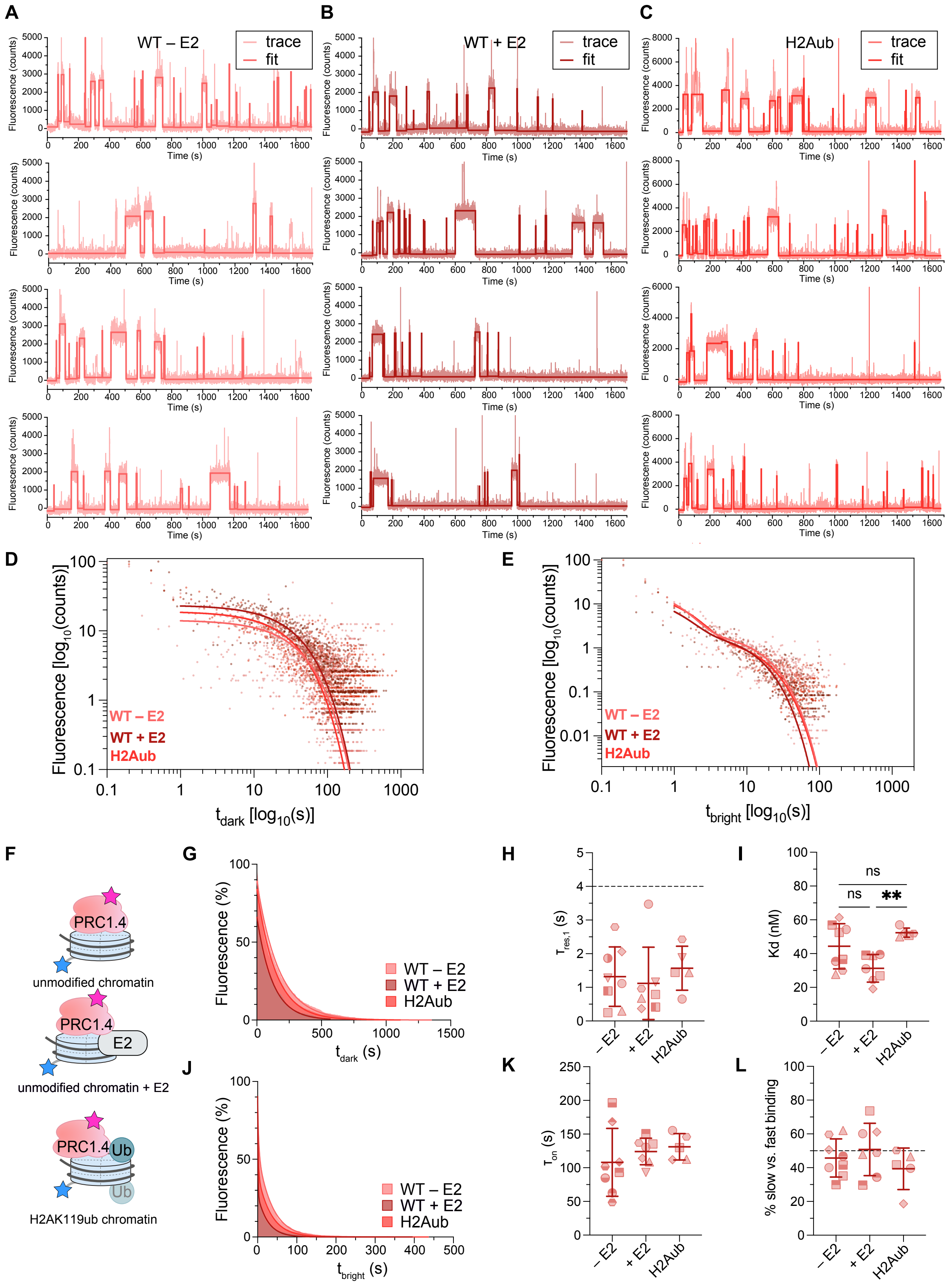


**Fig. S15. Binding dynamics of PRC1.4 to chromatin fibers.** PRC1.4 binding to **A)** unmodified, or ‘wild-type’, in the absence of E2 (WT – E2) or to **B)** unmodified chromatin in the presence of

E2 (WT + E2, 50 nM) or to **C**) H2AK119ub-containing (H2Aub) 12-mer chromatin fibers. The single-molecule fluorescence traces are fitted by a step regression function. Free ($t_{dark}$) and bound times ($t_{bright}$) are determined via thresholding. **D**) Association time histograms fitted by a mono-exponential $f(t) = A * exp(-t/\tau_{on})$ decay function to obtain the $\tau_{on}$ of PRC1.4 binding. (see panel J) for the cumulative histogram). **E**) Dissociation time histograms fitted by a two-exponential decay function $f(t) = \sum_{i=1}^{2} A_i * exp(-t/\tau_{res,i})$ to obtain the $\tau_{res}$ of PRC1.4 binding (see panel G) for the cumulative histogram). Plotted mean of n = 8 replicates for PRC1.4 on wild-type chromatin in absence of E2 (WT – E2, salmon), n = 7 replicates for PRC1.4 on WT chromatin in the presence of [E2] = 50 nM (WT + E2, dark red), n = 5 replicates for PRC1.4 on H2AK119ub chromatin (H2Aub, red) (from independent experiments). **F**) Scheme of binding experiment of PRC1.4 to indicated chromatin fibers, either unmodified or containing H2AK119ub, in the absence or presence of E2 (-E2 or +E2, 50 nM). **G**) Cumulative histogram of $t_{dark}$ times, indicating PRC1.4 binding to chromatin under indicated conditions, fitted by mono-exponential functions (solid lines). **H)** Dissociation time constants $\tau_{res,1}$ and **I**) dissociation rate calculated from the $\tau_{res,2}$ and $\tau_{on}$ of PRC1.4 binding under indicated conditions. **J**) Cumulative histogram of $t_{bright}$ times, indicating PRC1.4 dissociation from chromatin under indicated conditions, fitted by bi-exponential functions (solid lines). **K**) Association time constants $\tau_{on}$ and **L**) the percentage of slow dissociation reactions with $\tau_{res,2}$ over all dissociation events for PRC1.4 binding to chromatin under indicated conditions. Each symbol represents individual results: n = 8 replicates for PRC1.4 on wild-type chromatin in absence of E2 (WT – E2, salmon), n = 7 replicates for PRC1.4 on WT chromatin in the presence of [E2] = 50 nM (WT + E2, dark red), n = 5 replicates for PRC1.4 on H2AK119ub chromatin (H2Aub, red) (from independent experiments). Plotted mean, error bars, s.d.; Statistical testing: one-way ANOVA, Tukey's multiple comparison test (*) $p < 0.05$, (**) $p < 0.01$, ns: non-significant.

**Table S1. PRC1.1 binding kinetics.**

| Exp. # | [PRC1] [pM] | $\tau_{res,1}$ [s] | $\tau_{res,2}$ [s] | $K_{d,2}$ [nM] | $A_1/A_{tot}$ % fast | $A_2/A_{tot}$ % slow | $\tau_{on}$ [s] | molecular $k_{on}$ x $10^5$ [$M^{-1}s^{-1}$] |
|---|---|---|---|---|---|---|---|---|
| **PRC1.1 on wtChromatin** | | | | | | | | |
| 1 | 1000 | 2,002 | 21,46 | 75,55 | 43 | 57 | 135,1 | 6,17 |
| 2 | 1500 | 2,751 | 34,01 | 60,71 | 46 | 54 | 114,7 | 4,84 |
| 3 | 1100 | 2,124 | 33,97 | 68,62 | 43 | 57 | 176,6 | 4,29 |
| 4 | 800 | 1,780 | 22,03 | 76,13 | 60 | 41 | 174,7 | 5,96 |
| 5 | 1000 | 1,874 | 43,04 | 47,82 | 64 | 36 | 171,5 | 4,86 |
| 6 | 800 | 0,836 | 50,17 | 53,90 | 58 | 42 | 281,7 | 3,70 |
| | **Average** | **1,895** | **34,11** | **63,79** | **52** | **48** | **175,7** | **4,97** |
| | **s.d.** | **0,567** | **10,36** | **10,62** | **9** | **9** | **52,6** | **0,87** |
| **PRC1.1 on wtChromatin + E2** | | | | | | | | |
| 1 | 1000 | 3,253 | 55,75 | 28,09 | 53 | 47 | 130,5 | 6,39 |
| 2 | 1100 | 2,400 | 40,35 | 44,26 | 53 | 47 | 135,3 | 5,60 |
| 3 | 1100 | 1,723 | 24,54 | 58,47 | 50 | 50 | 108,7 | 6,97 |
| 4 | 1100 | 2,182 | 35,56 | 34,52 | 53 | 47 | 93,0 | 8,15 |
| 5 | 1100 | 3,168 | 58,83 | 25,60 | 56 | 44 | 114,1 | 6,64 |
| 6 | 800 | 0,504 | 28,02 | 46,87 | 56 | 44 | 136,8 | 7,62 |
| 7 | 800 | 0,352 | 25,21 | 57,39 | 62 | 38 | 150,7 | 6,91 |
| | **Average** | **1,940** | **38,32** | **42,17** | **55** | **45** | **124,2** | **6,90** |
| | **s.d.** | **1,078** | **13,12** | **12,28** | **4** | **4** | **18,3** | **0,76** |
| **PRC1.1 on H2Aub Chromatin** | | | | | | | | |
| 1 | 1000 | 0,925 | 28,87 | 74,98 | 64 | 36 | 180,4 | 4,62 |
| 2 | 1000 | 1,912 | 43,34 | 64,82 | 65 | 35 | 234,1 | 3,56 |
| 3 | 1000 | 2,020 | 79,99 | 42,04 | 75 | 25 | 280,2 | 2,97 |
| 4 | 1000 | 3,874 | 61,46 | 46,82 | 53 | 47 | 239,8 | 3,48 |
| 5 | 1000 | 0,463 | 30,06 | 68,42 | 63 | 37 | 171,4 | 4,86 |
| 6 | 1000 | 0,340 | 21,46 | 84,77 | 61 | 39 | 151,6 | 5,50 |
| | **Average** | **1,589** | **44,20** | **63,64** | **63** | **37** | **209,6** | **4,17** |
| | **s.d.** | **1,210** | **20,54** | **15,00** | **6** | **6** | **45,0** | **0,89** |

**Table S2. PRC1.4 binding kinetics.**

| Exp. # | [PRC1] [pM] | $\tau_{res,1}$ [s] | $\tau_{res,2}$ [s] | $K_{d,2}$ [nM] | $A_1/A_{tot}$ % fast | $A_2/A_{tot}$ % slow | $\tau_{on}$ [s] | molecular $k_{on}$ x $10^5$ [$M^{-1}s^{-1}$] |
|---|---|---|---|---|---|---|---|---|
| **PRC1.4 on wtChromatin** | | | | | | | | |
| 1 | 2000 | 0,293 | 27,75 | 54,28 | 60 | 40 | 62,8 | 6,64 |
| 2 | 1000 | 0,248 | 19,35 | 52,58 | 65 | 35 | 84,8 | 9,83 |
| 3 | 1000 | 1,113 | 18,22 | 61,40 | 49 | 51 | 93,2 | 8,94 |
| 4 | 500 | 1,275 | 39,61 | 29,75 | 40 | 60 | 196,4 | 8,49 |
| 5 | 500 | 2,795 | 36,53 | 27,68 | 38 | 62 | 168,5 | 9,90 |
| 6 | 1000 | 2,061 | 36,69 | 35,49 | 53 | 47 | 108,5 | 7,68 |
| 7 | 2000 | 0,892 | 20,52 | 56,98 | 70 | 30 | 48,7 | 8,55 |
| 8 | 1000 | 1,865 | 33,38 | 36,52 | 58 | 42 | 101,6 | 8,20 |
| | **Average** | **1,318** | **29,01** | **44,33** | **54** | **46** | **108,1** | **8,53** |
| | **s.d.** | **0,826** | **8,14** | **12,49** | **11** | **11** | **47,2** | **1,01** |
| **PRC1.4 on wtChromatin + E2** | | | | | | | | |
| 1 | 500 | 0,677 | 23,70 | 31,04 | 64 | 36 | 122,6 | 13,59 |
| 2 | 500 | 0,788 | 27,17 | 26,88 | 38 | 62 | 121,7 | 13,69 |
| 3 | 500 | 3,000 | 49,32 | 19,12 | 40 | 60 | 157,2 | 10,60 |
| 4 | 1000 | 2,643 | 42,79 | 24,48 | 56 | 44 | 87,3 | 9,55 |
| 5 | 1000 | 1,728 | 32,53 | 38,88 | 56 | 44 | 105,4 | 7,91 |
| 6 | 800 | 0,728 | 27,58 | 39,54 | 72 | 28 | 113,6 | 9,17 |
| 7 | 1000 | 0,929 | 26,11 | 39,10 | 70 | 30 | 85,1 | 9,80 |
| | **Average** | **1,499** | **32,74** | **31,29** | **57** | **43** | **113,3** | **10,62** |
| | **s.d.** | **0,904** | **8,94** | **7,57** | **13** | **13** | **22,7** | **2,06** |
| **PRC1.4 on H2Aub Chromatin** | | | | | | | | |
| 1 | 1000 | 1,439 | 32,71 | 57,01 | 60 | 40 | 155,4 | 5,36 |
| 2 | 1000 | 1,443 | 30,65 | 50,82 | 58 | 42 | 129,8 | 6,42 |
| 3 | 1000 | 0,650 | 25,69 | 52,27 | 81 | 19 | 111,9 | 7,45 |
| 4 | 1000 | 1,883 | 33,75 | 51,91 | 50 | 50 | 146,0 | 5,71 |
| 5 | 1000 | 2,426 | 37,87 | 35,71 | 53 | 47 | 112,7 | 7,39 |
| | **Average** | **1,568** | **32,13** | **49,54** | **61** | **39** | **131,2** | **6,47** |
| | **s.d.** | **0,585** | **3,99** | **7,24** | **11** | **11** | **17,4** | **0,85** |

**Table S3. PRC1 ubiquitylation kinetics.**

| Exp # | Used PRC1 [pM] | Tau_res,act [s] | tau_ub [s] | half-time [s] | # ub transferred [#] |
|---|---|---|---|---|---|
| **PRC1.1** | | | | | |
| 1 | 500 | 16,2 | 16,3 | 204 | 1,60 |
| 2 | 500 | 20,21 | 8,222 | 119 | 1,86 |
| 3 | 500 | 21,41 | 9,007 | 149 | 1,78 |
| | **tau from fit** | **19,54** | **10,38** | **157,33** | **1,75** |
| | **s.d.** | **0,4** | **0,54** | **43,11** | **0,13** |
| 4 | 800 | 20,74 | 7,13 | 145 | 2,06 |
| 5 | 800 | 21,38 | 8,356 | 88 | 1,91 |
| | **tau from fit** | **21,07** | **7,76** | **116,5** | **1,98** |
| | **s.d.** | **0,19** | **0,29** | **40,31** | **0,11** |
| **PRC1.4** | | | | | |
| 1 | 500 | 21,03 | 5,743 | 349 | 1,80 |
| 2 | 500 | 20,54 | 10,08 | 122 | 2,02 |
| 3 | 500 | 20,07 | 11,24 | 222 | 1,65 |
| | **tau from fit** | **20,59** | **8,89** | **231** | **1,82** |
| | **s.d.** | **0,32** | **0,48** | **113,77** | **0,19** |
